# An ER-resident lipid scramblase is crucial for biogenesis and function of apicomplexan parasite secretory organelles

**DOI:** 10.1101/2024.11.14.623710

**Authors:** G. Srinivas Reddy, Somesh M. Gorde, Kanika Saxena, Manash Kumar Behera, Abhijit S Deshmukh, Puran Singh Sijwali

**Affiliations:** CSIR-Centre for Cellular and Molecular Biology, Hyderabad-500007, India; Academy of Scientific and Innovative Research (AcSIR), Ghaziabad-201002, India; National Institute of Animal Biotechnology (NIAB), Hyderabad-500032, India

**Keywords:** Vacuole membrane protein 1, VMP1, *Toxoplasma*, *Plasmodium*, Apicomplexa, Micronemes, Rhoptries, Dense granules, Lipid droplets, DedA domain, Scramblase

## Abstract

Apicomplexan parasites, including *Plasmodium falciparum* and *Toxoplasma gondii*, contain specialized secretory organelles such as micronemes, rhoptries, and dense granules, which are essential for parasite motility, host cell invasion, development, and egress. DedA superfamily proteins are implicated in lipid mobilization, which is a key requirement for organelle biogenesis. Herein, we identified and investigated the vacuole membrane protein 1 (VMP1), a DedA superfamily member, of *P. falciparum* (PfVMP1) and *T. gondii* (TgVMP1). PfVMP1 and TgVMP1 are ER-localized lipid scramblases. TgVMP1 depletion adversely affected parasite development, motility, host cell invasion, and egress. These phenotypes were consistent with impaired rhoptry and dense granule biogenesis, and decreased secretion of micronemes, rhoptries, and dense granules in TgVMP1-depleted parasites, indicating a crucial role for TgVMP1 in the biogenesis and function of these organelles. TgVMP1 depletion impaired lipid droplet homeostasis and ER organization. Restoration of the ER-localized lipid scramblase by complementing TgVMP1-depleted parasites with PfVMP1 or a homolog as distant as human VMP1 rescued the depleted parasites, indicating their functional conservation and a crucial role for ER-resident lipid scramblase activity in the biogenesis and function of secretory organelles. The essentiality of TgVMP1 for parasite development and likely functional conservation of apicomplexan VMP1 proteins highlight their drug-target potential.

## Introduction

Protozoans of the phylum Apicomplexa are intracellular obligate parasites, which cause diseases in humans, livestock, and wild animals ^1^. *Plasmodium* and *Toxoplasma* are the most studied apicomplexans, causing malaria and toxoplasmosis, respectively. The central feature of apicomplexans is apical complex, consisting of the apical polar ring at the apex of the cell and specialized secretory organelles called rhoptries and micronemes ^2^. Another specialized secretory organelle of apicomplexans is dense granules, which are present throughout the cytoplasm. Microneme and rhoptry secretions are essential for parasite motility, invasion, formation of the parasitophorous vacuole (PV) around the parasite, and egress ^2–9^. Dense granule secretion contributes to the formation of intravacuolar network (IVN) within the PV, host cell remodeling, nutrient import, and export of various substances across the parasitophorous vacuole membrane (PVM) ^6,9,10^. As these organelles are unique and essential, proteins present in their secretions are being pursued as drug and vaccine targets. The biogenesis of micronemes, rhoptries, and dense granules is far from clear, nonetheless, ER and Golgi have been shown to be the sites of biogenesis of the vesicles and proteins destined to these organelles ^11^. Lipid scramblases, which move lipids between the membrane leaflets, have been shown to be crucial for various cellular processes, including membrane/organelle biogenesis, lipid distribution, and regulation of ER-organelle contacts ^12^. DedA (downstream (of hisT) DNA gene A) superfamily proteins are implicated in lipid distribution and organelle biogenesis, and has members in all the life forms ^13,14^. The vacuole membrane protein 1 (VMP1) is the best studied DedA superfamily member, and DedA superfamily proteins are yet to be reported in parasitic protozoans. This prompted us to identify and investigate the DedA superfamily proteins in apicomplexan parasites.

VMP1 was first identified as an overexpressed gene in the rat model of experimental acute pancreatitis ^15^. All the characterized VMP1 homologs are ER-localized multi-pass transmembrane proteins, and share a stretch of amino acids that is called DedA domain, a characteristic feature of the DedA superfamily ^13,16^. VMP1 has roles in a variety of cellular processes, including the formation of autophagosomes, regulation of the ER–organelle contact sites, lipid transport and distribution, secretion of lipoproteins, and formation of viral replication organelles ^17–27^.

A major function of VMP1 is in the formation of autophagosomes at an early stage. The mammalian VMP1 has been shown to recruit the PI3K complex (VPS34/Beclin 1/Vps15) at the phagophore assembly site via interaction with Beclin 1, which then generates PI3P to initiate phagophore formation ^28,29^. The cellular abnormalities observed due to the loss of VMP1 in mammalian cells are also mirrored in VMP1 knock-out unicellular eukaryotes *Dictyostelium discoideum* and *Chlamydomonas reinhardtii*, which exhibited fragmented ER, distorted Golgi, impaired endocytosis and exocytosis, damaged mitochondria, and accumulation of ubiquitinated-protein aggregates and lipid droplets (LDs) ^30–32^. VMP1 has been shown to activate the sarcoplasmic/endoplasmic reticulum Ca^2+^-ATPase (SERCA), which transports cytosolic Ca^2+^ into ER ^33^, thereby, inducing the dissociation of ER–membrane contacts. Consistently, depletion of *Homo sapiens* VMP1 (HsVMP1) increased the number of ER contacts with autophagosomes, mitochondria, LDs, and endolysosomes ^33,34^.

Mammalian VMP1 and the transmembrane protein 41B (TMEM41B), a DedA superfamily protein as well, have recently been shown to function as lipid scramblases, and implicated in maintaining lipid asymmetry across the ER leaflets and at the ER contact sites with the membranes of other organelles ^23,35^. VMP1 and TMEM41B have been shown to be required for the formation of virus-induced replication organelles in contact with the ER, and it has been proposed that their interaction with viral proteins and mobilization/distribution of lipids at the ER-virus replication organelle site could be the underlying mechanism ^26,27,36,37^. VMP1 has been shown to be key for the formation of LDs and secretion of lipoproteins, which likely depend on its scramblase activity ^24,34^.

The pleiotropic functions of VMP1 are plausible based on its scramblase activity and involvement in several ER-associated processes, as ER is the hub of lipid synthesis, organelle biogenesis, and establishes dynamic physical contacts with various organelles and structures^38^. In this study, we identified a VMP1 homolog in *T. gondii* (TgVMP1) and *P. falciparum* (PfVMP1), and investigated whether it has a role in the biogenesis and function of micronemes, rhoptries, and dense granules. We show that TgVMP1 and PfVMP1 are ER-localized lipid scramblases. *T. gondii* VMP1 is crucial for the biogenesis and function of secretory organelles, maintenance of ER organization, and LD homeostasis.

## Results

### Apicomplexan parasites contain DedA superfamily members

The phylum Apicomplexa contains obligate intracellular parasites, including *Plasmodium*, *Toxoplasma*, *Cryptosporidium*, *Babesia*, *Eimeria*, *Theileria* and *Neospora*, which cause diseases in humans, livestock and wild animals. To identify the apicomplexan DedA superfamily members, we searched the genome databases of *P. falciparum* and *T. gondii* (https://veupathdb.org/veupathdb/app) by BLAST using the amino acid sequences of DedA superfamily proteins of human (HsVMP1), *S. cerevisiae* (ScTVP38), and *Escherichia coli* (YohD, YdjZ, YqaA) as queries. The BLAST searches predicted multiple DedA-domain containing proteins in *T. gondii* (TGME49_244370, TGME49_279370, TGME49_248870, and TGME49_301350) and *P. falciparum* (PF3D7_1474600, PF3D7_1365700, and PF3D7_0611000). Of these predicted proteins, TGME49_244370 and PF3D7_1474600 showed higher sequence identity (29.3-30.3%) with HsVMP1 than the remaining proteins (10.2-20.1%), and we termed TGME49_244370 and PF3D7_1474600 as the *T. gondii* VMP1 (TgVMP1) and *P. falciparum* VMP1 (PfVMP1), respectively. Since VMP1 homologs have crucial roles in several processes, including organelle biogenesis, we focussed on TgVMP1 and PfVMP1. TgVMP1 and PfVMP1 homologs are also present in several apicomplexans, which share 28.1-32.6% sequence identity with HsVMP1, and 34.5-83.3% sequence identity among themselves (**Table S1**). Apicomplexan VMP1 homologs, including TgVMP1 and PfVMP1, contain the characteristic multi-transmembrane domain architecture and DedA domain of the DedA superfamily (**Figure S1A**). DedA domain is characterized by [F/Y]XXX[R/K] and GXXX[V/I/L/M]XXXX[F/Y] sequence motifs, and two predicted re-entrant loops (**Figure S1B**) ^13^. These motifs are “YXXXY” and “GXXXMXXXXF” in PfVMP1 and TgVMP1. The “Gly” residue in the GXXX[V/I/L/M]XXXX[F/Y] motif has been shown to be essential for the function of *D. discoideum* VMP1 ^34^, and is also conserved in TgVMP1 and PfVMP1. The AlphaFold structures of PfVMP1 and TgVMP1 also overlapped with that of HsVMP1 (RMSD of 3.736 Å between HsVMP1 and PfVMP1, and 3.362 Å between HsVMP1 and TgVMP1) (**Figure S1C**) ^39,40^. The conserved DedA superfamily domain architecture and structural similarity of PfVMP1 and TgVMP1 with HsVMP1 indicate that these belong to the DedA superfamily.

### PfVMP1 and TgVMP1 localize to ER

Towards characterization of apicomplexan VMP1 proteins, we replaced wild-type PfVMP1 and TgVMP1 coding sequences in *P. falciparum* D10 and *T. gondii* RHTIR1 strains with the coding sequences of PfVMP1GFP and TgVMP1AID_HA_, respectively. Cloned recombinant *P. falciparum* (PfVMP1GFP_KI_) and *T. gondii* (RHTIR1-VMP1AID) parasites showed desired modification of the respective target locus (**Figure S2**). PfVMP1GFP_KI_ parasites showed expression of PfVMP1GFP throughout the asexual erythrocytic stages (**Figures 1A** and **1B**), which co-localized with ER-Tracker (**Figure 1C**), indicating that PfVMP1 is an ER-resident protein.

**Figure 1.**
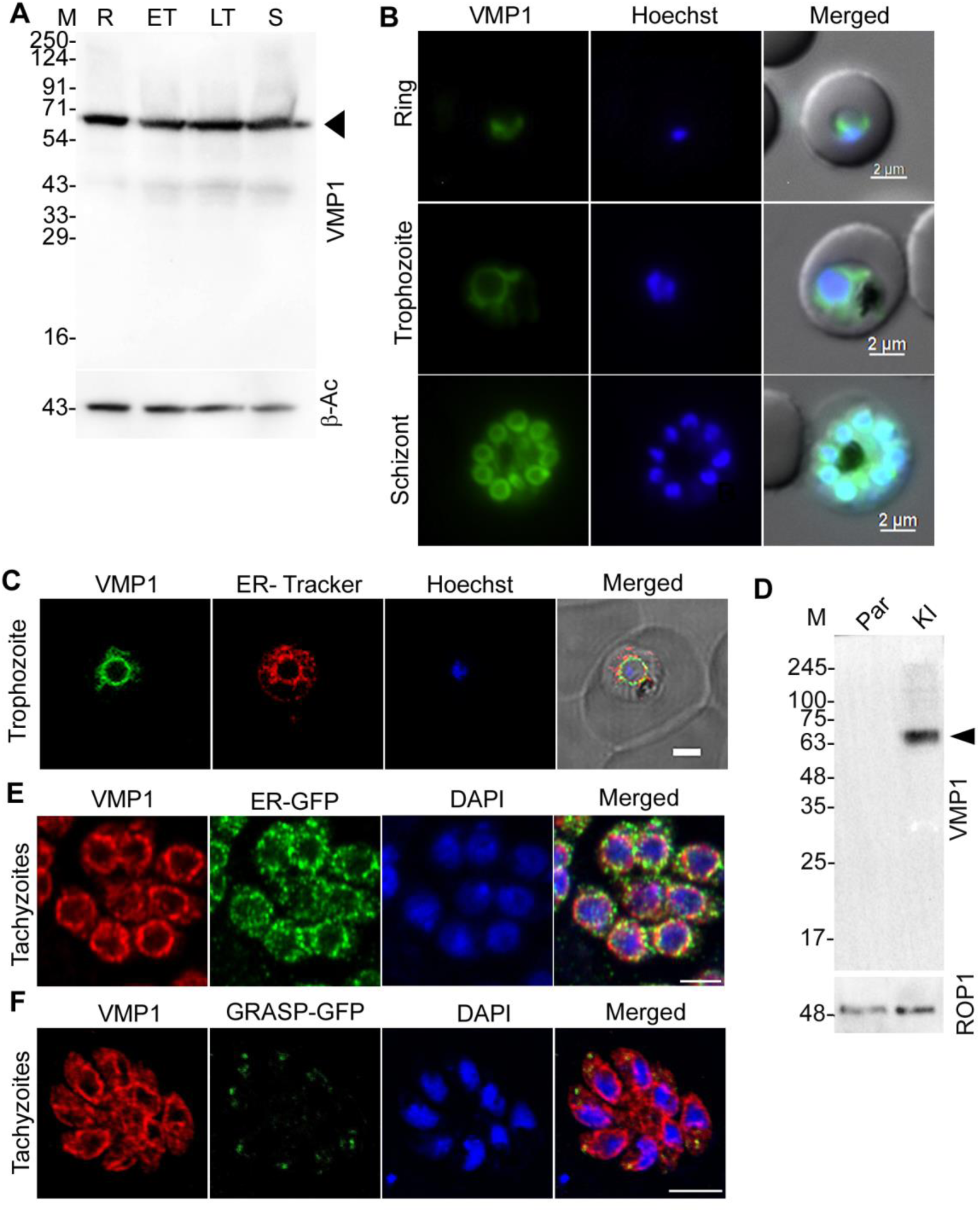
PfVMP1 and TgVMP1 localize to ER. **A.** Total lysates of synchronized PfVMP1GFPKI parasites corresponding to the ring (R), early trophozoite (ET), late trophozoite (LT) and schizont (S) stages were processed for western blotting using antibodies to GFP (VMP1) and β-actin (β-Ac) as a loading control. The arrowhead indicates PfVMP1GFP band (∼73 kDa), and protein size markers (M) are in kDa. **B.** The indicated stages of PfVMP1GFPKI parasites were evaluated for localization of PfVMP1GFP by live cell fluorescence microscopy. The panels are for PfVMP1GFP (VMP1), nucleus (Hoechst), and overlap of these two panels with bright field panel (Merged). Scale bar is shown in the merged panel. **C**. PfVMP1GFPKI trophozoite stage were evaluated for localization of PfVMP1GFP and ER-Tracker by live cell fluorescence confocal microscopy. The images show signal for PfVMP1GFP (VMP1), ER-Tracker, nucleus (Hoechst), and overlap of these three images with bright field image (Merged). The merged image shows co-localization of VMP1 with ER-Tracker (Pearson’s correlation coefficient = 0.643±0.09; n = 30 images). Scale bar is 2 µm. **D.** The total lysates of parental RHTIR1 strain (Par) and RHTIR1-VMP1AID (KI) parasites were processed for western blotting using antibodies to HA (VMP1) and rhoptry protein 1 (ROP1) as a loading control. The arrowhead indicates TgVMP1AIDHA (∼76.3 kDa), and protein size markers (M) are in kDa. **E.** The ER reporter (ER-GFP)-expressing RHTIR1-VMP1AID tachyzoites were evaluated for localization of TgVMP1AIDHA and ER-GFP by IFA. The confocal images show signal for TgVMP1AIDHA (VMP1), ER-GFP, nuclear stain (DAPI), and overlap of the three images (Merged). The merged image shows co-localization of TgVMP1AIDHA with ER-GFP (Pearson’s correlation coefficient = 0.68±0.06; n = 43 images). Scale bar is 5 µm. **F.** The Golgi-reporter (GRASP-GFP)-expressing RHTIR1-VMP1AID tachyzoites were evaluated for localization of TgVMP1AIDHA and GRASP-GFP by IFA. The confocal images show signals for TgVMP1AIDHA (VMP1), GRASP-GFP, nuclear stain (DAPI), and overlap of the three images (Merged). The merged image shows co-localization of TgVMP1 with GRASP-GFP (Pearson’s correlation coefficient = 0.57±0.08; n = 35 images). Scale bar is 5 µm.

RHTIR1-VMP1AID parasites would express the AID receptor TIR1 and TgVMP1 with C-terminal mini-AID-3×HA (TgVMP1AID_HA_). mAID or mAID-fusion protein undergoes proteasomal degradation in the presence, but not in the absence, of auxin/indole-3-acetic acid (IAA) in the cells expressing TIR1 ^41^. RHTIR1-VMP1AID tachyzoites expressed TgVMP1AID_HA_ (**Figure 1D**). To check the subcellular localization of TgVMP1, we designed a GFP-based ER-reporter (ER-GFP) that contained BiP signal sequence in the beginning and ER-retention signal (HDEL) at the end of GFP. We also used the *P. falciparum* Golgi reassembly stacking protein (GRASP)-GFP as a Golgi-reporter (GRASP-GFP) ^42,43^. We episomally expressed ER-GFP or GRASP-GFP in RHTIR1-VMP1AID parasites, which showed co-localization of these reporters with TgVMP1AID_HA_ (**Figures 1E** and **1F**), indicating that TgVMP1 is present both in ER and Golgi.

### TgVMP1 is essential for parasite development

We used two knock-down approaches for PfVMP1. In the first approach, wild-type PfVMP1 coding region was replaced with PfVMP1-cDDHA-*glmS* ribozyme sequence in *P. falciparum* D10 strain to obtain a trimethoprim- and glucosamine-inducible conditional knock-down line (PfVMP1dKD), which would express PfVMP1cDD_HA_ fusion protein (**Figure S3A)**. In the second approach, wild-type PfVMP1 coding region was replaced with PfVMP1-mAID_HA_ sequence in the *P. falciparum* 3D7 strain to generate an auxin-inducible knock-down line (PfVMP1AID), which would express PfVMP1AID_HA_ fusion protein (**Figure S4A)**. PfVMP1dKD and PfVMP1AID parasites expressed the respective fusion proteins, but neither showed any noticeable depletion of the fusion protein nor any growth defect under knock-down conditions (**Figures S3** and **S4**), which might be due to the intrinsic nature and membrane localization of PfVMP1. Hence, we could not investigate PfVMP1 further.

On the other hand, RHTIR1-VMP1AID parasites showed efficient depletion of TgVMP1AID_HA_ in the presence of IAA (+IAA), with a near complete depletion in 3 hrs, whereas the levels of control proteins (SAG1 and ROP1) remained unchanged (Figure 2A), confirming IAA-inducible depletion of TgVMP1AID_HA_. Consistently, TgVMP1AID_HA_ signal, but not SAG1 signal, was almost absent in the IFA images of +IAA RHTIR1-VMP1AID parasites, whereas both the signals were prominent in -IAA parasites (Figure 2B).

**Figure 2.**
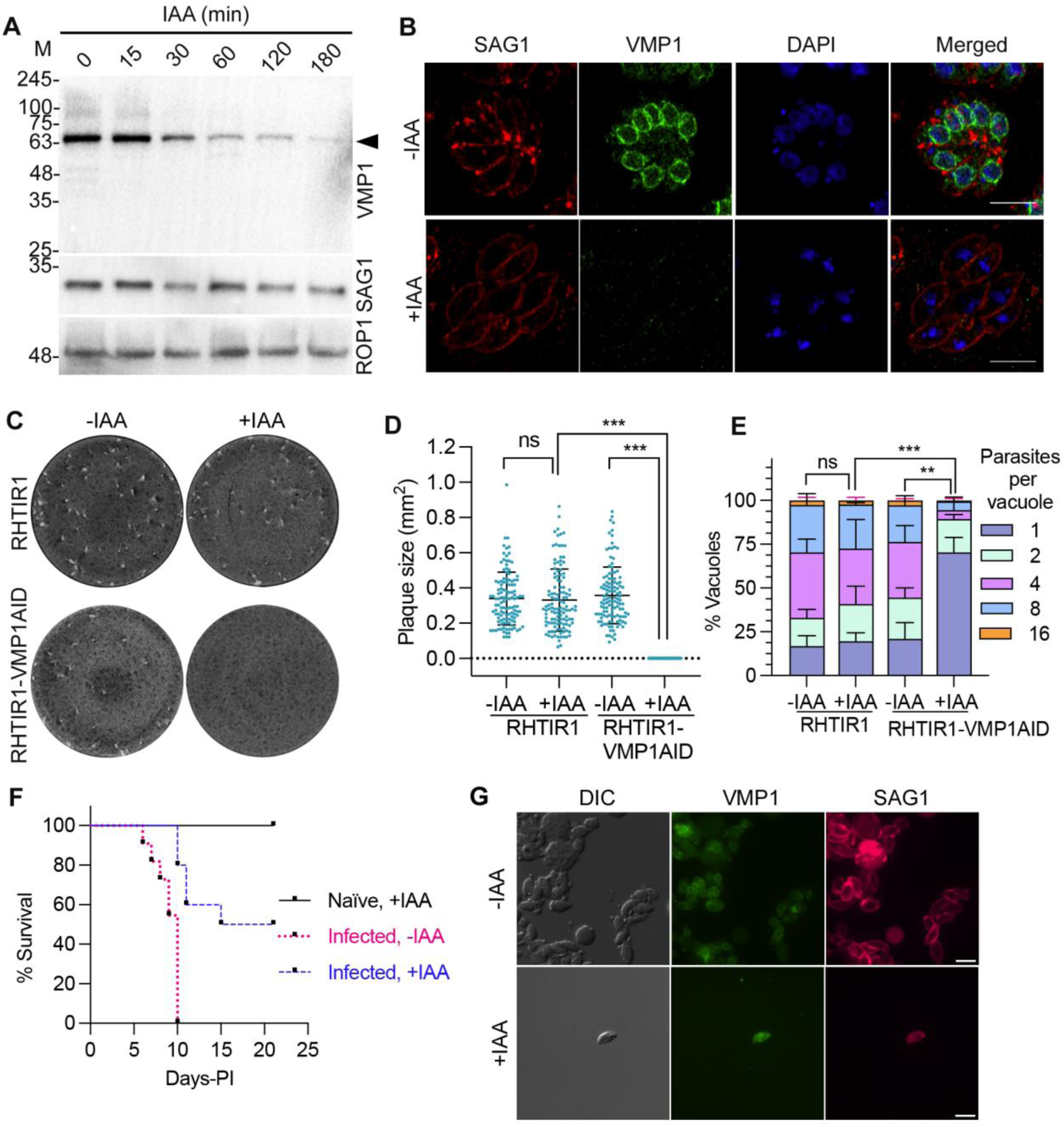
TgVMP1 is essential for parasite development. RHTIR1-VMP1AID tachyzoites were grown in vitro or in mice with (+IAA) or without (-IAA) IAA, and compared for growth. **A.** RHTIR1-VMP1AID parasites were cultured with IAA, parasites were collected at the indicated time points, and lysates were evaluated for TgVMP1AIDHA, SAG1 and ROP1 proteins by western blotting. The blot shows IAA-inducible depletion of TgVMP1AIDHA (VMP1) with time, whereas SAG1 and ROP1 levels remain unchanged. The arrowhead indicates TgVMP1AIDHA, and protein size markers (M) are in kDa. **B.** RHTIR1-VMP1AID parasites were cultured without (-IAA) or with (+IAA) IAA, and evaluated for the localization of TgVMP1AIDHA and SAG1 proteins by IFA. The Airyscan images show signal for SAG1, TgVMP1AIDHA (VMP1), nuclear stain (DAPI), and overlap of all the three images (Merged). Scale bar = 10 µm. Note that VMP1 signal is nearly absent in +IAA parasites as compared with –IAA parasites. **C.** RHTIR1 and RHTIR1-VMP1AID parasites were cultured in vitro for 8 days with (+IAA) or without (-IAA) IAA, stained with Giemsa, and observed for plaques. The white areas represent plaques, which are absent in +IAA RHTIR1-VMP1AID parasites. **D.** The plaque sizes (mm^2^) in “C” were measured, and shown as mean plaque size (Plaque size (mm^2^)) on y-axis for the indicated parasites on x-axis. Each data is mean with SD error bar based on at least 100 plaques from 3 independent experiments. **E.** RHTIR1 and RHTIR1-VMP1AID tachyzoites were cultured in vitro in the presence (+IAA) or absence (-IAA) of IAA, stained for the PV marker GRA1, and the number of parasites/PV was counted. The number of PVs containing different number of parasites was plotted as a percentage of the total number of PVs (% Vacuoles) on y-axis for the indicated parasites on x-axis. Each data is mean with SD error bar based on at least 100 PVs from each of 3 independent experiments. The significance of difference is for the PVs containing a single parasite only. **F.** Two groups of mice (Infected, n = 10 mice/group) were infected with RHTIR1-VMP1AID parasites, and one group of mice (Naïve, n = 5) was used as a control. The infected mice were maintained with (Infected, +IAA) or without (Infected, -IAA) IAA, the naïve mice were given IAA (Naïve, +IAA), and all the mice were monitored for 21 days. The plot shows percent survival of mice (% Survival) on y-axis over days post-infection (Days-PI) on x-axis. **G**. Peritoneal exudates of infected +IAA and –IAA mice that succumbed to infection were visualized by IFA using antibodies to SAG1 and VMP1 proteins. Shown are the representative images for bright field (DIC), TgVMP1AIDHA (VMP1), and SAG1 (scale bar = 5 µm). Very few parasites were observed in the exudate of +IAA mice as compared with that of –IAA mice. The significance of difference between two groups is indicated with p-value (** = P < 0.01, *** = P < 0.001, ns: not significant).

We next assessed the effect of TgVMP1AID_HA_ depletion on parasite development. *T. gondii* tachyzoites invade host cells, replicate every 6-8 hrs for about 2-3 days to produce 64-128 cells within the PV, and finally egress to infect the neighboring uninfected host cells ^44^. Repeated rounds of lytic cycles cause lysis of infected host cells, leaving clear areas in between the uninfected cells, which are called plaques. RHTIR1-VMP1AID and parental RHTIR1 parasites were cultured in the presence (+IAA) or absence (-IAA) of IAA for 8 days, and assessed for plaques. +IAA or -IAA RHTIR1 parasites and -IAA RHTIR1-VMP1AID parasites formed plaques of similar size, indicating similar growth (Figures 2C and **2D**). On the other hand, +IAA RHTIR1-VMP1AID parasites did not form plaques (Figures 2C and **2D**), indicating that TgVMP1 is essential for overall parasite development. To assess intracellular development, RHTIR1 and RHTIR1-VMP1AID tachyzoites were cultured for 24 hrs with (+IAA) or without (-IAA) IAA, and the number of parasites/PV were counted. The majority of +IAA RHTIR1-VMP1AID parasites did not progress beyond 1 parasite/PV stage, whereas the majority of -IAA RHTIR1-VMP1AID parasites and ±IAA RHTIR1 parasites developed to 4-8 parasites/PV stage (Figure 2E), further supporting an essential role of TgVMP1 in intracellular tachyzoite development.

We next evaluated the effect of TgVMP1 depletion on in vivo development of RHTIR1-VMP1AID tachyzoites. BALB/c mice were infected with RHTIR1-VMP1AID tachyzoites, maintained with (+IAA) or without (-IAA) IAA, and monitored for 21 days. As a control, a group of naïve mice was also maintained with IAA (+IAA). All the infected -IAA mice succumbed to infection by day 10, whereas only 50% of the infected +IAA mice succumbed to infection by day 15 (Figure 2F). The peritoneal exudates of dead +IAA mice had few parasites as compared with several parasites in the peritoneal exudates of dead –IAA mice (Figure 2G). The peritoneal exudates of infected +IAA mice that survived for 21 days did not show parasites. The +IAA naïve mice survived and did not exhibit any visible symptoms, indicating that IAA was not toxic at the concentration used. The delayed death and 50% survival of infected +IAA mice indicate that TgVMP1 is critical for tachyzoite development in vivo.

### TgVMP1 is crucial for gliding motility, invasion, and egress

To understand how TgVMP1 depletion affected parasite development, we next evaluated TgVMP1-depleted parasites for gliding motility, invasion, and egress, which are the major steps of its lytic cycle.

The motile and invasive stages of apicomplexan parasites, including *P. falciparum* and *T. gondii*, rely on gliding motility to migrate, invade the host cells, egress from the infected cell, and disperse ^45^. Tachyzoites shed surface proteins like SAG1 during gliding, which can be used to track gliding trails. For assessing motility, RHTIR1-VMP1AID parasites were grown with (+IAA) or without (-IAA) IAA, purified, allowed to glide on FBS-coated coverslips, fixed, and stained for SAG1. 64.03% of the -IAA RHTIR1-VMP1AID tachyzoites exhibited gliding motility with an average trail length of 53.0 µm as compared with 21.8% of the +IAA RHTIR1-VMP1AID tachyzoites with an average trail length of 25.5 µm (Figures 3A-3C), indicating a significant defect in gliding motility upon TgVMP1 depletion. Gliding motility is powered by glideosome, an actomyosin motor between the parasite plasma membrane and inner membrane complex. The plasma membrane protein SAG1 and inner membrane complex protein 1 (IMC1) showed similar localization in +IAA and –IAA RHTIR1-VMP1AID tachyzoites (**Figure S5**), indicating that these two cellular structures are unaffected in TgVMP1-depleted parasites and unlikely to be responsible for the gliding motility defect.

**Figure 3.**
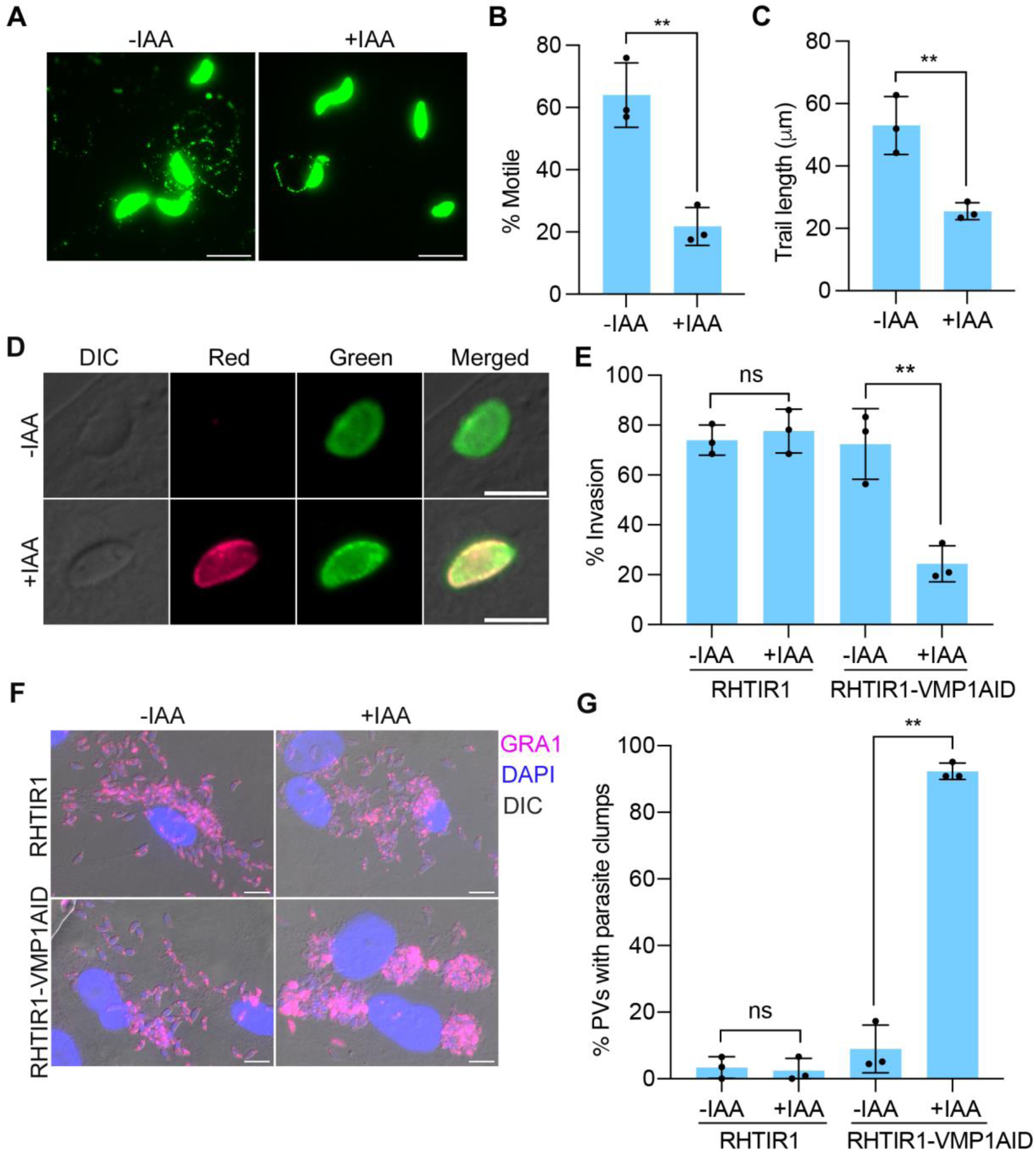
TgVMP1 is crucial for gliding motility, invasion, and egress. RHTIR1 and RHTIR1-VMP1AID parasites were cultured with (+IAA) or without (-IAA) IAA, and assessed for gliding motility, host cell invasion, and egress. **A.** Purified +IAA and –IAA RHTIR1-VMP1AID tachyzoites were allowed to glide on FBS-coated coverslips, processed for IFA using anti-SAG1 antibodies, and observed for gliding trails. The representative images show gliding trails (scale bar = 5 µm). **B.** The parasites showing gliding trails in “A” were counted and plotted as a percentage of the total number of parasites (% Motile) on y-axis for the indicated parasites on x-axis. Each data is based on at least 100 parasites in each of the 3 independent experiments. **C.** The length of gliding trails in “A” was measured and shown as mean trail length (Trail length (μm)) with SD error bar on y-axis for the indicated parasites on x-axis. Each data is based on at least 90 parasites in each of the 3 independent experiments. **D.** Purified +IAA and –IAA tachyzoites were allowed to invade HFF cells, fixed, and labelled with a red fluorophore-conjugated antibody, which would label extracellular parasites (Red). The parasites were permeabilized and labelled again with a green fluorophore-conjugated antibody, which would label both extracellular and invaded parasites (Green). The representative images show invaded or extracellular parasites. **E.** The parasites in “D” were scored for different labels, and the number of invaded parasites is shown as a percentage of the total number of parasites (% Invasion) on y-axis for the indicated parasites on x-axis. Each data is mean with SD error bar based on at least 100 parasites from each of the 3 independent experiments. **F.** Intracellular RHTIR1 and RHTIR1-VMP1AID parasites were grown with (+IAA) or without (-IAA) IAA, treated with ionomycin to induce egress, fixed, and stained with antibodies to the PV protein GRA1 and nuclear stain DAPI. The representative images for the indicated parasites show PVs from which parasites egressed and dispersed or did not egress and remained as clumps. Scale bar = 10 µm. **G.** The number of PVs in “F” was determined, and the number of parasite clumps is shown as a percentage of the total number of PVs (% PVs with parasite clumps) on y-axis for the indicated parasites on x-axis. Each data is mean with SD error bar based on at least 100 PVs from each of the 3 independent experiments. The significance of difference between two groups is shown as p-value (** = p < 0.01; ns: not significant).

For assessing invasion, purified +IAA and –IAA tachyzoites were allowed to invade HFF cells, fixed, and labelled for extracellular and invaded parasites. The +IAA RHTIR1-VMP1AID parasites showed significantly reduced invasion efficiency as compared with –IAA RHTIR1-VMP1AID parasites and ±IAA RHTIR1 parasites (Figures 3D and **3E**), indicating that TgVMP1 is crucial for host cell invasion.

For assessing egress, +IAA and –IAA intracellular RHTIR1-VMP1AID or RHTIR1 tachyzoites were treated with the calcium ionophore ionomycin, which has been used to induce rapid egress of intracellular parasites ^46^, and processed for IFA using antibodies to the PV protein GRA1. RHTIR1 tachyzoites showed dispersal irrespective of the IAA treatment. The majority of -IAA RHTIR1-VMP1AID parasites were dispersed away from the PV, whereas +IAA RHTIR1-VMP1AID parasites remained as clumps near the PV (Figures 3F and **3G**), indicating a role of TgVMP1 in egress.

### TgVMP1 depletion did not affect conoid extrusion

Conoid extrusion, gliding motility, and secretions of micronemes and rhoptries are key sequential events for host cell invasion by *T. gondii*. Conoid is a hollow cone-like structure of tubulin fibers in the apical complex of *T. gondii* and certain other apicomplexans ^47^. Conoid extrusion regulates motility, invasion, and egress by activating microneme secretion and actomyosin motor ^3^. Since TgVMP1 depletion decreased gliding motility, invasion, and egress, we compared purified –IAA and +IAA RHTIR1-VMP1AID tachyzoites for conoid extrusion by treating with the calcium ionophore A23187, which induces conoid extrusion ^48^. The number of parasites showing conoid extrusion was similar in +IAA and –IAA parasites (**Figures S6A** and **6B**), ruling out a role of TgVMP1 in this key event.

### TgVMP1 is crucial for microneme secretion

Decreased motility, host cell invasion, egress, and intracellular development of TgVMP1-depleted parasites prompted us to look into the effect of TgVMP1 depletion on micronemes, rhoptries, and dense granules, which are crucial for these processes in *Toxoplasma* and *Plasmodium* ^2,49^. Micronemes are situated at the apical end, and their secretion is required for motility, invasion, and egress ^3^. We compared the localization and secretion of selected microneme proteins in +IAA and –IAA RHTIR1-VMP1AID tachyzoites. For microneme organization, +IAA and -IAA intracellular parasites were processed for IFA and compared for localization of selected microneme proteins (apical membrane antigen 1 (AMA1), microneme protein 2 (MIC2), and the microneme protein 3 (MIC3)). All these proteins showed similar predominant apical localization in +IAA and –IAA parasites, ruling out a role of TgVMP1 in microneme organization (Figures 4A-4D; **Figures S7A** and **7B**). For microneme secretion, purified +IAA and –IAA RHTIR1-VMP1AID parasites were treated with ethanol, which induces microneme secretion that is known as extracellular secreted antigens (ESA) ^50^. We used MIC2 as an ESA marker, which is proteolytically processed by the microneme protein proteases MPP1 and MPP2, and predominantly present in the shorter form in ESA ^51^. The level of larger MIC2 form were higher in the pellet fraction of +IAA parasites than that of –IAA parasites (Figures 4E and **4F**). The ESA fraction of +IAA parasites had less amount of the processed form of MIC2 along with the larger MIC2 form as compared with the presence of only processed form of MIC2 in –IAA parasites (Figures 4E and **4G**), which indicated both impaired microneme secretion and processing of MIC2 upon TgVMP1 depletion.

**Figure 4.**
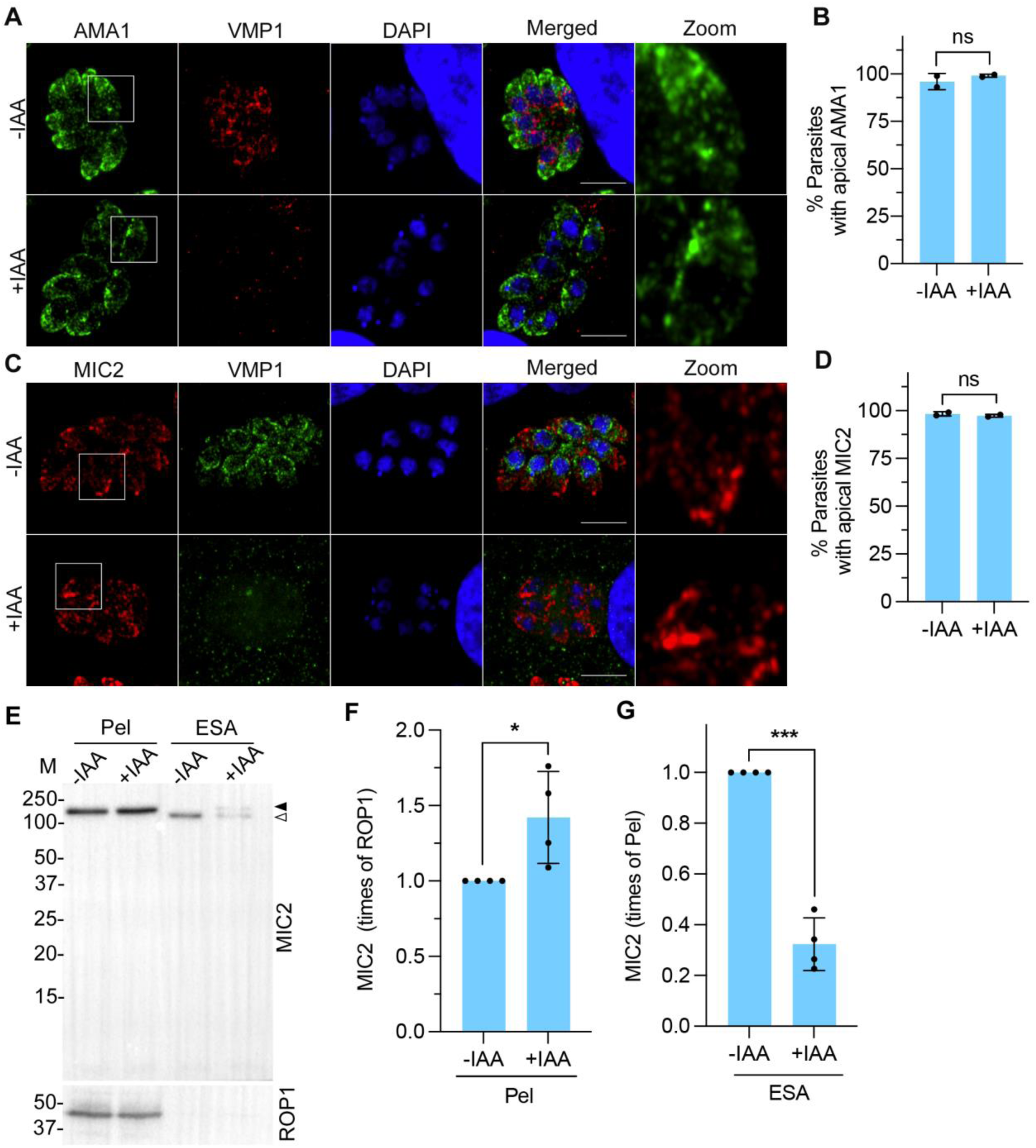
TgVMP1 depletion did not affect microneme organization, but impaired microneme secretion. -IAA and +IAA RHTIR1-VMP1AID parasites were evaluated for microneme organization and secretion using microneme proteins as markers. **A** and **C.** -IAA and +IAA intracellular RHTIR1-VMP1AID parasites were assessed for microneme organization by labelling the parasite with antibodies to AMA1 (**A**) and MIC2 (**C**). The representative Airyscan micrographs show localization of AMA1 or MIC2, TgVMP1AIDHA (VMP1), nucleus (DAPI), and overlap of the three images (Merged). Scale bar = 10 µm, and zoom-in of the boxed area in AMA1 or MIC2 panel is shown as “Zoom”. Note the similar apical localization of AMA1 and MIC2 in –IAA and +IAA parasites. **B** and **D.** The number of parasites with apical localization of AMA1 (in A) or MIC2 (in C) was determined, and shown as a percentage of the total number of parasites observed (% Parasites with apical AMA1 or MIC2) on y-axis for the indicated parasites on x-axis. Each data is mean with SD error bar based on at least 100 observations from each of the 2 independent experiments. **E.** Equal number of purified -IAA and +IAA RHTIR1-VMP1AID parasites were incubated with 1% ethanol to induce microneme secretion. The ESA and pellet (Pel) fractions were assessed for the presence of MIC2 and ROP1 as a control by western blotting. The immunoblot shows the larger form of MIC2 in pellet fraction of both the parasites, processed form of the MIC2 in ESA fraction of -IAA parasites, and both larger (filled arrowhead) and processed (empty arrowhead) forms of MIC2 in ESA fraction of +IAA parasites. The protein size markers (M) are in kDa. **F.** The signal intensities of ROP1 and MIC2 in the pellet fractions in “E” were measured, the signal intensity of MIC2 was normalized with that of ROP1 for the corresponding sample, and plotted as times of the signal intensity of ROP1 (MIC2 (times of ROP1)) on y-axis for the pellet fractions (Pel) of indicated parasites on x-axis. **G.** The signal intensities of MIC2 in ESA fractions were measured, and plotted as times of the signal intensity of MIC2 in the corresponding pellet fraction (MIC2 (times of Pel) on y-axis for the indicated parasites on x-axis. Each data is mean with SD error bar from 4 independent experiments. The significance of difference between two data sets is indicated with p-value (*= P<0.05, *** = P < 0.001, ns: not significant).

### TgVMP1 is crucial for rhoptry biogenesis and secretion

Rhoptries are situated at the apical end and their secretion is essential for host cell invasion, formation of PV around the parasite inside the host cell, and modulation of host cell responses ^4,5^. We compared intracellular +IAA and –IAA RHTIR1-VMP1AID parasites for rhoptry organization and secretion using ROP1 as a marker. For rhoptry organization, intracellular +IAA and –IAA tachyzoites were processed for IFA using anti-ROP1 antibodies or transmission electron microscopy (TEM). ROP1 was localized to distinct elongated structures with a bulbous base, the characteristic rhoptry shape, in the majority (85.3%) of -IAA parasites as compared with only a small number (11.3%) of +IAA parasites (Figures 5A and **5B**). Moreover, the ROP1 signal was associated with fragmented structures in purified +IAA parasites as opposed to distinct elongated structures in -IAA parasites (Figure 5C). The TEM images of +IAA parasites confirmed the absence of rhoptries in the majority of parasites, whereas -IAA parasites had characteristic rhoptries (Figure 5D). The loss of rhoptries in TgVMP1-depleted parasites indicates a crucial role for TgVMP1 in rhoptry formation.

**Figure 5.**
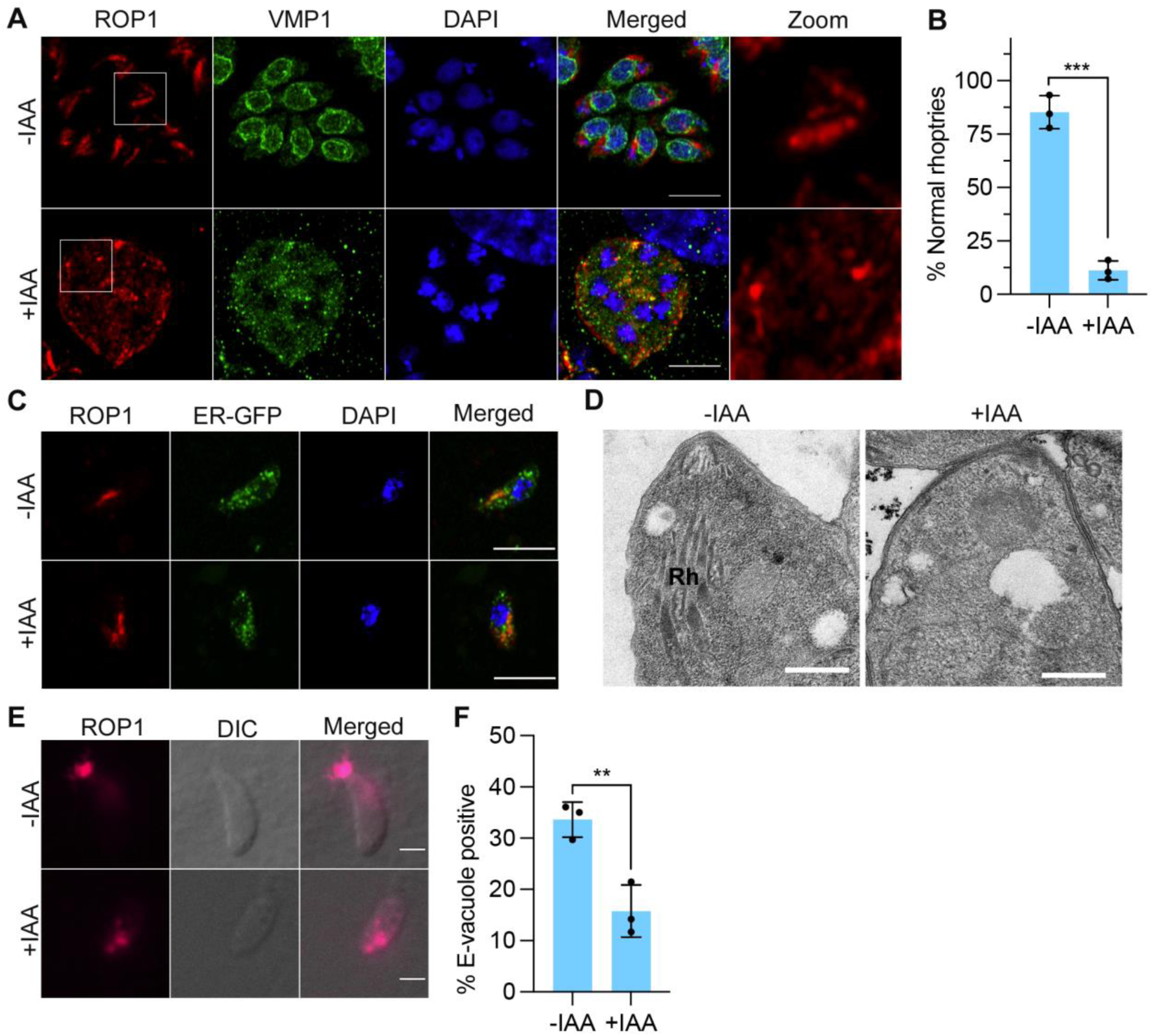
TgVMP1 is crucial for rhoptry biogenesis and secretion. +IAA and –IAA RHTIR1-VMP1AID parasites were compared for rhoptry organization and secretion. **A.** Intracellular +IAA and –IAA RHTIR1-VMP1AID parasites were examined for rhoptries using antibodies to ROP1. The representative Airyscan micrographs show rhoptries (ROP1), TgVMP1AIDHA (VMP1), nucleus (DAPI), and overlap of all the three images (Merged). Scale bar = 10 µm, and the boxed region of ROP1 panel is zoomed-in (Zoom). Note that ROP1 is restricted to elongated structures with a bulbous base in -IAA parasites, as compared with predominantly fragmented structures in +IAA parasites. **B.** The number of parasites showing ROP1-labelled elongated structures with a bulbous base in “A**”** was determined, and plotted as a percentage of the total number of parasites observed (% Normal rhoptries) on y-axis for the indicated parasites on x-axis. Each data is mean with SD error bar based on at least 100 parasites from each of the 3 independent experiments. **C**. The representative merged Airyscan confocal micrographs show signal of ROP1, ER-GFP, and nuclear stain DAPI in purified -IAA and +IAA parasites. Scale bar = 5 µm. Note the ROP1-labeled fragmented structures in +IAA panel as compared with the elongated structure in –IAA panel. **D.** The representative TEM micrographs show rhoptries (Rh) in -IAA parasites, which are absent in +IAA parasites. Scale bar is 500 nm. These observations are based on multiple -IAA (n=117) and +IAA (n=78) parasites from 3 independent experiments. **E.** Cytochalasin D (CytD)-treated purified -IAA and +IAA RHTIR1-VMP1AID parasites were incubated with HFF cells, and labelled with anti-ROP1 antibodies. The representative micrographs show ROP1 signal (ROP1), bright field (DIC), and overlap of the two panels (Merged) for the indicated parasites. Scale bar = 2 µm. Note the ROP1 signal in the host cell close to the parasite apical end, which represents e-vacuoles, in –IAA parasites. On the other hand, ROP1 signal remained inside the +IAA parasites. **F.** The number of parasites exhibiting e-vacuoles, in “E**”** was determined, and shown as a percentage of the total parasites observed (% E-vacuole positive) on y-axis for the indicated parasites on x-axis. Each data is mean with SD error bar based on at least 100 parasites from each of the 3 independent experiments. The significance of difference between the data sets is indicated with p-value (**= P<0.01; ***= P<0.001).

For rhoptry secretion, we checked for ROP1-labelled e-vacuoles, which are discharged into the host cell during invasion. E-vacuole discharge can be monitored by treating the parasites with cytochalasin D (CytD) that prevents invasion without affecting e-vacuole secretion. The CytD-treated purified +IAA and –IAA parasites were allowed to invade HFF cells, the cells were processed for IFA using anti-ROP1 antibodies, and scored for ROP1 positive e-vacuoles. ∼16% of the +IAA parasites secreted e-vacuoles into the host cell as compared with ∼34% of the –IAA parasites (Figures 5E and **5F**), indicating a crucial role for TgVMP1 in rhoptry secretion.

### TgVMP1 is required for IVN formation and dense granule biogenesis

Dense granules are secretory organelles throughout the parasite cytoplasm, which secrete an arsenal of secretory factors, collectively called GRA proteins, into the PV lumen immediately after invasion and during parasite development ^6,9,10^. Dense granule secretion contributes to the formation of IVN, which has been suggested to act as a scaffold to position and organize the replicating parasites within the PV ^9,10^. Some GRA proteins have been shown to be transported to the PVM and host cell wherein they mediate transport and modulate host processes, respectively ^6,49^. We evaluated intracellular +IAA and –IAA RHTIR1-VMP1AID parasites for IVN organization by IFA using the IVN-associated proteins GRA1 and GRA2 as markers, and by TEM. Both GRA1 and GRA2 were localized to distinct tubulo-vesicular IVN in the PV of -IAA parasites, whereas these proteins showed diffuse localization in the PV of +IAA parasites (Figures 6A and **S8A**). The number of parasites with GRA1- and GRA2-associated IVN was significantly lower in +IAA parasites (16.5% for GRA1, 21.0% for GRA2) than that in –IAA parasites (90.4% for GRA1, 85.4% for GRA2) (Figures 6B and **S8B**). The loss of IVN was also reflected in TEM micrographs of +IAA parasites (Figure 6C), which indicates a crucial role for TgVMP1 in IVN formation.

**Figure 6.**
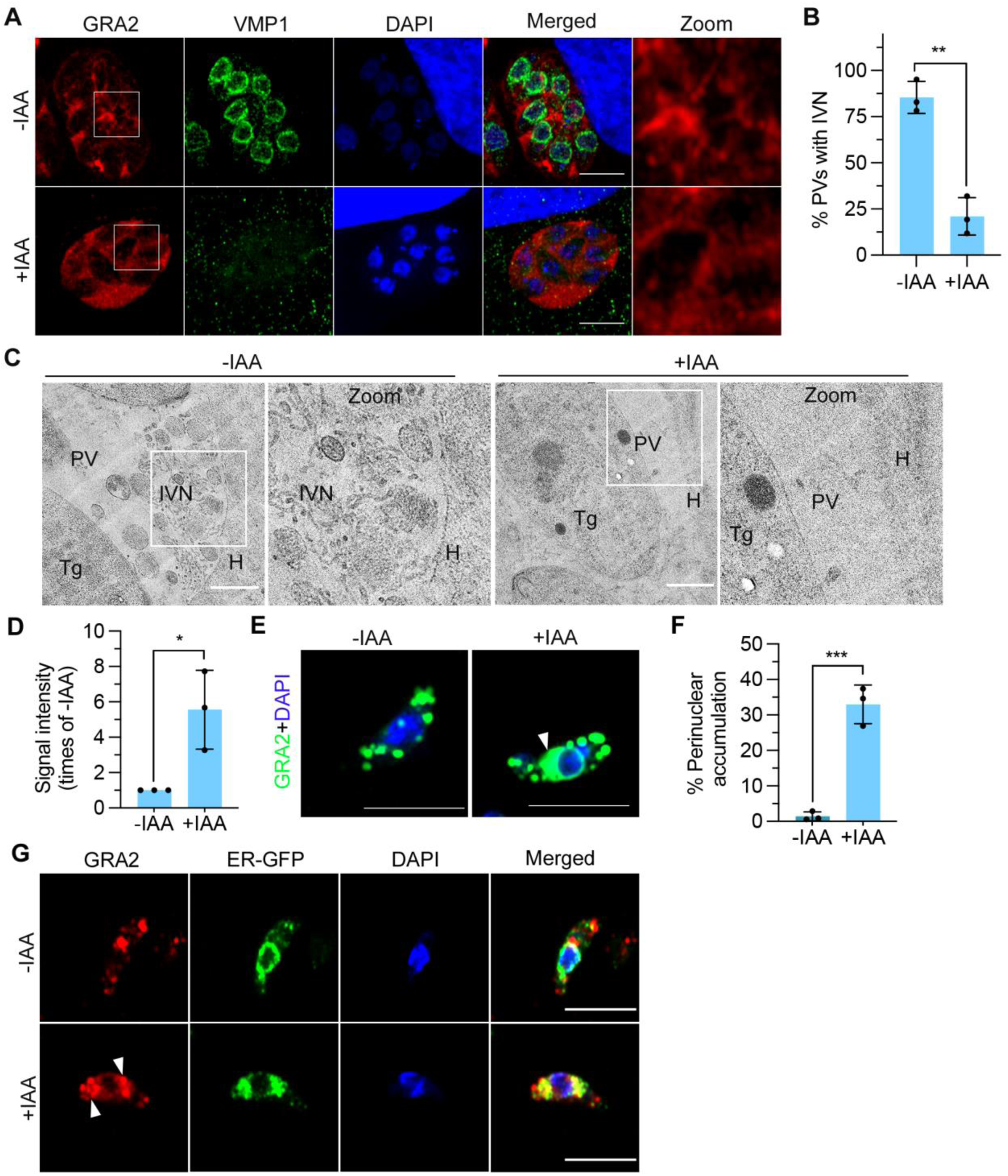
TgVMP1 is crucial for IVN formation and dense granule biogenesis. -IAA and +IAA RHTIR1-VMP1AID parasites were evaluated for IVN and dense granule biogenesis using the dense granule protein GRA2 as a marker. **A.** Intracellular –IAA and +IAA parasites were processed for IFA using antibodies to GRA2, and observed for IVN. The representative Airyscan micrographs show localization of GRA2, TgVMP1AIDHA (VMP1), nucleus (DAPI), and overlap of all the three images (Merged). Scale bar is 10 µm, and the boxed area in GRA2 panel is zoomed-in (Zoom). Note the GRA2-labeled distinct tubulo-vesicular IVN in the PV of –IAA parasites, whereas GRA2 signal is diffuse and associated with fragmented structures in the PV of +IAA parasites. **B.** The number of PVs showing IVN in “A” was determined and plotted as a percentage of the total number of PVs observed (% PVs with IVN) on y-axis for the indicated parasites on x-axis. Each data is mean with SD error bar based on ≥50 PVs from each of the 3 independent experiments. **C.** The TEM micrographs show tubulo-vesicular IVN in the PVs of intracellular –IAA parasites, which are absent in +IAA parasites. Scale bar = 500 nm, and the boxed area is zoomed-in (Zoom). The images represent multiple observations (-IAA, n = 37; +IAA, n = 19) in 2 independent experiments. The labels are: Tg, *T. gondii*; PV; IVN; H, host cell. **D.** Purified -IAA and +IAA RHTIR1-VMP1AID parasites were processed for GRA2 localization by IFA, and assessed for the signal intensity of GRA2. The signal intensity of GRA2 in each of the -IAA and +IAA parasites was measured, and the mean signal intensity of GRA2 in +IAA parasites was plotted as times of the signal intensity of -IAA parasites (Signal intensity (times of –IAA)) on y-axis for the indicated parasites on x-axis. Each data is mean signal intensity with SD error bar based on multiple parasites (-IAA, n=338; +IAA, n=383) from 3 independent experiments. **E.** The representative Airyscan micrographs show merged image of GRA2 and DAPI panels for –IAA and +IAA parasites. Scale bar is 5 µm. The arrowhead indicates perinuclear accumulation of GRA2 in +IAA parasites, which is absent in -IAA parasites. **F.** The number of parasites showing perinuclear accumulation of GRA2 in “E” was plotted as a percentage of the total number of parasites observed (% Perinuclear accumulation) on y-axis for the indicated parasites on x-axis. Each data is based on multiple parasites (-IAA, n=338; +IAA, n=383) from 3 independent experiments. **G.** The representative Airyscan micrographs show localization of GRA2, ER-GFP, DAPI, and overlap of the three images (Merged) in the indicated parasites. Scale bar is 5 µm. Note the prominent co-localization of GRA2 and ER-GFP in the perinuclear region in +IAA parasites, which is nearly absent in -IAA parasites. The significance of difference between the data sets is indicated with p-value (*= P<0.05, **= P<0.01, ***= P<0.001).

For dense granule biogenesis, purified +IAA and –IAA tachyzoites were processed for IFA and compared for the levels of intracellular GRA1 and GRA2. +IAA parasites showed higher levels of these proteins than –IAA parasites (Figures 6D and **S8C)**. -IAA RHTIR1-VMP1AID parasites showed distinct GRA1- or GRA2-labeled dense granules in the cytoplasm (Figures 6E and **S8D)**. On the other hand, in addition to the cytoplasmic GRA1- and GRA2-labeled dense granules, +IAA RHTIR1-VMP1AID parasites also showed perinuclear accumulation of GRA1 and GRA2 (Figures 6E and **S8D**). The perinuclear accumulation of GRA1 or GRA2 was significantly higher in +IAA parasites than that in –IAA parasites (Figures 6F and **S8E**), and co-localized with ER-GFP (Figures 6G and **S8F**), which indicated a key role of TgVMP1 in dense granule biogenesis.

### TgVMP1-depletion altered LD homeostasis

The DedA superfamily proteins are proposed to be lipid scramblases; human VMP1 and TMEM41B have been shown to have roles in lipoprotein and lipid transport, LD biogenesis, and lipid distribution ^22–24^. This prompted us to evaluate the effect of TgVMP1 depletion on lipid homeostasis using LDs as a marker. LDs contain neutral lipids such as triglycerides and cholesterol esters, are formed in the ER bilayer, and bud off into the cytoplasm ^52^. We stained purified +IAA and –IAA RHTIR1-VMP1AID tachyzoites with Nile red to label LDs ^53^, and observed the parasites by live cell confocal microscopy. +IAA parasites had a significant reduction in the number and increase in the size of LDs as compared with -IAA parasites (Figures 7A-7C). TEM micrographs also showed larger size LDs in +IAA parasites as compared with -IAA parasites (Figure 7D), indicating a role for TgVMP1 in LD homeostasis. Although we attempted to determine membrane lipid distribution in TgVMP1-depleted parasites using click lipids and fluorophore, we did not succeed due to widespread non-specific labelling of the parasite by click fluorophores. Further optimization of the use of click lipids or alternative approaches is required to determine if TgVMP1-depletion affects membrane lipid distribution.

**Figure 7.**
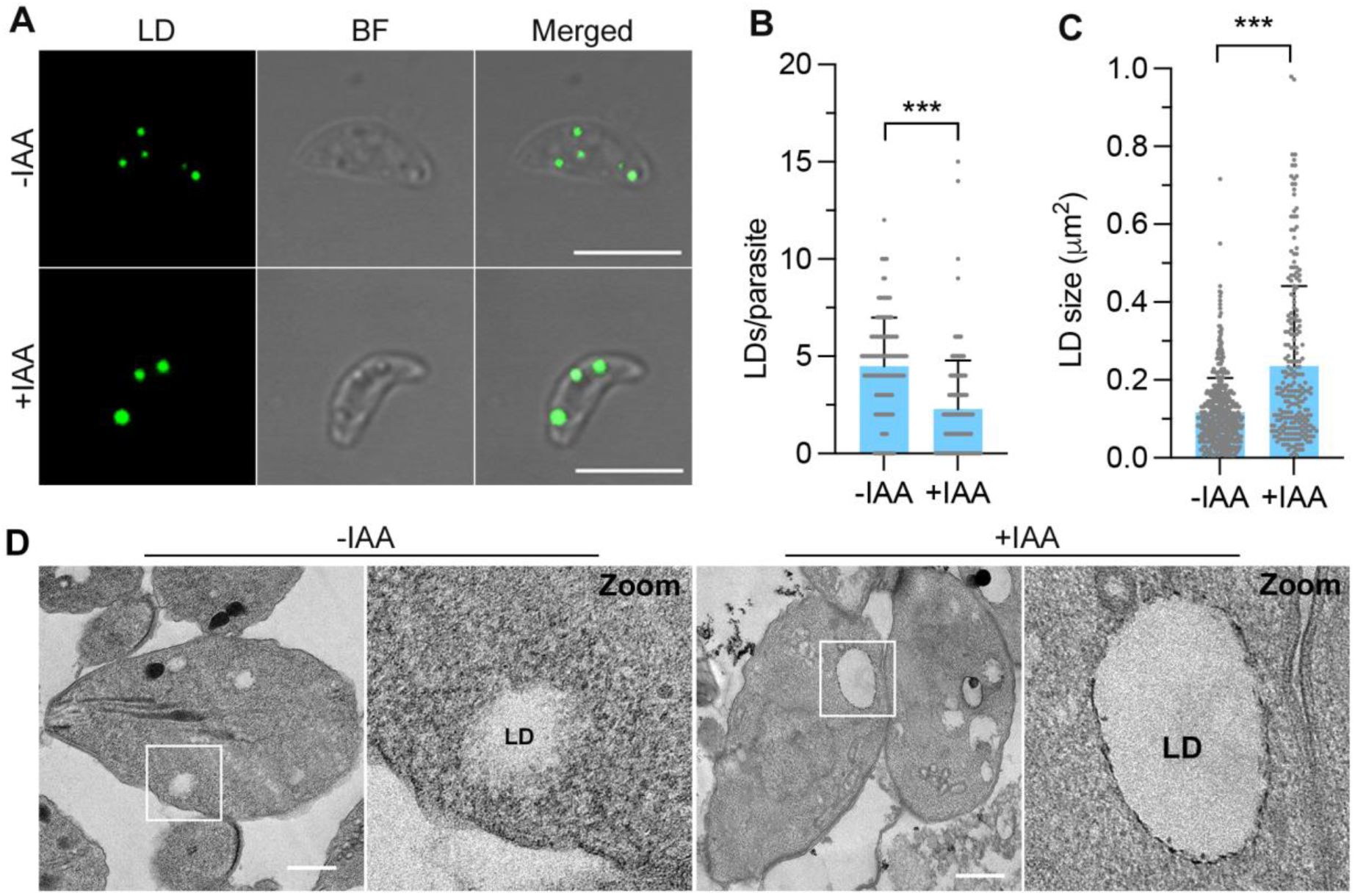
TgVMP1 depletion affected LD homeostasis. Purified +IAA and –IAA RHTIR1-VMP1AID parasites were evaluated for LDs by Nile red staining or TEM. **A.** The representative confocal micrographs show signal for Nile red-stained LDs, bright field (BF), and overlap of the two images (Merged). Scale bar is 5 µm. **B.** Number of LDs in “A” was determined, and shown as the number of LDs/parasite (LDs/parasite) on y-axis for the indicated parasites on x-axis. Each data is mean with SD error bar based on multiple parasites from 2 independent experiments (-IAA, n = 137; +IAA, n = 140). **C.** The size of each LD in “A” was measured, and plotted as mean LD size (LD size (μm^2^)) on y-axis for the indicated parasites on x-axis. Each data is mean with SD error bar based on multiple parasites from 2 experiments (-IAA, n = 473; +IAA, n = 232). **D.** The representative TEM micrographs show LDs in the indicated parasites. Scale bar is 500 nm, and the boxed area is zoomed-in (Zoom). Note the larger size LDs in +IAA parasites as compared with –IAA parasites. Each data represents multiple parasites in 2 independent experiments (-IAA, n = 265; +IAA, n = 115). The significance of difference between two data sets in indicated with p-value (***= P<0.001).

### TgVMP1 depletion disrupted ER organization

The microneme, rhoptry, and dense granule proteins have been shown to be synthesized in ER, and vesicles containing these proteins bud off the ER/Golgi regions and follow endosomal vesicular trafficking pathway to reach/mature into these organelles ^11^. Perinuclear clustering of dense granule proteins GRA1 and GRA2 with ER-GFP in TgVMP1-depleted parasites prompted us to investigate whether TgVMP1 depletion compromised the ER. We compared the localization of ER-GFP in intracellular -IAA and +IAA RHTIR-VMP1AID::BiP-GFP-HDEL parasites by IFA. –IAA parasites showed uniform ER-GFP localization around the nucleus, which was also extended toward the apical region (**Figures S9A** and **S9B**). On the other hand, ER-GFP was prominently clustered around the nucleus in a significant number of +IAA parasites (**Figures S9A** and **S9B**), indicating a defect in ER organization, which might contribute to various phenotypes observed in TgVMP1-depleted parasites. This data highlights a key role of TgVMP1 in the ER functioning, which could include the maintenance of ER structure, formation of functional subdomains, and communication with other organelles.

### TgVMP1 interactome supports its roles in the secretory organelle function and biogenesis

To understand how TgVMP1 depletion might impair the function and biogenesis of secretory organelles, we determined and analyzed the TgVMP1 interactome (**Figure S10, Table S2**). TgVMP1 interactome contained proteins that have been shown to localize to micronemes (MIC1, MIC3, and MIC7), rhoptries (ROP8, ROP9, ROP12, ROP13, ROP26, ROP37, armadillo repeats only protein, RON3, and CARP), and dense granules (GRA8, GRA12, GRA25, and GRA71). The interactome also contained the proteins associated with gliding motility (PhiL1, GAPM2B, GAP45, GAPM1A, myosin-light-chain kinase, GRA12, GRA8), processing of the microneme and rhoptry proteins (aspartyl protease ASP3, V-type proton ATPase subunit a1), and vesicle transport (sortilin-like receptor and Rab2). Additionally, TgVMP1 interactome contained a large number of proteins related to protein biosynthesis, cytoskeleton, maintenance of the parasite shape, and virulence. Among these proteins, the *T. gondii* armadillo repeats only protein (TgARO), sortilin-like receptor (TgSORTLR), V-type proton ATPase, and aspartyl protease ASP3 are specifically relevant in the context of secretory organelles. TgSORTLR is required for trafficking of the microneme and rhoptries proteins, and TgSORTLR knock-down impaired the biogenesis of these organelles ^54^. TgARO is required for apical positioning of rhoptries ^55^, most likely, via its interaction with myosin F, which is also present in the TgVMP1 interactome. The V-type proton ATPase is localized to the plasma membrane and endosomal compartments, and required for the processing of microneme and rhoptry proteins during their transport through the endosomal compartments ^56^. Furthermore, the V-type proton ATPase is also required for activation of the proteases that process microneme and rhoptry proteins. The aspartyl protease ASP3 is present in endosomal compartments, and processes several microneme and rhoptry proteins during their transport through the endosomal compartments to the respective organelles ^57^. The presence of proteins localized to/associated with micronemes, rhoptries, dense granules and the motor complex in TgVMP1 interactome suggests a direct or indirect role of TgVMP1 in functioning and biogenesis of these organelles, which is in agreement with the defects observed in these organelles upon TgVMP1 depletion. The presence of several motility-associated proteins in the TgVMP1 interactome suggests an alternative mechanism by which TgVMP1 could contribute to parasite motility, however, this requires further investigation.

### TgVMP1 and PfVMP1 are lipid scramblases, and restoration of the ER-localized scramblase activity rescued TgVMP1-depleted parasites

TgVMP1-depleted parasites showed dense granule accumulation in the ER, impaired LD homeostasis, and disrupted ER organization, suggesting a role for TgVMP1 in ER and ER-organelle contact sites. The ER-localized DedA superfamily proteins HsVMP1 and HsTMEM41B have been shown to have lipid scramblase activity, which has been proposed to be required for lipoprotein and lipid transport, LD biogenesis, lipid distribution, and the formation of viral replication organelles ^22,23,35,26^. TgVMP1, PfVMP1 and HsVMP1 share similar domain architecture and AlphaFold structure, which prompted us to ask whether TgVMP1 and PfVMP1 function as lipid scramblases and restoration of the ER-localized scramblase activity in RHTIR1-VMP1AID parasites could rescue the growth defects.

Lipid scramblases translocate lipids between two membrane leaflets, which can be measured by monitoring the fluorescence signal intensity of fluorescent phospholipid (NBD-PS)-containing liposomes in the presence or absence of a membrane impermeable quencher (dithionite). The fluorescence signal of NBD-PS on the outer liposome leaflet would be quenched by dithionite, a membrane impermeable reducing agent, causing ∼50% decrease in fluorescence intensity (Figure 8A). On the other hand, translocation of NBD-PS between the outer and inner leaflets of the liposome by a lipid scramblase would result in the >>50% decrease in fluorescence intensity (Figure 8A). The addition of Triton X-100 would permeabilize the liposomes, allowing dithionite to enter and quench the NBD-PS fluorescence completely (Figure 8A). For lipid scramblase activity, PfVMP1AID_HA_ and TgVMP1AID_HA_ were purified from RHTIR1-VMP1AID and PfVMP1AID parasites, respectively (**Figure S11A**). Purified TgVMP1AID_HA_ or PfVMP1AID_HA_ was incorporated into NBD-PS-containing liposomes to obtain TgVMP1AID_HA_-liposomes or PfVMP1AID_HA_-liposomes, and only NBD-PS-containing liposomes (empty-liposomes) were used as a control. Indeed, TgVMP1-liposomes and PfVMP1-liposomes showed >>50% decrease in fluorescence intensity upon addition of dithionite as compared with ∼50% decrease in fluorescence intensity of empty-liposomes (Figure 8B), indicating that TgVMP1 and PfVMP1 are lipid scramblases.

**Figure 8.**
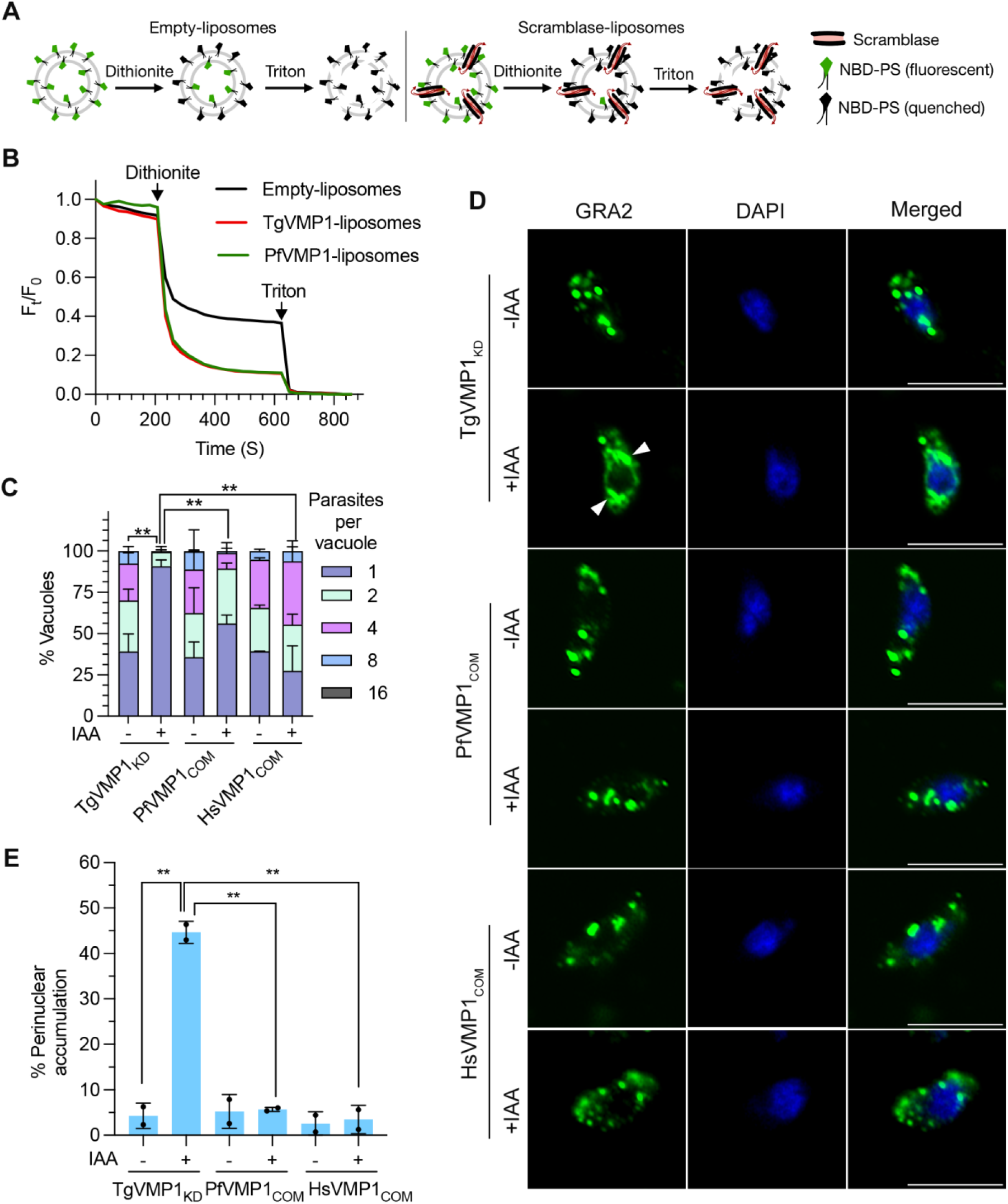
TgVMP1 and PfVMP1 are lipid scramblases, and functionally conserved along with HsVMP1. **A.** The schematic shows empty-liposomes and scramblase-liposomes with the indicated constituents. Addition of dithionite, a membrane impermeable reducing agent, would quench the fluorescence of NBD-PS on the outer leaflet of the liposome, causing ∼50% decrease in the fluorescence signal intensity of the empty-liposomes. On the other hand, continuous translocation of NBD-PS between the outer and inner leaflets of the scramblase-liposomes by the scramblase would cause >>50% decrease in fluorescence signal intensity. Addition of Triton X-100 would make the liposome membrane permeable to dithionite, causing complete quenching of NBD-PS, which would decrease fluorescence signal intensity to almost zero. **B.** The plot shows relative fluorescence intensities (Ft/F0) of empty-liposomes, TgVMP1AIDHA-containing liposomes (TgVMP1-liposomes), and PfVMP1AIDHA-containing liposomes (PfVMP1-liposomes) before, after addition of dithionite, and after addition of Triton X-100. Arrows indicate the time points at which dithionite or Triton X-100 was added to the liposomes. A decrease of >>50% in the fluorescence signal intensities of the TgVMP1-liposomes and PfVMP1-liposomes upon addition of dithionite indicates that TgVMP1 and PfVMP1 have lipid scramblase activity. Each data is average of 3 independent scramblase assays. **C.** RHTIR1-VMP1AID (TgVMP1KD), RHTIR1-VMP1AID/PfVMP1 (PfVMP1COM), and RHTIR1-VMP1AID/HsVMP1 (HsVMP1COM) tachyzoites were cultured in the presence (+IAA) or absence (-IAA) of IAA, stained for the PV marker GRA1, and the number of parasites/PV was counted. The number of PVs containing different number of parasites was plotted as a percentage of the total number of PVs (% Vacuoles) on y-axis for the indicated parasites on x-axis. Each data is mean with SD error bar based on at least 100 PVs in each of 3 independent experiments. The significance of difference is for the PVs containing a single parasite only. **D.** The indicated parasites were cultured with (+IAA) or without (-IAA) IAA, tachyzoites were purified and evaluated for dense granule biogenesis defect using GRA2 as a marker. The Airyscan micrographs show signal for GRA2, nuclear stain DAPI, and the merged image of the GRA2 and DAPI panels. Scale bar is 5 µm. The arrowheads indicate perinuclear accumulation of GRA2 in +IAA TgVMP1KD, but not in +IAA PfVMP1COM and +IAA HsVMP1COM parasites, indicating restoration of the dense granule biogenesis by PfVMP1 and HsVMP1, respectively. **E.** The number of parasites showing perinuclear accumulation of GRA2 in “D” was determined, and plotted as a percentage of the total number of parasites observed (% Perinuclear accumulation) on y-axis for the indicated parasites on x-axis. Each data is based on at least 100 parasites in each of the 2 independent experiments. The significance of difference between the data sets is indicated with p-value (**= P<0.01).

For restoration of the ER-localized scramblase activity in RHTIR1-VMP1AID parasites, we ectopically expressed Myc-tagged PfVMP1 (PfVMP1_Myc_) and HsVMP1 (HsVMP1_Myc_) proteins in RHTIR1-VMP1AID parasites to obtain complemented parasites (RHTIR1-VMP1AID/PfVMP1_Myc_ and RHTIR1-VMP1AID/HsVMP1_Myc_, respectively). We next compared the complemented parasites with RHTIR1-VMP1AID parasites for overall growth and intracellular development in the presence (+IAA) or absence (-IAA) of IAA. The complemented parasites expressed PfVMP1_Myc_ or HsVMP1_Myc_ (**Figure S11B**), which co-localized with TgVMP1AID_HA_ (**Figure S11C**), indicating that these proteins retain ER localization, even in a heterologous system. As expected, all the three parasites formed plaques under –IAA condition, whereas only complemented parasites, but not RHTIR1-VMP1AID parasites, formed plaques under +IAA condition (**Figure S11D**), which indicated restoration of the overall growth by PfVMP1 or HsVMP1 in the absence of TgVMP1. 90.6% of the +IAA RHTIR1-VMP1AID parasites were arrested at single parasite/PV stage as compared to 27.5% (for HsVMP1) and 56.1% (for PfVMP1) of the +IAA complemented parasites (Figure 8C), indicating restoration of the intracellular development by PfVMP1 or HsVMP1 in the absence of TgVMP1. We next checked if PfVMP1 and HsVMP1 could reverse the perinuclear accumulation of the dense granule protein GRA2 in TgVMP1-depleted parasites. <5.7% of the +IAA complemented parasites showed perinuclear accumulation of GRA2 as compared with 44.7% of the +IAA RHTIR1-VMP1AID parasites (Figures 8D and **8E**), indicating rescue of the dense granule biogenesis defect. The rescue of growth and dense granule biogenesis defects of TgVMP1-depleted parasites by PfVMP1 and HsVMP1 indicate their functional conservation. This provides a mechanistic proof for a critical role of the ER-localized scramblase activity in the biogenesis and function of apicomplexan secretory organelles.

## Discussion

Micronemes, rhoptries, and dense granules are specialized secretory organelles in apicomplexans, and have essential roles in parasite motility, invasion of the host cells, development, and virulence ^2–4,6,49^. ER is the site for synthesis of the proteins and biogenesis of the vesicles destined to these organelles, which would require lipid mobilization. Since DedA superfamily proteins are implicated in lipid distribution and organelle biogenesis ^13,14^, we identified and investigated the *P. falciparum* and *T. gondii* DedA superfamily protein VMP1 towards understanding its role in the biogenesis and function of secretory organelles. We show that PfVMP1 and TgVMP1 are ER-resident lipid scramblases. TgVMP1 is crucial for the biogenesis of rhoptries and dense granules, and secretion of micronemes, rhoptries, and dense granules **(Figure S13)**. Notably, restoration of the ER-resident scramblase activity by complementing TgVMP1-depleted parasites with PfVMP1 or HsVMP1 alleviated the growth and dense granule biogenesis defect caused by TgVMP1 depletion, indicating their functional conservation and supporting a crucial role of the ER-localized scramblase activity in the biogenesis and function of apicomplexan secretory organelles.

PfVMP1 and TgVMP1 proteins share the DedA superfamily domain organization, are structurally similar to HsVMP1, and localize to ER, indicating that the predicted PfVMP1 and TgVMP1 proteins belong to the DedA superfamily. Although the ER-retention/retrieval signals are yet to be identified for VMP1 proteins, the C-terminal cytoplasmic tails of VMP1 homologs contain one or more “di-Lys” motif or its variants (**Figure S1D**), which might mediate the ER localization of VMP1 proteins, as has been reported for several ER-membrane proteins ^58^.

TgVMP1 depletion decreased parasite motility, host cell invasion, egress, and intracellular development, indicating a crucial role for it in these processes. TgVMP1 depletion caused decreased microneme secretion, impaired rhoptry formation and secretion, and impaired dense granule formation and secretion, which indicated that TgVMP1 is critical for the biogenesis and function of these secretory organelles. As micronemes, rhoptries, and dense granules are essential for the parasite motility, invasion, egress, and intracellular development ^2–10,49^, the phenotypic defects of TgVMP1-depleted parasites could be directly attributed to the impaired biogenesis and functions of these secretory organelles.

TgVMP1 depletion did not affect microneme organization, but decreased microneme secretion. Microneme secretion is induced by a signaling cascade of cyclic guanosine monophosphate (cGMP), cytosolic Ca^2+^, and phosphatidic acid ^59,60^. +IAA RHTIR1-VMP1AID parasites showed elevated Ca^2+^ level as compared with -IAA RHTIR1-VMP1AID parasites **(Figure S12**), which would enhance microneme secretion. On the contrary, TgVMP1-depletion decreased MIC2 secretion, and also impaired MIC2 processing, which has been shown to be crucial for the secretion of MIC2 and several other microneme proteins ^51,61^. This suggests that TgVMP1 depletion adversely affected the MIC2-processing proteases or the processing environment, thereby, decreased microneme secretion. TgVMP1 interactome contained V-type proton ATPase and ASP3, which have been shown to be required for the processing of microneme and rhoptry proteins during their transport through the endosomal compartments^56,57^, suggesting that TgVMP1 regulates microneme secretion via interaction with these proteins.

Rhoptry proteins are synthesized in ER and transported in secretory vesicles arising from the trans-Golgi network (TGN) to precursor structures that mature to rhoptries ^4,62^. Rhoptry discharge process is not clear, except that microneme secretion is one of the triggers ^63,64^. TgVMP1-depleted parasites had none or fragmented rhoptries, which is consistent with decreased rhoptry secretion. TgVMP1 interactome contained TgSORTLR and TgARO, which have been shown to be required for the trafficking of microneme and rhoptry proteins and apical positioning of rhoptries ^54,55^, respectively, suggesting that TgVMP1 contributes to rhoptry biogenesis and secretion through interaction with TgSORTLR and TgARO, and the loss of this interaction in TgVMP1-depleted parasites could have adversely affected rhoptry biogenesis. The decreased rhoptry secretion could be a consequence of decreased microneme secretion.

Dense granules secrete GRA proteins into the PV, which contribute to the formation of IVN within the PV and host cell remodeling ^6,9,10^. Most of the characterized GRA proteins are synthesized in ER, transported by vesicle transport pathway to TGN wherefrom dense granules bud off ^6^. TgVMP1 depletion caused accumulation of dense granules in ER, which would reduce dense granule secretion into the PV, resulting in the loss of IVN. The presence of dense granule proteins (GRA8, GRA12, GRA25, and GRA71) in the TgVMP1 interactome suggests their interaction with TgVMP1, which could facilitate the biogenesis of dense granules.

Lipid scramblase activity of TgVMP1 and PfVMP1 and rescue of the TgVMP1-depleted parasites by PfVMP1 or a known ER-localized lipid scramblase as distant as HsVMP1 indicate that an ER-resident lipid scramblase activity is crucial, which likely functions via translocation of lipids in the ER and the precursor vesicles destined to the secretory organelles. This further indicates that the loss of ER-localized lipid scramblase is the major reason for impaired secretory organelle biogenesis, ER organization, and LD homeostasis in TgVMP1-depleted parasites. Since TgVMP1 depletion did not affect trafficking of the proteins destined to micronemes, parasite plasma membrane, and IMC, TgVMP1 may contribute to the formation of specific ER subdomains wherefrom precursor rhoptry and dense granule vesicles arise. This proposal is in agreement with the reported role of HsVMP1 in the formation of ER microdomains that make contact with diverse organelles ^34^.

Altogether, we demonstrate that TgVMP1 and PfVMP1 are ER-resident lipid scramblases, which are crucial for the biogenesis and function of apicomplexan secretory organelles, most likely via translocation of lipids in the ER wherefrom vesicles destined to the secretory organelles arise **(Figure S13)**. Functional conservation of TgVMP1, PfVMP1, and HsVMP1 hints to a fundamental function of this family of proteins in eukaryotes. Since DedA superfamily proteins are present in all the life forms, it would be exciting to investigate whether lipid scramblase activity is a general feature of this superfamily and how it mediates specialized functions depending on the subcellular location. The essentiality of TgVMP1 for parasite development and likely functional conservation of apicomplexan VMP1 proteins highlight their drug-target potential.

## Materials and Methods

### Materials

All the routinely used biochemicals were of molecular biology or cell culture grade, and were purchased from Thermo Fisher Scientific, Sigma-Aldrich, MP Biomedicals or Serva unless otherwise stated. Cell culture media and supplements, blasticidin S-hydrochloride, gentamicin, penicillin-streptomycin, and Giemsa stain were from Lonza and Thermo Fisher Scientific. Lipids (POPC, POPS, and NBD-PS) were from Avanti Polar Lipids. Dithionite was from Sigma-Aldrich. All the plasticware for cell culture was procured from standard manufacturers, including Thermo Fisher Scientific, Corning Inc, Nalgene, TPP, and Tarsons. Restriction enzymes and DNA modifying enzymes were from New England Biolabs and Thermo Fisher Scientific. DNA and plasmid isolation kits were from QIAGEN and MACHERY-NAGEL. PrimeScript 1^st^ strand cDNA synthesis kit was from Takara Bio. ProLong Diamond Antifade Mountant, DAPI, Hoechst, ER-Tracker Red stain (BODIPY TR Glibenclamide), and SuperSignal Chemiluminescent substrates were from Thermo Fisher Scientific. Trimethoprim, N-acetylglucosamine, indole-3-acetic acid (IAA), thapsigargin, xanthine, mycophenolic acid, calcium ionophore A23187, formaldehyde and Nile red were purchased from Merck. Ionomycin was from Tocris Bioscience. 5Ph-IAA was from MedChemExpress. Pyrimethamine was from MP Biomedicals. Protease inhibitor cocktail was from Roche. Glutaraldehyde, formaldehyde, osmium tetroxide, and sodium cacodylate were from Electron Microscopy Sciences. Uranyl acetate was from Lobachemie and epoxy resin components were from TED Pella. Antibodies were from Thermo Fisher Scientific, Cell Signaling Technology, Santa Cruz Biotechnology or Jackson ImmunoResearch. GFP-Trap and HA-Trap antibodies were from ChromoTek. Pierce protein A/G magnetic beads were from Thermo Fisher Scientific. Human Foreskin Fibroblasts (HFF) were from ATCC, *P. falciparum* 3D7 and D10 strains, *T. gondii* RH TIR1-3FLAG strain, and antibodies to *T. gondii* proteins (AMA1, MIC2, MIC3, ROP1, GRA1, GRA2 and SAG1) were procured from the Biodefense and Emerging Infections Research Resources Repository (BEI Resources). WR99210 was a kind gift from David Jacobus (Jacobus Pharmaceutical, Princeton, U.S.A.). Rabbit anti-IMC1 polyclonal antibodies were a kind gift from Abhijit S. Deshmukh (NIAB, India). pTKO-HXGPRT plasmid was a kind gift from Dr. Nishit Gupta (BITS, India). pTUB1:YFP-mAID-3HA-DHFR-TS:HXGPRT (Addgene#87259) and pSAG1::Cas9-U6::sgUPRT (Addgene #54467) plasmids were a kind gift from Dr. David Sibley. pCTG-Fluc (originally sourced from Dr. Shobhona Sharma, TIFR, India) and pARL1a-PfGRASP-GFP (originally sourced from Dr. Lilach Sheiner, University of Glasgow, UK) were kind gifts from Dr. Swati Patankar (IIT Bombay, India).

Human blood was collected from healthy volunteers after obtaining informed consent according to the approved protocol of the Institutional Ethics Committee (IEC-103/2023, IEC-38/2015, IEC-38-R1/2015, IEC-38-R2/2015 and IEC-38-R3/2015) of Centre for Cellular and Molecular Biology, India. *P. falciparum* and *T. gondii* cultures and related experiments were carried out according to the approved protocols of the Institutional Biosafety Committee of Centre for Cellular and Molecular Biology, India. Animals were housed in cabin type isolators at room temperature (22-25°C, 40–70% humidity, and 12/12 h dark/light photoperiod), and all the experiments on animals were performed according to the approved protocols (IAEC 09/2023) of the Institutional Animal Ethics Committees (IAEC) of Centre for Cellular and Molecular Biology, India. All the studies were carried out as per the Declaration of Helsinki principles.

## Methods

### Sequence analysis

The amino acid sequences of VMP1 proteins of model organisms (*Homo sapiens*, ID: Q96GC9; *Drosophila melanogaster*, ID: Q9W2S1; *Caenorhabditis elegans*, ID: Q9XWU8; *Arabidopsis thaliana*, ID: Q5XF36 (KSM1) and F4I8Q7 (KSM2); *Chlamydomonas reinhardtii*, ID: A0A2K3DNE8; *Dictyostelium discoideum*, ID: Q54NL4; *Danio rerio*, ID: Q6NYY9), *S. cerevisiae* TVP38 (ID: P36164), and selected *E. coli* DedA superfamily proteins (YohD, ID: P33366; YdjZ, ID: P76221; YqaA, ID: P0ADR0) were obtained from the Uniprot database (https://www.uniprot.org/). The amino acid sequences of HsVMP1, ScTVP38, and *E. coli* DedA superfamily proteins (YohD, YdjZ, YqaA) were used as queries in the BLAST searches of the eukaryotic pathogen database VEupathDB (https://veupathdb.org/) to identify homologs in selected reference apicomplexan parasites, including *T. gondii* and *P. falciparum* (**Table S1**). The amino acid sequences of apicomplexan VMP1 proteins were analyzed for the presence of conserved domains using CD BLAST (ncbi.nlm.nih.gov) or MOTIF (genome.jp/tools/motif/), and sequences were aligned using Clustal Omega ^65^.

### Parasite culture

*P. falciparum* 3D7 and D10 strains were maintained in human RBCs (at 2% haematocrit) in RPMI-1640 medium (supplemented with 2 g/l sodium bicarbonate, 2 g/l glucose, 25 mg/ml gentamicin, 300 mg/l L-glutamine, 100 mM hypoxanthine, 0.5% albumax II) under a mixed gas environment (5% CO_2_, 5% O_2_ and 90% N_2_) at 37°C ^66–68^. The synchrony of *P. falciparum* culture was maintained by treating the cells with 5% sorbitol when majority of the parasites were at ring stage ^69^. Genomic DNA was isolated from late trophozoite/schizont stage parasites using the NucleoSpin Tissue kit as recommended by the manufacturer.

Human foreskin fibroblast (HFF) cells were maintained in D10 medium (DMEM supplemented with 10% FBS, 2 mM glutamine, 3.7 g/l sodium bicarbonate, 155 mg/l sodium pyruvate, 1% pen-strep) at 37℃ under 5% CO_2_ environment. *T. gondii* TIR1-3FLAG RH strain (RHTIR1) ^70^ was grown in confluent HFF cells as has been reported previously ^71^. Fully confluent HFF cells infected with RHTIR1 tachyzoites were scraped, and the suspension was passed through a 3 µm filter to remove host debris. The filtrate was centrifuged at 800×g for 8 min, washed with incomplete DMEM medium (DMEM without antibiotics and FBS), and the purified parasite pellet was resuspended in an appropriate buffer as has been described in the following sections.

Genomic DNA (gDNA) was isolated from purified *T. gondii* RHTIR1 tachyzoites using the NucleoSpin Tissue kit following the manufacturer’s instructions. For *T. gondii* cDNA preparation, total RNA was isolated from the purified RHTIR1 *T. gondii* tachyzoites using the NucleoSpin RNA isolation kit according to the manufacturer’s instructions. 5 μg of the DNA-free total RNA was used to prepare cDNA using PrimeScript 1^st^ strand cDNA synthesis kit as per the manufacturer’s instructions.

### Generation of recombinant *P. falciparum* parasites

The primers and synthetic DNAs used for the generation of transfection constructs are listed in **Table S3**. To knock-in GFP immediately downstream of the 3’-PfVMP1coding sequence, the 3’-coding sequence of PfVMP1 corresponding to 249 amino acid residues was amplified from *P. falciparum* 3D7 genomic DNA using PfVMP1-F1/PfVMP1-R1 primers, and cloned into the pGEM-T vector to obtain pGEMT-PfVMP1c plasmid. The pGEMT-PfVMP1c plasmid was digested with ApaI/KpnI to excise the PfVMP1c fragment, which was subcloned into similarly digested pGT-GFP plasmid to obtain pGT-PfVMP1c-GFP. The pGT-PfVMP1c-GFP plasmid was digested with ApaI/XhoI to excise PfVMP1c-GFP insert, which was subcloned into similarly digested pNC vector to obtain pNC-PfVMP1c-GFP transfection plasmid. pNC and pGT-GFP plasmids have been described earlier ^72,73^.

We constructed a dual knock-down plasmid, which would cause *glmS*-mediated mRNA degradation and mutant *E. coli* DHFR (cDD)-mediated protein degradation as has been described previously ^74^. The 3’-coding sequence corresponding to 249 amino acid residues (flank1) of PfVMP1 coding sequence and 1087 bp of PfVMP1 3’-UTR region (flank2) were amplified from *P. falciparum* 3D7 genomic DNA using PfVMP1-fl1F/PfVMP1-R1 and PfVMP1-Fl2F/PfVMP1-Fl2R primer pairs, respectively. The flank1 and flank2 PCR products were cloned into the HBPFA18/cDD_HA_/GlmSAc plasmid at NotI/KpnI and AvrII/KasI sites, respectively, to obtain HB-PfVMP1-cDD_HA_-GlmSAc transfection plasmid. HBPFA18/cDD_HA_/GlmSAc plasmid has been described earlier ^75^.

We also employed auxin-inducible knock-down strategy for PfVMP1. We purchased synthetic XTEN-mAID-2×HA (mAID_HA_) and PcDT5U/Tir1-Flag-2A/HD/PcDT3’U (TIR1-HD) DNAs, which were cloned into the HBPFA18/cDD_HA_/GlmSAc at KpnI/XhoI and AgeI/AvrII sites, respectively, to obtain HB-mAID_HA_/TIR1-HD plasmid. The mAID-2×HA and TIR1-Flag-2A/HD coding sequences are codon-optimized for *P. falciparum*. The above PfVMP1 flank1 and flank2 regions were cloned into the HB-mAID_HA_/TIR1-HD plasmid at NotI/KpnI and AvrII/KasI sites, respectively, to obtain HB-PfVMP1-mAID_HA_/TIR1-HD transfection plasmid.

Ring stage *P. falciparum* D10 (for PfVMP1c-GFP and HB-PfVMP1-cDD_HA_-GlmSAc plasmids) or 3D7 (for HB-PfVMP1-mAID_HA_/TIR1-HD plasmid) parasites were transfected with 100 µg of plasmid DNA by electroporation, and transfected parasites were selected with blasticidin S hydrochloride (at 2 µg/ml for pNC-PfVMP1c-GFP) or WR99210 (at 1 nM for HB-PfVMP1-cDD_HA_-GlmSAc and HB-PfVMP1-mAID_HA_/TIR1-HD) to obtain resistant parasites as has been previously described ^76,77^. Blasticidin resistant parasites were cloned by dilution cloning to obtain clonal populations. The genomic DNA of each of the clonal parasite lines was isolated, assessed by PCR for plasmid integration using locus-specific primer sets (5’-integration: PfVMP1-5con/GFPSEQR1; 3’-integration: PcDT5U-R1/PfVMP1-3con; wild-type: PfVMP1-5con/PfVMP1-3con; positive control: PfA8fl2-F1 and PfA8fl2-R1). WRR99210 resistant parasites were cloned by dilution cloning to obtain clonal lines, genomic DNA was isolated from clonal lines, and assessed for plasmid integration into the target locus by PCR using locus-specific primer sets (HB-PfVMP1-cDD_HA_-GlmSAc-transfected parasites (5’-integration: PfVMP1-5con/PvAc-Con-R; 3’-integration: Hrp2-SeqF/PfVMP1-3con2; wild type-specific: PfVMP1-5con/ PfVMP1-3con2; positive control: PfA8-F1 and PfA8fl2-R1), HB-PfVMP1-mAID_HA_/TIR1-HD-transfected parasites (PfVMP1-5con/PfVMP1-3con2; positive control: PfUCH-F/PfUCH-R). The cloned recombinant parasites with PfVMP1c-GFP (PfVMP1GFP_KI_), PfVMP1/cDD_HA_-GlmS (PfVMP1dKD) or PfVMP1-mAID_HA_/TIR1-HD (PfVMP1AID) locus were used for various experiments.

### Generation of recombinant *T. gondii* parasites

The transfection cassette containing TgVMP1-mAID-3×HA coding sequence and HXGPRT selection cassette (TgVMP1-mAID-3HA/DHFR-TS:HXGPRT) was amplified from the “pTUB1:YFP-mAID-3HA, DHFR-TS:HXGPRT” plasmid using TgVMP1-Fkd2/TgVMP1-RP primers ^78^. The TgVMP1-Fkd2 primer contained 50 bp homology sequence corresponding to the immediately upstream of the TgVMP1 stop codon and overlap with the 5’-mAID-3HA sequence. The TgVMP1-RP primer contained 50 bp homology sequence corresponding to 133 bp downstream of the TgVMP1 stop codon. The primer (SgDNA1) corresponding 20 nucleotide protospacer sequence in the 3’ UTR of TgVMP1 along with SgDNARP primer were used to amplify pSAG1::Cas9-U6::sgUPRT plasmid ^79^, which generate PCR product with the target guide RNA sequence and Cas9 expression cassette (pSAG1::Cas9-U6::sgTgVMP1). The PCR product was transformed into DH5α cells, and sequenced to ensure the correctness of guide RNA coding sequence.

*T. gondii* RH TIR1-3×FLAG (RHTIR1) tachyzoites were transfected with a mixture of TgVMP1-mAID-3HA/DHFR-TS:HXGPRT transfection cassette and pSAG1::Cas9-U6::sgTgVMP1 plasmid as has been described previously ^41^. In brief, fully confluent HFF cells infected with RHTIR1 tachyzoites were scraped, tachyzoites were purified using a 3 µm filter to remove host debris, centrifuged at 800×g for 8 min, and washed in incomplete DMEM medium (DMEM without antibiotics and FBS). 5×10^6^ tachyzoites were mixed with 15 µg each of the transfection cassette and plasmid, the volume was adjusted to 400 µl using incomplete medium, transferred to a 2 mm cuvette, and electroporated (at 1500 V, 10 µF capacitance, ∞ resistance). The electroporated parasites were transferred to confluent HFF cells in T25 flasks, grown in drug-free complete medium for 24 hrs, followed by in selection medium (complete medium with 25 µg/ml mycophenolic acid and 50 µg/ml xanthine) for 48 hrs. The intracellular parasites were syringe lysed and 500 µl of the lysed suspension was added to fully confluent HFF cells in a T25 flask, and grown in the selection medium. Resistant parasites were diluted to achieve 0.5 parasite/well in a 96-well tissue culture plate containing HFF monolayer, and allowed to grow until parasites appeared. These parasites were screened by PCR for the presence of integration locus using TgVMP1-5con/TgVMP1-3con primer set along with the TgVP1-5con/TgVP1-3con primer set for TgVP1 gene as a positive control. Parasites showing the absence of wild-type TgVMP1 and presence of the integration locus were checked for expression of the TgVMP1-mAID-3×HA fusion protein (TgVMP1AID_HA_) by western blotting using anti-HA antibodies. Pure clonal recombinant parasites (RHTIR1-VMP1AID) were used for further studies.

### Expression of ER and Golgi reporters in RHTIR1-VMP1AID parasites

Tubulin (Tub) promoter region was amplified from RHTIR1 gDNA using Tub F/Tub R primers by PCR, and cloned into pMyc3x.LIC-DHFR plasmid at PacI/EcoRV sites to obtain pTub-DHFR plasmid. The N-terminal signal sequence (MTAAKKLSLFSLAALFCLLSVATLRPVAASD) and C-terminal ER-retention signal (HDEL) of *T. gondii* chaperonin BiP (TGRH88_051090) were added to the N-terminal and C-terminal of GFP by PCR to generate the ER reporter BiP-GFP-HDEL. In brief, the BiP N-terminal signal sequence was amplified from *T. gondii* gDNA using BipSig-F/BipSig-GFP-Rrec primers. GFP coding sequence was amplified from pTKO-HXGPRT plasmid using GFP-Bip-Frec/GFP-HDEL-R primers. The two PCR fragments were recombined by PCR using primers BipSig-F/GFP-HDEL-R to obtain BiP-GFP-HDEL (ER-GFP) coding sequence, which was cloned at EcoRV/AvrII sites in pTub-DHFR plasmid to obtain pTub-ER-GFP plasmid. 15 µg of this plasmid was transfected into RHTIR1-VMP1AID parasites to obtain RHTIR1-VMP1AID::ER-GFP parasites as has been described above in “Generation of recombinant *T. gondii* parasites” section.

For the expression of Golgi reporter PfGRASP-GFP, which has been used as a Golgi reporter previously ^42^, 15 µg of the pARL1a-PfGRASP-GFP plasmid was transfected into RHTIR1-VMP1AID tachyzoites to obtain RHTIR1-VMP1AID::GRASP-GFP parasites as has been described in the “Generation of recombinant *T. gondii* parasites” section.

### VMP1 expression

Cloned PfVMP1GFP_KI_ parasites, which would express PfVMP1GFP fusion protein, and wild-type *P. falciparum* D10 parasites were cultured and synchronized as has been described in the parasite culture section. Synchronized PfVMP1GFP_KI_ parasites were harvested at ring, early trophozoite, late trophozoite, and schizont stages, purified by saponin lysis, and parasite pellets were resuspended in 6× pellet volume of 10 mM Tris (pH 7.4). The resuspension was mixed with 1/3^rd^ volume of 4× SDS-PAGE sample buffer (1× buffer has 50 mM Tris-HCl, 10% glycerol (v/v), 2% SDS, 2.5% β-mercaptoethanol (v/v), 0.05% bromophenol blue (w/v), pH 6.8), heated at 99°C for 10 min, centrifuged at 20,000g for 20 min, and supernatants were processed for western blotting. In brief, the lysates were run in 12% SDS-PAGE gel, transferred to an Immobilon-P-PVDF membrane, incubated with blocking buffer (TBS-T (10 mM Tris, 150 mM NaCl, 0.1% Tween-20, pH 7.4) with 3% fat-free milk powder), probed using rabbit anti-GFP monoclonal antibodies, followed by goat anti-rabbit-HRP conjugated secondary antibodies (at 1:20000 dilution). The membrane was washed and the signal was developed using SuperSignal chemiluminescent substrate. For loading control, the membrane was stripped off, blocked, incubated with mouse anti-β-actin-HRP conjugated antibodies (at 1:30000 dilution), and the signal was developed using SuperSignal chemiluminescent substrate.

RHTIR1 and RHTIR1-VMP1AID parasites were cultured in confluent HFF cells for 24 hrs post-invasion as has been described in the parasite culture section. Infected HFF cells were syringe lysed to release parasites, the lysate was passed through a 3 µm filter, and the filtrate containing parasites was centrifuged at 800g. The parasite pellet was resuspended in 6× pellet volume of 10 mM Tris (pH 7.4), the suspension was mixed with 1/3^rd^ volume of 4× SDS-PAGE sample buffer, heated at 99°C for 10 min, centrifuged at 20,000g for 20 min, and supernatants were processed for western blotting as has been described above for PfVMP1GFP_KI_ parasites. The membranes were incubated with rabbit anti-HA monoclonal antibodies (for TgVMP1AID_HA_, 1:1000 dilution), followed by goat anti-rabbit-HRP conjugated secondary antibodies (at 1:20000 dilution). For loading control, the blot was stripped, blocked, and incubated with mouse anti-ROP1 antibodies (at 1:1000 dilution) for 2 hrs, followed by HRP-conjugated goat anti-mouse antibodies.

To assess for TgVMP1 depletion, fully confluent HFF cells in T25 flasks were infected with the RHTIR1-VMP1AID parasites and allowed to grow for 24 hrs. Parasites were subjected to grow in the presence of 500 μM auxin (+IAA) for 0, 15, 30, 60, 120 and 180 min, respectively. Following this, cells were harvested by scraping the parasite-infected HFF cells and subjecting them to centrifugation at 800×g. The obtained pellets were washed with 1×PBS, and centrifuged again at 800×g. The resultant pellets were processed for western blotting analysis as described above. TgVMP1AID_HA_ levels were determined by probing the blots using rabbit anti-HA antibodies, followed by goat anti-rabbit-HRP conjugated secondary antibodies, as described above. Mouse anti-ROP1 and anti-SAG1 antibodies (at 1:1000 dilution), followed by anti-mouse-HRP antibodies (at 1:20,000 dilution) were used to probe ROP1 and SAG1 as the loading controls.

### VMP1 localization

To determine subcellular localization of PfVMP1GFP, mixed stage PfVMP1GFP_KI_ parasites were washed with PBS, stained with Hoechst (1 µg/ml in PBS), washed with PBS, immobilized on a poly-L-lysine coated slides, washed again to remove unattached cells, and covered with a glass cover slip. Live parasites were observed under the 100× objective of Zeiss Axioimager.Z2 fluorescence microscope, images were acquired using AxioVision software, and analysed using ImageJ software. For co-localization, trophozoite stage PfVMP1GFP_KI_ parasites were grown in complete medium containing the ER-Tracker Red stain (200 nM) for 20 min at 37°C, Hoechst (1 µg/ml in complete medium) was added to the culture, and the culture was incubated for 10 min. The cells were washed with PBS, immobilized on a poly-L-lysine coated slide, washed again to remove unattached cells, and covered with a glass cover slip, and images of live cells were captured using the 100× objective of Olympus FV300 confocal laser scanning microscope. The images were analysed for co-localization and Pearson’s correlation coefficient was determined using JACoP plugin in Fiji.

For localization of TgVMP1AID_HA_, RHTIR1-VMP1AID parasites were grown in HFF cells on coverslips in 12 well plates for 38 hrs post-invasion without indole-3-acetic acid (IAA), and processed for immunofluorescence assay (IFA). In brief, the culture medium was aspirated, cells were washed with PBS, fixed with formaldehyde (4% in PBS) for 30 min, washed with PBS, permeabilized using 0.1% Triton-X 100 for 30 min, washed with PBS, and blocked with blocking buffer (3% BSA in PBS) for 1 hr. The cells were incubated with rabbit anti-HA antibodies (at 1:200 dilution) for 2 hrs. The coverslips were washed with PBS, incubated with Alexa Fluor 488-conjugated F(ab’)_2_ donkey anti-rabbit IgG (at 1:1000 dilution) for 1 hr at room temperature, incubated with DAPI (10 µg/ml in PBS) for 10 min, washed with PBS, air dried, and mounted using the ProLong^TM^ diamond antifade mountant. The cells were observed under the 100× objective, and images were acquired on Zeiss LSM 880 confocal laser scanning microscope. Airyscan mode was used in Zeiss LSM 880 microscope to obtain super-resolution micrographs, images were exported using Zen Black software, and processed using ImageJ software. Maximum intensity projections of z-stack images were obtained using z-projection of ImageJ software. Pearson’s correlation coefficient was determined using the JACoP plugin of Fiji software.

For co-localization of TgVMP1AID_HA_ with ER-GFP, RHTIR1-VMP1AID::ER-GFP parasites were grown without IAA, and processed for IFA using mouse anti-HA antibodies (at 1:200 dilution) and rabbit anti-GFP antibodies (at 1:200 dilution), followed by incubation with appropriate secondary antibodies (Alexa Fluor 488-conjugated F(ab’)_2_ donkey anti-rabbit IgG and Alexa Fluor 647-conjugated F(ab’)_2_ donkey anti-mouse IgG) as has been described above for TgVMP1. Images were acquired using Zeiss LSM 880 confocal laser scanning microscope. Airyscan mode was used in Zeiss LSM 880 microscope to obtain super-resolution micrographs, images were exported using Zen Black software, and processed using ImageJ software. Pearson’s correlation coefficient was determined using JACoP plugin in Fiji. For co-localization of TgVMP1AID_HA_ with GRASP-GFP, RHTIR1-VMP1AID::GRASP-GFP parasites were grown without IAA, and processed for IFA using mouse anti-HA (at 1:200 dilution) and rabbit anti-GFP antibodies (at 1:200 dilution) antibodies, followed by incubation with appropriate secondary antibodies (Alexa Fluor 488-conjugated F(ab’)_2_ donkey anti-rabbit IgG and Alexa Fluor 647-conjugated F(ab’)_2_ donkey anti-mouse IgG) as has been described above for TgVMP1. The cells were observed under the 100× objective of Olympus FV3000 confocal laser scanning microscope, images were captured using Fluoview FV31S-SW software, and processed as has been described above for TgVMP1. Pearson’s correlation coefficient was determined using JACoP plugin in Fiji.

For TgVMP1 localization under IAA-induced depletion condition, RHTIR1-VMP1AID parasites were grown in HFF cells on coverslips in 12 well plates for 24 hrs post-invasion, followed by growth for additional 14 hrs in the presence of 0.1% ethanol (-IAA) or 500 µM (+IAA) indole-3-acetic acid, and processed for IFA as has been described for TgVMP1. Infected HFF cells were processed for IFA using rabbit anti-HA (at 1:200 dilution) and mouse anti-SAG1 (at 1:500 dilution) antibodies for 2 hrs, followed by incubation with appropriate secondary antibodies (Alexa Fluor 488-conjugated goat anti-rabbit and Alexa Fluor 647-conjugated goat anti-mouse antibodies) as has been described above for TgVMP1. The cells were observed under the 100× objective, and images were acquired using Airyscan mode on Zeiss LSM 880 confocal laser scanning microscope, exported using Zen Black software, and processed using ImageJ software. Maximum intensity projections of the z-stacks were obtained using z-projection of ImageJ software.

To study the effect of TgVMP1 depletion on the inner membrane complex, RHTIR1-VMP1AID parasites were grown with (+IAA) or without (-IAA) indole-3-acetic acid, and processed for IFA using rabbit anti-IMC1 antibodies (at 1:500 dilution) and mouse anti-HA (at 1:200 dilution) antibodies, followed by appropriate secondary antibodies (Alexa Fluor 488-conjugated F(ab’)2 donkey anti-rabbit IgG and Alexa Fluor 647-conjugated F(ab’)2 donkey anti-mouse IgG (1:1000 dilution)) as has been described above for TgVMP1. The cells were observed under the 100× objective of Zeiss LSM 880 confocal laser scanning microscope, images were acquired using Airyscan mode, exported using Zen Black software, and processed using ImageJ software. Maximum intensity projections of the z-stacks were obtained using z-projection option in ImageJ software.

To study the effect of TgVMP1 depletion on ER, the RHTIR1-VMP1AID::ER-GFP parasites were grown in HFF cells on coverslips in 12 well plates for 24 hrs post-invasion, and then grown for additional 14 hrs with (+IAA) or without (-IAA) IAA. The intracellular or purified parasites were fixed in formaldehyde (4%) and processed for IFA using rabbit anti-GFP (at 1:200 dilution) and mouse anti-HA (at 1:200 dilution) antibodies, followed by appropriate secondary antibodies (Alexa Fluor 488-conjugated goat anti-rabbit or Alexa Fluor 647-conjugated goat anti-mouse antibodies) as has been described above for TgVMP1. The cells were observed under the 100× objective of Olympus FV3000 confocal laser scanning microscope, and images were acquired using Fluoview FV31S-SW software, and processed using ImageJ software.

### Effect of PfVMP1 depletion on parasite growth

Wild-type *P. falciparum* D10 and 3D7 strains, PfVMP1dKD, and PfVMP1AID parasites were cultured and synchronized as has been described in the parasite culture section. PfVMP1dKD and PfVMP1AID parasites were assessed for PfVMP1 depletion and its effect on parasite growth. To check conditional PfVMP1 depletion, ring stage PfVMP1dKD parasites were grown at 8-10% parasitemia under control (+TMP-GlcN), single knock-down (-TMP-GlcN, +TMP+GlcN) or dual knock-down (-TMP+GlcN) conditions for two consecutive cycles (96 hrs) as has been described in the “Parasite culture” section. Parasites (at 8-10% parasitemia) were harvested at 48 and 96 hrs time points, and parasite lysates were processed for western blotting using rabbit anti-HA antibodies (1:1000 dilution), followed by HRP-conjugated goat anti-rabbit antibodies (1:20000 dilution) as has been described in “VMP1 expression” section. For loading control, the membrane was stripped off and probed with mouse HRP-conjugated anti-β-actin-antibodies (at 1:30,000 dilution). To assess the effect of PfVMP1 deletion on parasite growth, synchronized ring stage PfVMP1dKD parasites were grown under control (+TMP-GlcN), single knock-down (-TMP-GlcN; +TMP+GlcN) or dual knock-down (-TMP+GlcN) conditions for 4 consecutive cycles (192 hrs). The number of parasite-infected RBCs was determined by FACS using BD Fortessa FACS analyser as has been described earlier ^80^, data was presented as % parasitemia over 4 cycles using GraphPad Prism.

Likewise, PfVMP1AID parasites were assessed for 5Ph-IAA-inducible PfVMP1 depletion and the effect of depletion on parasite growth. Briefly, three different clonal lines of the ring stage PfVMP1AID parasites were cultured in the presence or absence of 1 μM 5Ph-IAA for two consecutive erythrocytic cycles as has been described in the “Parasite culture” section. Parasites (at 8-10% parasitemia) were harvested at 24, 48, 72, and 96 hrs time points, the parasite lysates were processed for western blotting using rabbit anti-HA antibodies (1:1000 dilution), followed by HRP-conjugated goat anti-rabbit antibodies (1:20000 dilution) as has been described in “VMP1 expression” section. The membrane was stripped off and probed for β-actin as a loading control using mouse HRP-conjugated anti-β-actin-antibodies (at 1:30,000 dilution). For assessing the effect of PfVMP1 depletion on parasite growth, three different clonal lines of the ring stage PfVMP1AID parasites were cultured in the presence (+A) or absence (-A) of 1 μM 5Ph-IAA for two consecutive erythrocytic cycles as has been described in the “Parasite culture” section. Parasitemia was maintained at 3-4%, Giemsa-stained smears were made at the end of each cycle, observed under the 100× objective of a light microscope, and at least 1000 RBCs were counted to determine the percent parasitemia. Data was presented as % parasitemia over three cycles using the GraphPad Prism.

### Effect of TgVMP1 depletion on parasite growth

To assess the effect of TgVMP1 depletion at protein level, we compared RHTIR1 and RHTIR1-VMP1AID parasites for the overall development of by plaque assay and intracellular development by replication assay essentially as has been previously described ^81,82^. For plaque assay, fully confluent HFF monolayers in 6-well plates were infected with 500 freshly egressed RHTIR1 or RHTIR1-VMP1AID parasites per well, grown in the presence of 0.1% ethanol (-IAA) or 500 µM (+IAA) indole-3-acetic acid for 8 days at 37⁰C. The cells were fixed with methanol for 5 min at room temperature, stained with Giemsa for 10 min, and washed with distilled water. The plates were air dried, observed for plaques, and images were acquired using the ChemiDoc Imaging System (Bio-Rad). The area of each of the 100 plaques from 3 independent experiments was measured using ImageJ software, and presented as mean plaque area (mm^2^) using the GraphPad Prism software.

For replication assay, approximately 10^6^ freshly purified RHTIR1 or RHTIR1-VMP1AID parasites were added to fully confluent HFF cells grown on coverslips in 12-well plates. After 3 hrs of invasion, extracellular parasites were removed by washing with PBS, infected cells were grown for 24 hrs in the presence of 0.1% ethanol (-IAA) or 500 µM (+IAA) IAA. The culture media was carefully aspirated, cells were fixed with formaldehyde (4% in PBS), and processed for IFA using mouse anti-GRA1 antibodies (1:500), followed by Alexa Fluor 647-conjugated F(ab’)2 donkey anti-mouse IgG antibodies and DAPI (10 µg/ml) staining as has been described for TgVMP1 in the “VMP1 localization” section. The number of parasites/PV was determined by observing at least 100 random PVs from 3 independent experiments, and presented as the number of PVs containing different number of parasites as a percentage of the total number of PVs.

### Effect of TgVMP1 depletion on in vivo development

Female BALB/cJ mice were infected with RHTIR1-VMP1AID parasites to determine the effect of TgVMP1 depletion on in vivo development as has been described previously ^83^. RHTIR1-VMP1AID parasites were cultured without IAA for 48 hrs post-infection, and purified as has been described in the “Parasite culture” section. 8-10 weeks old 20 female BALB/cJ mice were infected intraperitoneally with purified RHTIR1-VMP1AID tachyzoites (200 parasites/mouse), and five age-matched mice were used as uninfected naïve control. The infected mice were divided into 2 groups (10 mice/group). Mice in one infected group were given 0.2 ml water with 12.5 mg/ml indole-3-acetic acid (+IAA) by oral gavage every day, and also provided drinking water with IAA (0.5 mg/ml) and sucrose (5% (w/v)). Another infected group was given 0.2 ml water only by oral gavage every day and also provided drinking water with (5% sucrose (w/v)). The naïve control group was given 0.2 ml water with 12.5 mg/ml IAA by oral gavage every day and also provided drinking water with IAA (0.5 mg/ml) and sucrose (5% (w/v)). Drinking water was changed every 2-3 days, the mice were monitored for 21 days. Peritoneal exudates of mice were collected on the day of their death and examined for the presence of parasites using mouse anti-SAG1 and TgVMP1AID_HA_ using rabbit anti-HA antibodies by IFA as has been described in the “VMP1 localization” section.

### Effect of TgVMP1 depletion on gliding motility

To assess the effect of TgVMP1 depletion on gliding motility of tachyzoites, we compared the motility of RHTIR1-VMP1AID parasites cultured with or without IAA as has been previously described ^84^. In brief, RHTIR1-VMP1AID parasites were added to confluent HFF cells in T75 flask, allowed to invade the cells for 3 hrs, and extracellular parasites were removed by washing the cells with D10 medium. The cells were grown for 24 hrs, followed by additional 14 hrs in the presence of 0.1% ethanol (-IAA) or 500 µM (+IAA) indole-3-acetic acid, and parasites were purified as has been described in the “Parasite culture” section. 2×10^6^ purified tachyzoites were resuspended in HBSS, allowed to glide on FBS-coated coverslips in a 12-well plate for 15 min, fixed with formaldehyde (4% in PBS), and processed for IFA using mouse anti-SAG1 antibodies (at 1/500 dilution), followed by Alexa Fluor 488-conjugated F(ab’)2 donkey anti-mouse IgG antibodies as has been described for TgVMP1 in the “VMP1 localization” section. Parasites were imaged for gliding trails under 100× objective of the Axioimager.Z2 using AxioVision software, and analysed using ImageJ software. At least 100 parasites were scored for the presence of gliding trails in each of the three independent experiments, and the number of parasites showing gliding trails was presented as a percentage of the total number of parasites observed. At least 90 motile parasites in each of the three experiments were observed for measuring the gliding trail length, and presented as mean trail length (μm).

### Effect of TgVMP1 depletion on host cell invasion

The invasion efficiency of RHTIR1-VMP1AID parasites was determined according to the previously reported two-colour invasion assay ^85^. RHTIR1-VMP1AID parasites were cultured with or without IAA as has been described in the “Effect of TgVMP1 depletion on gliding motility” section. Purified tachyzoites were added to confluent HFF cells grown on coverslips in 12-well plates (MOI 2:1), and incubated for 3 hrs with 0.1% ethanol (–IAA) or 500 µM (+IAA) indole-3-acetic acid. The cells were washed with PBS, fixed with formaldehyde (4% in PBS), and processed for IFA as has been described for TgVMP1 in the “VMP1 localization” section. In brief, the cells were first stained for extracellular parasites with mouse anti-SAG1 antibodies, followed by Alexa Fluor 647-conjugated F(ab’)2 donkey anti-mouse IgG antibodies (at 1:1000 dilution). The cells were washed with PBS, permeabilized with 0.1% Triton-X100, washed with PBS, blocked, incubated again with mouse anti-SAG1 antibodies to stain intracellular parasites, followed by Alexa Fluor 488-conjugated F(ab’)2 donkey anti-mouse IgG antibodies. The cells were observed for red/green (failed to invade) and green only (successful invasion) parasites under the 100× objective of Zeiss Axioimager.Z2 fluorescence microscope, and images were acquired using AxioVision software. The number of red and green parasites was determined, and the number of green parasites was presented as a percentage of the total number of parasites (% Invasion) observed. The experiment was repeated three times and at least 100 parasites were observed in each experiment.

### Effect of TgVMP1 depletion on egress

HFF cells were grown on coverslips in 12-well plates, infected with purified RHTIR1-VMP1AID parasites (at 1:1 MOI), and grown for 48 hrs, followed by additional 14 hrs with 0.1% ethanol (-IAA) or 500 µM (+IAA) indole-3-acetic acid. The cells were washed with PBS, treated with HBSS containing 2 µM ionomycin, a calcium ionophore that induces rapid *T. gondii* egress ^86^. HBSS was gently removed from the wells, the cells were fixed with 4% formaldehyde (in PBS), processed for IFA using mouse anti-GRA1 antibodies, followed by Alexa Fluor 647-F(ab’)_2_ donkey anti-mouse IgG and DAPI as described in the “VMP1 localization” section. The cells were observed under 100× objective of the Zeiss Axioimager.Z2 microscope, images were acquired using AxioVision software, and the total number of sites with well dispersed parasites or clumps of parasites, which mark the PVs from which parasites egressed was determined by counting at least 100 vacuoles in each of the three independent experiments. The number of PVs with parasite clumps was presented as a percentage of the total number of PVs observed using the GraphPad Prism.

### Conoid extrusion assay

RHTIR1-VMP1AID parasites were cultured with 0.1% ethanol (-IAA) or 500 µM indole-3-acetic acid (+IAA) as has been described in the “Effect of TgVMP1 depletion on gliding motility” section, and evaluated for conoid extrusion using the calcium ionophore A23187 as has been previously described ^87^. Briefly, ∼2×10^6^ purified tachyzoites were washed with PBS, resuspended in HBSS, and allowed to settle on coverslips in a 12-well plate for 10 min. The parasites were treated with 2 µM A23187 (in HBSS) for 10 min, fixed with 4% formaldehyde (in PBS), observed under 100× objective of the Axioimager.Z2 microscope, and images were acquired using AxioVision software, and analysed using ImageJ software. At least 100 parasites were observed for conoid extrusion in each of the three independent experiments, and the number of parasites showing conoid extrusion was presented as a percentage of the total number of parasites observed using GraphPad Prism.

### Effect of TgVMP1 depletion on microneme organization and secretion

RHTIR1-VMP1AID parasites were cultured with 0.1% ethanol (-IAA) or 500 µM indole-3-acetic acid (+IAA) as has been described in the “Effect of TgVMP1 depletion on gliding motility” section, and assessed for microneme organization and secretion using antibodies to microneme proteins. For microneme organization, parasite infected cells were fixed with 4% formaldehyde (in PBS), and processed for IFA as has been described in “VMP1 localization” section. The cells were incubated with antibodies to microneme proteins (rabbit anti-AMA1 at 1:500 dilution; mouse anti-MIC2 at 1:500 dilution; mouse anti-MIC3 at 1:500 dilution) and rabbit or mouse anti-HA antibodies (for TgVMPAID_HA_ at 1:200 dilution), followed by appropriate secondary antibodies (Alexa 488- or Alexa 647-conjugated F(ab’)2 donkey anti-mouse or anti-rabbit IgG) and DAPI (10 µg/ml). The cells were observed under 100× objective of the Zeiss LSM 880 confocal laser scanning microscope, and images were acquired in Airyscan mode using Zen Black software, and processed using ImageJ software. Maximum intensity projections of z-stack images were obtained using Z projection option of ImageJ software.

Purified tachyzoites were assessed for microneme secretion according to the previously reported method ^88^. Briefly, ∼3×10^7^ purified tachyzoites were resuspended in 100 µl of DGH medium (DMEM with 2 mM glutamine and 10 mM HEPES), treated with ethanol (1% v/v) at 37⁰C for 2 min to induce secretion, incubated on ice to block further secretion, and centrifuged at 1,000×g for 8 min. The supernatant containing the extracellular secreted antigens (ESA) and the pellet were separated, each fraction was mixed with equal volume of SDS-PAGE sample buffer, and assessed for MIC2 level by western blotting using mouse anti-MIC2 (at 1:2000 dilution) and anti-ROP1 antibodies as has been described in the “VMP1 expression” section. The signal intensities of ROP1 and MIC2 in pellet fractions were measured using ImageJ, the signal intensity of MIC2 was normalized as times of the ROP1 signal intensity for the corresponding parasites, and plotted using GraphPad Prism. The signal intensities of MIC2 in ESA fractions were measured, normalized as times of the MIC2 signal intensity in the corresponding pellet fraction, and plotted using GraphPad Prism. The experiment was repeated four times.

### Effect of TgVMP1 depletion on rhoptry organization

RHTIR1-VMP1AID parasites were cultured with 0.1% ethanol (-IAA) or 500 µM indole-3-acetic acid (+IAA) as has been described in the “Effect of TgVMP1 depletion on gliding motility” section, and assessed for rhoptry organization by IFA using anti-ROP1 antibodies as has been described in the “VMP1 localization” section. The infected cells were incubated with mouse anti-ROP1 (at 1:500 dilution) and rabbit anti-HA antibodies (for TgVMPAID_HA_, at 1:200 dilution), followed by incubation with appropriate secondary antibodies (Alexa Fluor 488-conjugated F(ab’)2 donkey anti-rabbit IgG, Alexa Fluor 647-conjugated F(ab’)2 donkey anti-mouse IgG) and DAPI (10 µg/ml). The cells were observed under the 100× objective of the Zeiss LSM 880 confocal laser scanning microscope, and images were acquired in Airyscan mode using Zen Black software, and processed using ImageJ software. Maximum intensity projections of z-stack images were obtained using Z-projection of ImageJ software.

For co-localization of rhoptries with the ER-GFP, RHTIR1-VMP1AID::ER-GFP parasites were cultured with 0.1% ethanol (-IAA) or 500 µM indole-3-acetic acid (+IAA) as has been described in the “Effect of TgVMP1 depletion on gliding motility” section, tachyzoites were purified, and processed for IFA as described in the “VMP1 localization” section using mouse anti-ROP1 (at 1:500 dilution) and rabbit anti-GFP antibodies (for ER-GFP, at 1:200 dilution), followed by incubation with appropriate secondary antibodies (Alexa Fluor 488-conjugated F(ab’)2 donkey anti-rabbit IgG, Alexa Fluor 647-conjugated F(ab’)2 donkey anti-mouse IgG) and DAPI (10 µg/ml). The cells were observed under 100× objective of the Zeiss LSM 880 confocal laser scanning microscope, images were acquired in Airyscan mode using Zen Black software, and processed using ImageJ software.

### Effect of TgVMP1 depletion on rhoptry secretion

Rhoptries discharge their contents as vacuoles (e-vacuoles) into the host cell upon attachment with the host cell during invasion, which can be monitored using ROP1 as a marker as has been previously described ^89^. Briefly, RHTIR1-VMP1AID parasites were cultured with 0.1% ethanol (-IAA) or 500 µM indole-3-acetic acid (+IAA) as has been described in the “Effect of TgVMP1 depletion on gliding motility” section. 10^7^ purified tachyzoites were treated with 1 µM cytochalasin D (CytD) for 10 min, which prevents invasion, but not rhoptry discharge ^90^. 5×10^6^ CytD-treated parasites were added to confluent HFF cells grown on coverslips in 12-well plates, incubated at 37℃ for 15 min, washed with PBS, fixed with formaldehyde (4% in PBS), and processed for IFA using mouse anti-ROP1 antibodies, followed by incubation with Alexa Fluor 647-conjugated F(ab’)2 donkey anti-mouse IgG and DAPI (10 µg/ml) as has been described in the “VMP1 localization” section. The cells were observed under 100× objective of the Zeiss Axioimager. Z2 microscope, and at least 100 parasites were observed for the presence of e-vacuoles. The experiment was repeated three times, and data was presented as the number of e-vacuole-positive parasites as a percentage of the total parasites using GraphPad Prism.

### Effect of TgVMP1 depletion on IVN formation and dense granule biogenesis

RHTIR1-VMP1AID parasites were cultured with 0.1% ethanol (-IAA) or 500 µM indole-3-acetic acid (+IAA) as has been described in the “Effect of TgVMP1 depletion on gliding motility” section. For IVN formation, intracellular parasites were evaluated for localization of dense granule proteins GRA1 and GRA2 in the PV by IFA as has been described in the “VMP1 localization” section. In brief, intracellular parasites were incubated with mouse anti-GRA1 (at 1:500 dilution) or mouse anti-GRA2 (at 1:500 dilution) and rabbit anti-HA antibodies (for TgVMPAID_HA_, at 1:200), followed by incubation with appropriate secondary antibodies (Alexa Fluor 488-conjugated F(ab’)2 donkey anti-rabbit IgG and Alexa Fluor 647-conjugated F(ab’)2 donkey anti-mouse IgG) and DAPI (10 µg/ml). The cells were observed for PV organization under 100× objective of the Zeiss LSM 880 confocal laser scanning microscope, images were acquired in Airyscan mode using Zen Black software, and processed using ImageJ software. Maximum intensity projections of z-stack images were obtained using z-projection of ImageJ software. ≥50 PVs were examined for the presence of GRA1- or GRA2-labelled IVN (tubulo-vesicular structures), and the experiment was repeated three times. The number of PVs with IVN was presented as a percentage of total PVs observed using GraphPad Prism.

To determine biogenesis of dense granules, purified -IAA and +IAA tachyzoites were added on poly-L-lysine-coated slides, and processed for IFA using mouse anti-GRA1 (at 1:500 dilution) or mouse anti-GRA2 (at 1:500 dilution) antibodies, followed by incubation with appropriate secondary antibodies (Alexa Fluor 488-conjugated F(ab’)2 donkey anti-mouse IgG) and DAPI (10 µg/ml). The cells were observed under the 100× objective of the Olympus FV3000 confocal laser scanning microscope, and images were captured using Fluoview FV31S-SW software. For super resolution images, cells were observed under the 100× objective of the Zeiss LSM 880 confocal laser scanning microscope, and images were acquired in Airyscan mode using Zen Black software. Images were processed using ImageJ software. The fluorescence signal intensity of GRA1 or GRA2 in each tachyzoite was measured using “multi measure” option of the ROI manager, and calculated using the formula “IntDen-(Area of interest × mean signal intensity of the background)”. The mean signal intensity of +IAA parasites was plotted as a percentage of the mean signal intensity of -IAA parasites (% Signal intensity) based on multiple -IAA (n=338) and +IAA (n=383) parasites from three independent experiments. The cells were also analyzed for intraparasitic distribution of GRA1 or GRA2 signal, and the number of parasites with perinuclear accumulation of GRA1 or GRA2 signal was presented as a percentage of the total number of parasites observed.

For co-localization of dense granules with ER-GFP, RHTIR1-VMP1AID::ER-GFP parasites were cultured with 0.1% ethanol (-IAA) or 500 µM indole-3-acetic acid (+IAA) as has been described in the “Effect of TgVMP1 depletion on gliding motility” section, tachyzoites were purified, and processed for IFA as described in the “VMP1 localization” section using mouse anti-GRA1 (at 1:500 dilution) or mouse anti-GRA2 (at 1:500 dilution) and rabbit anti-GFP antibodies (for ER-GFP, at 1:200 dilution), followed by incubation with appropriate secondary antibodies (Alexa Fluor 488-conjugated F(ab’)2 donkey anti-rabbit IgG and Alexa Fluor 647-conjugated F(ab’)2 donkey anti-mouse IgG) and DAPI (10 µg/ml). The cells were observed under 100× objective of the Zeiss LSM 880 confocal laser scanning microscope, images were acquired in Airyscan mode using Zen Black software, and processed using ImageJ software.

### Effect of TgVMP1 depletion on lipid droplets

RHTIR1-VMP1AID parasites were cultured with 0.1% ethanol (-IAA) or 500 µM indole-3-acetic acid (+IAA) as has been described in the “Effect of TgVMP1 depletion on gliding motility” section, tachyzoites were purified, stained with Nile red (250 ng/ml in complete DMEM) at 37⁰C for 30 min, washed with PBS, and observed for fluorescence (λex = 488 nm, λem = 530 nm) under 100× objective of the Zeiss LSM880 confocal laser scanning microscope, images were acquired using Zen Black software, and processed using ImageJ software. The number of lipid droplets (LDs) was determined in multiple parasites (-IAA, n = 137; +IAA, n = 140) from two independent experiments and presented as the number of LDs/parasite. We also measured the surface area (µm^2^) of each LD in multiple parasites (-IAA, n = 473; +IAA, n = 232) from two experiments using the analyze particles option of ImageJ software, and data was presented as mean LD size (μm^2^) using GraphPad Prism.

### Immunoprecipitation and mass spectrometry

For immunoprecipitation of TgVMP1AID_HA_, RHTIR1 and RHTIR1-VMP1AID parasites were cultured without IAA for 48 hrs post-invasion as has been described in the “Parasite culture” section. The purified tachyzoite pellets were resuspended in 300 µl of lysis buffer (10 mM Tris, 150 mM NaCl, 5 mM EDTA, 0.1% NP-40, 10% glycerol, 1× protease inhibitor cocktail, pH 7.5), incubated for 30 min on ice with intermittent mixing, sonicated using Bioruptor for 5 min (30 sec On/30 Sec Off at high amplitude), and the lysates were incubated for 30 min in ice. The lysates were centrifuged at 4°C for 15 min at 20,000g, and the supernatants were collected. The pellets were again extracted with 150 µl of lysis buffer as described above, and supernatants were pooled with the respective first supernatant. The supernatant was incubated with 5 µg of rabbit anti-HA antibodies for 12 hrs at 4°C, mixed with protein A/G magnetic beads (10 µl slurry/2 mg of protein) for 2 hrs at 4°C, and washed with lysis buffer. The beads were washed with lysis buffer, mixed with equal volume of 2× SDS PAGE sample, incubated at 99°C for 10 min, and supernatant was collected as an eluate. Aliquots of input, flow through, washes and eluate were processed for western blotting using mouse anti-HA antibodies (1:1000 dilution), followed by goat HRP-conjugated anti-mouse antibodies (1:20,000 dilution), as has been described in the “VMP1 expression” section. The remaining eluate was subjected to mass spectrometry analysis.

The immunoprecipitates were run on a 12% SDS-PAGE gel until the protein size ladder completely entered the gel. The protein-containing region was excised, and processed for trypsin digestion and peptide extraction as has been described previously ^74,75^. The peptide solution was run on Q Executive HF (Thermo Scientific) for HCD mode fragmentation and LC-MS/MS analysis. The raw data were imported into the Proteome Discoverer 1.2 software (Thermo Scientific), and proteins were identified by searching against the *T. gondii* RH-88 Uniprot databases using the SEQUEST HT algorithm. The analysis parameters included trypsin enzyme specificity, a maximum of two missed cleavages, variable modifications, such as carbamidomethylation of cysteine, oxidation of methionine, and deamidation of asparagine/glutamine. Precursor tolerance was set to 5 ppm, and fragmentation tolerance was set to 0.05 Da. Peptide spectral matches (PSM) and peptide identifications grouped into proteins were validated using the Percolator algorithm and filtered to a 1% false discovery rate (FDR). The proteins identified in RHTIR1 immunoprecipitates were excluded from the TgVMP1AID_HA_ immunoprecipitates, and the remaining proteins were considered as the TgVMP1 interactome.

### Scramblase assay

TgVMP1AID_HA_ and PfVMP1AID_HA_ proteins were purified from RHTIR1-VMP1AID and PfVMP1AID parasites, respectively, as described in the “Immunoprecipitation and mass spectrometry” section. Briefly, the parasite lysate supernatants were incubated with HA-Trap magnetic agarose beads (ChromoTek; 25 µl beads for 2 mg/ml protein) for 12 hrs at 4°C. The beads were washed (wash buffer: 20 mM Tris, 500 mM NaCl, 0.03% n-dodecyl β-D-maltoside (DDM), pH 7.4), and the bound proteins were eluted in 150 µl elution buffer (25 mM Tris, 150 mM NaCl, pH 7.4) with HA peptide (0.64 mg/ml). The eluate was diluted ∼100 fold with storage buffer (20 mM Tris, 150 mM NaCl, 0.03% DDM, 5% glycerol and 1× protease inhibitor cocktail, pH 7.4), concentrated to ∼50 µl using Amicon Ultra centrifugal filter (10 kDa cut-off) at 4°C, flash frozen in liquid nitrogen, and stored at −80°C until further use. Aliquots of input, flow through, washes, and eluate fractions were checked for the presence of TgVMP1AID_HA_ or PfVMP1AID_HA_ proteins by western blotting using rabbit anti-HA antibodies (at 1:1000 dilution), followed by goat HRP-conjugated anti-rabbit antibodies (1:20,000) as has been described in the “VMP1 expression” section.

Liposomes were prepared as has been reported previously ^22,91^. POPC, POPS, and NBD-PS were dissolved in chloroform. POPC and POPS were mixed at 9:1 molar ratio along with 0.4 mol% NBD-PS. The lipid mixture was dried in a glass bottle under nitrogen stream to form a uniform layer, followed by vacuum dried at room temperature for 12 hrs. The dried lipid layer was rehydrated using rehydration buffer (50 mM HEPES, 100 mM NaCl, pH 7.5) to obtain multilamellar vesicles (MLVs) at a final concentration of 10.5 mM. MLVs were subjected to 10 rounds of freeze-thaw cycles, and diluted in rehydration buffer to 5.25 mM. The sample was passed 21 times through a 0.4 µm filter, followed by 11 times through a 0.2 µm filter using a mini extruder (Avanti Polar Lipids) to obtain uniform size liposomes. The liposomes were stored at 4°C until further use.

Scramblase assay was performed as has been described previously ^22,91^. Liposomes (5.25 mM) were destabilized by mixing with DDM (0.36% w/v final), the mixture was incubated at room temperature for 3 hrs at 10 rpm on a nutator. The destabilized liposomes were mixed with storage buffer (with 0.36% DDM) or purified proteins (TgVMP1AID_HA_ or PfVMP1AID_HA_ in storage buffer with 0.36% DDM) at 2:1 ratio (v/v) to prepare empty-liposomes or proteoliposomes (TgVMP1-liposomes or PfVMP1-liposomes), respectively. The samples were incubated at room temperature for 1.5 hrs to allow protein incorporation into the liposomes. DDM was removed by incubating the samples serially with 25 mg Bio-Beads for 1 hr at room temperature, 25 mg Bio-Beads for 2 hrs at room temperature, and 50 mg Bio-Beads for overnight at 4°C. The liquid phase containing liposomes was separated and diluted to 200 µM using rehydration buffer. 50 µl of the empty or proteoliposome sample was transferred to a 96 well flat bottom white polystyrene plate, and assessed for fluorescence signal (λex = 470 nm, λem = 530 nm) using the Spectra Max iD3 multi-mode microplate reader (Molecular Devices). The fluorescence signal intensity was measured at every 26 sec interval for 208 sec, followed by with dithionite (10 mM final concentration) for 416 sec, and then with Triton X-100 (0.5% (w/v)) for 208 sec. Relative fluorescence intensities of empty and proteoliposomes were calculated for each time point, and plotted as (F_t_-F_Triton_)/(F_0_-F_Triton_) where F_t_ is the fluorescence intensity at the corresponding time point, F_0_ is the initial fluorescence intensity, and F_Triton_ is the fluorescence intensity after addition of Triton X-100 (at the 208 sec time point). The relative fluorescence intensity was plotted on y-axis over time on x-axis using GraphPad Prism.

### Expression of PfVMP1 and HsVMP1 in RHTIR1-VMP1AID parasites

A synthetic and mammalian-codon optimized PfVMP1 coding sequence (PfVMP1syn) was amplified from pUC-PfVMP1syn plasmid using PfVMP1-Frec/PfVMP1-R primers. The *T. gondii* tubulin promoter region was amplified from pCTG-Fluc plasmid using TgTub-F/TgTub-PfVMP1-Rrec primers ^92^. The PfVMP1-Frec and TgTub-PfVMP1-Rrec primers contain overlapping sequence, which allowed recombination of PfVMP1syn and tubulin promoter fragments by PCR using TgTub-F/PfVMP1-R primers. The recombined PCR product was digested with PacI/AvrII and cloned into similarly digested pMyc3×.LIC-DHFR plasmid to obtain pTub-PfVMP1syn-3×Myc-DHFR plasmid. A synthetic and *T. gondii*-codon-optimized HsVMP1 coding sequence (HsVMP1syn) was cloned into pJET1.2/Blunt vector, excised with EcoRV/AvrII, and subcloned into similarly digested pTub-PfVMP1syn-3×Myc-DHFR plasmid to obtain pTub-HsVMP1syn-3×Myc-DHFR plasmids. 15 µg of pTub-PfVMP1syn-3×Myc-DHFR or pTub-HsVMP1syn-3×Myc-DHFR plasmid was transfected into RHTIR1-VMP1AID parasites as has been described in the “Generation of recombinant *T. gondii* parasites” section. The transfected parasites were selected with 1 µM pyrimethamine without IAA until the resistant parasites emerged. The resistant parasites with pTub-PfVMP1syn-3×Myc-DHFR plasmid (RHTIR1-TgVMP1AID::PfVMP1_Myc_) would express PfVMP1_Myc_, and the resistant parasites with pTub-HsVMP1syn-3×Myc-DHFR plasmid would express HsVMP1_Myc_. These parasites were cultured with 0.1% ethanol (-IAA) or 500 µM indole-3-acetic acid (+IAA). The parasite lysates were evaluated for expression of TgVMP1AID_HA_ (rabbit anti-HA antibodies), PfVMP1_Myc_ or HsVMP1_Myc_ (mouse monoclonal anti-Myc antibodies at 1: 1000 dilution), and SAG1 (mouse anti-SAG1 antibodies) as a loading control by western blotting as has been described in the “VMP1 expression” section. The RHTIR1-TgVMP1AID::PfVMP1_Myc_ and RHTIR1-TgVMP1AID::HsVMP1_Myc_ parasites were assessed for co-localization of TgVMP1AID_HA_ with PfVMP1_Myc_ or HsVMP1_Myc_, respectively, by IFA as has been described in the “VMP1 localization” section. The parasites were incubated with mouse anti-Myc (at 1:200 dilution) and rabbit anti-HA antibodies (for TgVMP1, at 1:200 dilution), followed by appropriate secondary antibodies (Alexa Fluor 488-conjugated F(ab’)2 donkey anti-rabbit IgG and Alexa Fluor 647-conjugated F(ab’)2 donkey anti-mouse IgG) and DAPI (10 µg/ml). The cells were observed under 100× objective of the Zeiss LSM 880 confocal laser scanning microscope, images were acquired in Airyscan mode using Zen Black software, and processed using ImageJ software. Pearson’s correlation coefficient for co-localization was determined using JACoP plugin in Fiji.

To study whether PfVMP1_Myc_ or HsVMP1_Myc_ complement the growth of TgVMP1-depleted parasites, the parental RHTIR1-VMP1AID and complemented (RHTIR1-TgVMP1AID::PfVMP1_Myc_, RHTIR1-VMP1AID::HsVMP1_Myc_) parasites were grown without (–IAA) or with (+IAA) IAA, and compared for overall growth by plaque assay and intracellular development by replication assay as has been described in the “Effect of TgVMP1 depletion on parasite growth” section. Plaques were observed using the 1× objective at 40× digital zoom of AXIO Zoom.V16 microscope, and images were taken using the Zen Blue software.

To evaluate whether PfVMP1_Myc_ or HsVMP1_Myc_ can rescue the dense granule biogenesis defect, the parental RHTIR1-VMP1AID and complemented (RHTIR1-TgVMP1AID::PfVMP1_Myc_, RHTIR1-VMP1AID::HsVMP1_Myc_) parasites were grown without (–IAA) or with (+IAA) IAA, and compared for localization of dense granule protein GRA2 as has been described in the “Effect of TgVMP1 depletion on IVN formation, and dense granule secretion and biogenesis” section.

### Intracellular calcium monitoring

RHTIR1-VMP1AID parasites were cultured without (-IAA) or with (+IAA) indole-3-acetic acid for 14 hrs, and tachyzoites were purified as has been described in the “Parasite culture” section. The parasites were evaluated for intracellular calcium levels using Fluo-4 AM as has been described previously ^93^. The purified parasites were washed with Ringer’s solution (10 mM HEPES, 2 mM CaCl_2_, 1 mM MgCl_2_, 4.5 mM KCl, 155 mM NaCl, 10 mM D-glucose, pH 7.4), incubated with 250 nM of Fluo-4 AM for 10 min, washed with and resuspended in Ringer’s solution, transferred to poly-L-lysine coated glass slides, and observed under the 100× objective of Olympus FV3000 confocal microscope. Images were acquired using Fluoview FV31S-SW software, the fluorescence signal intensity/tachyzoite was measured using the “multi measure” option of the ROI manager of ImageJ software, and calculated using the formula “IntDen-(Area of interest × mean signal intensity of the background)”. Multiple parasites (-IAA, n = 415; +IAA, n = 356) from three independent experiments were observed. The mean signal intensity/parasite of +IAA parasites was presented as a percentage of the mean signal intensity/parasite of -IAA parasites.

### Transmission electron microscopy

RHTIR1-VMP1AID parasites were cultured with 0.1% ethanol (-IAA) or 500 µM indole-3-acetic acid (+IAA) as has been described in the “Effect of TgVMP1 depletion on gliding motility” section. Intracellular parasites were evaluated for IVN in PVs, and purified tachyzoites were evaluated for rhoptries and LDs by transmission electron microscopy (TEM) as has been previously described ^94^. The cells were washed twice with 0.1 M sodium cacodylate buffer (pH 7.2), and fixed with 2.5% glutaraldehyde for 2 hrs at 4⁰C. The fixed cells were scraped off, washed thrice with 0.1 M sodium cacodylate buffer for 5 min each, fixed with 1% osmium tetroxide (in collidine buffer) for 30 min, washed thrice with 0.1 M sodium cacodylate buffer, and stained *en bloc* with 2% uranyl acetate for 30 min. The cells were washed with 0.1 M sodium cacodylate buffer, dehydrated sequentially using increasing concentration of ethanol (50%, 70%, 95% and 100%), each for 3 min with 3 repeats. The cells were washed twice with propylene oxide, each for 5 min, treated with 1:1 mixture of epoxy and propylene oxide for 15 min, followed by 3:1 mixture of epoxy and propylene oxide for 20 min. The cells were incubated with pure epoxy for 10 min, and then polymerized in epoxy at 62⁰C for 18 hrs. Ultrathin sections (60-70 nm thickness) were cut using Leica ultramicrotome with a glass knife. The sections were transferred onto a copper grid, air dried, stained with uranyl acetate for 1 hr, followed by lead citrate for 5 min. The sections were imaged using Talos L120C transmission electron microscope equipped with CETA camera, and images were analyzed using Acquisition-velox software. Purified RHTIR1-VMP1AID parasites were also processed for TEM analysis by essentially following the same protocol.

## Statistical analysis

The significance of difference between two groups was determined by using two-tailed Student’s t-test. Statistical significance is indicated as P-value where * is p<0.05, ** is p<0.01, *** is p<0.001.

## Data availability

TgVMP1 IP-MS data has been deposited to PRIDE database with the identifier PXD056007.

## Acknowledgments

The authors are thankful to the staff of Advanced Microscopy and Imaging, Proteomics, Cryo-EM, and Animal House facilities of the Centre for Cellular and Molecular Biology for their assistance in imaging, mass spectrometry, EM imaging and animal related experiments, respectively. This study was supported by funds from Council of Scientific and Industrial Research, India (MLP0148) and Department of Biotechnology, India (BT/PR53849/BMS2/156/10/2024 (CN 20169)) to PS. GSR, KS, and MKB are the recipients of PhD fellowship from the Council of Scientific and Industrial Research, India. SMG is a recipient of PhD fellowship from University Grants Commission, India. The funders had no role in study design, data collection and analysis, decision to publish, or preparation of the manuscript.

## Author contributions

GSR and PSS conceived the study, designed the experiments, interpreted the data, and wrote the manuscript. GSR performed the majority of experiments. SMG performed egress assays, in vivo mouse experiments, and mass spectrometry. KS maintained and provided HFF cells for infection, and carried out invasion and microneme secretion assays. MKB constructed ER-GFP and HsVMP1 plasmids. ASD assisted in establishing the *T. gondii* culture, and provided technical guidance for construction of plasmids and transfection of *Toxoplasma*.

## Supplementary Figures S1–S12

Figure S1. PfVMP1 and TgVMP1 share the domain architecture of DedA superfamily.

Figure S2. Generation of PfVMP1GFP_KI_ and RHTIR1-VMP1AID parasites.

Figure S3. Dual-knock-down strategy for PfVMP1.

Figure S4. Auxin-inducible knock-down strategy for PfVMP1.

Figure S5. TgVMP1 depletion did not affect SAG1 and IMC1 transport.

Figure S6. Conoid extrusion is not affected upon TgVMP1 depletion.

Figure S7. TgVMP1 depletion did not affect microneme organization.

Figure S8. TgVMP1 is crucial for IVN formation and dense granule biogenesis.

Figure S9. TgVMP1 is crucial for ER organization.

Figure S10. TgVMP1 immunoprecipitation.

Figure S11. TgVMP1 and PfVMP1 are lipid scramblases, and restoration of the ER-localized scramblase activity rescued TgVMP1-depleted parasites.

Figure S12. TgVMP1 depletion elevated calcium level.

Figure S13. A model for the functions of TgVMP1.

## Supplementary Tables S1-S3

Table S1. VMP1 homologues in reference apicomplexan parasites and model organisms.

Table S2. Proteins identified in the TgVMP1 immunoprecipitates.

Table S3. Primers and synthetic DNAs used in this study.

