## Supplementary Tables for "An ER-resident lipid scramblase is crucial for biogenesis and function of apicomplexan parasite secretory organelles"

### List of tables

Table S1. VMP1 homologues in reference apicomplexan parasites and model organisms.

Table S2. Proteins identified in the TgVMP1 immunoprecipitates.

Table S3. Primers and synthetic DNAs used in this study.

**Table S1**. **VMP1 homologues in reference apicomplexan parasites and model organisms.** The apicomplexan VMP1 homologs are shaded. Percent identity of each protein with PfVMP1, TgVMP1 and HsVMP1 is shown. *Indicates a VMP1 homolog could not be identified.

| **Family** | **Organism** | **VMP1 homolog** | **% Identity** | | |
|---|---|---|---|---|---|
| | | | **PfVMP1** | **TgVMP1** | **HsVMP1** |
| Haemoproteidae | *Haemoproteus tartakovskyi* strain SISKIN1 | Htart_000483300 (Conserved *Plasmodium* membrane protein, unknown function), partial sequence | 61.5 | 39.5 | 27.2 |
| Plasmodiidae | *Hepatocystis sp. ex Piliocolobus tephrosceles, strain* 2019 | HEP_00034000 (Vacuole membrane protein 1, putative) | 70.3 | 40.1 | 32.6 |
| | *Plasmodium falciparum* 3D7 | PF3D7_1474600 (Vacuole membrane protein 1, putative) | 100 | 41.8 | 28.7 |
| Babesiidae | *Babesia bovis* T2Bo | * | - | - | - |
| Theileriidae | *Cytauxzoon felis* Winnie | * | - | - | - |
| | *Theileria parva strain* Muguga | * | - | - | - |
| Cryptosporidiidae | *Cryptosporidium parvum* Iowa II | cgd5_3820 (VMP1 like integral membrane protein) | 38.4 | 37.0 | 28.1 |
| Eimeriidae | *Cyclospora cayetanensis* CHN_HEN01 | cyc_05733 (Hypothetical protein) | 40.7 | 53.8 | 29.8 |
| | *Eimeria tenella* Houghton 2021 | ETH2_1533500 (Hypothetical protein) | 40.1 | 43.6 | 28.8 |
| Sarcocystidae | *Besnoitia besnoiti* Bb-Ger1 | BESB_014990 (TDC1, putative) | 42.2 | 78.3 | 29.1 |
| | *Hammondia hammondi* strain H.H.34 | HHA_244370 (TDC1, putative) | 41.7 | 97.1 | 28.8 |
| | *Sarcocystis neurona* SN3 | SN3_01400325 (TDC1, putative) | 41.1 | 51.3 | 31.1 |
| | *Cystoisospora suis* Wien I | CSUI_010169 (TDC1, putative) | 41.6 | 71.5 | 28.2 |

| | | | | | |
|---|---|---|---|---|---|
| | *Neospora caninum* Liverpool | NCLIV_018930 (GI15163, related) | 40.7 | 85.9 | 28.7 |
| | *Toxoplasma gondii* ME49 | TGME49_244370 (TDC1, putative) | 42.60 | 100 | 29.1 |
| Gregarinidae | *Gregarina niphandrodes* | * | - | - | - |
| Porosporidae | *Porospora* cf. gigantea A | * | - | - | - |
| Hominidae | *Homo sapiens* | Q96GC9 9 (Vacuole membrane protein 1) | 29.3 | 30.1 | 100 |
| Drosophilidae | *Drosophila melanogaster* | Q9W2S1 (TANGO5) | 28.2 | 29.6 | 52.0 |
| Rhabditidae | *Caenorhabditis elegans* | Q9XWU8 (Ectopic P granules protein 3 (epg-3)) | 30.4 | 30.2 | 48.7 |
| Brassicaceae | *Arabidopsis thaliana* | Q5XF36 (Vacuole membrane protein KMS1) | 28.8 | 30.3 | 34.8 |
| | | F4I8Q7 (Vacuole membrane protein KMS2) | 27.8 | 35.6 | 33.5 |
| Chlamydomonadaceae | *Chlamydomonas reinhardtii* | A0A2K3DNE8 (Vacuole membrane protein) | 30.0 | 39.6 | 36.0 |
| Dictyosteliaceae | *Dictyostelium discoideum* | Q54NL4 (Vacuole membrane protein 1 homolog) | 30.0 | 29.1 | 43.9 |
| Cyprinidae | *Danio rerio* | Q6NYY9 (Vacuole membrane protein) | 26.6 | 28.5 | 73.2 |

**Table S2. Proteins identified in the TgVMP1 immunoprecipitates.** The table shows UniProt identification number (ID), score (Sc), coverage (Co) and unique peptides (UP) for the proteins identified in independent experiments. The cellular location, function, and related reference for the characterized/reported proteins are also mentioned. No MS data (-) in some of the experiments and unavailability of information for some of proteins (Unknown) are mentioned. The mass spectrometry proteomics data have been deposited to the ProteomeXchange Consortium via the PRIDE partner repository with the dataset identifier "PXD056007".

| **ID** | **Protein** | **Experiment 1** | | | **Experiment 2** | | | **Experiment 3** | | | **Cellular location** | **Function** | **Reference** |
|---|---|---|---|---|---|---|---|---|---|---|---|---|---|
| | | **Sc** | **Co** | **UP** | **Sc** | **Co** | **UP** | **Sc** | **Co** | **UP** | | | |
| A0A7J6K7V4 | TDC1, putative (TgVMP1) | 56.19 | 39.38 | 11 | 59.37 | 32.36 | 11 | 52.61 | 30.65 | 11 | ER | Regulator of secretory organelles | This study |
| A0A7J6KD16 | Dense granule protein GRA12 | 26.67 | 19.27 | 9 | 14.86 | 29.13 | 5 | 29.89 | 30.28 | 8 | PV, IVN | Virulence | [1-3] |
| A0A7J6K3E8 | 60S ribosomal protein L18a | 14.83 | 42.08 | 3 | 8.83 | 42.62 | 4 | 4.60 | 16.94 | 1 | Ribosome | Protein translation | Unknown |
| A0A7J6JUT1 | Ribosomal protein RPS9 | 14.80 | 48.40 | 5 | 4.37 | 19.15 | 2 | 3.83 | 28.19 | 2 | Ribosome | Protein translation | Unknown |
| A0A7J6JUS3 | Transmembrane protein | 12.23 | 32.19 | 4 | 1.72 | 33.62 | 2 | 6.00 | 19.66 | 1 | Unknown | | |
| A0A7J6K107 | Apical cap protein AC5 | 12.07 | 27.27 | 2 | 10.25 | 32.76 | 1 | 13.85 | 27.10 | 2 | Cortical microtubules | Unknown | [4] |
| A0A7J6K6L5 | Preprotein translocase Sec61, putative | 11.89 | 18.18 | 4 | 15.35 | 18.18 | 4 | 21.10 | 23.68 | 6 | Protein translocase complex on ER | Protein translocation into the ER lumen/ER membrane | Unknown |
| Q3LRU1 | Photosensitized INA-labelled protein PHIL1 | 11.14 | 35.76 | 4 | 9.83 | 33.94 | 4 | 13.75 | 21.82 | 3 | Pellicle cytoskeleton | Maintenance of parasite morphology, Host cell invasion | [5,6] |
| A0A7J6KF30 | Ribosomal protein RPS4 | 10.26 | 28.21 | 5 | 5.53 | 15.71 | 1 | 5.91 | 26.60 | 4 | Ribosome | Protein translation | Unknown |
| A0A7J6JXJ2 | Ribosomal protein RPL7 | 9.74 | 30.62 | 2 | 7.54 | 24.03 | 2 | 4.77 | 17.83 | 1 | Ribosome | Protein translation | Unknown |
| P90614 | SAG-related sequence SRS29A | 8.73 | 19.76 | 1 | 6.85 | 15.00 | 2 | 9.89 | 31.67 | 4 | Parasite surface | Antigen | [7,8] |
| A0A7J6K0A6 | Ribosomal protein RPS19 | 7.74 | 20.58 | 3 | 2.81 | 28.81 | 2 | 6.64 | 11.11 | 2 | Ribosome | Protein translation | Unknown |

| | | | | | | | | | | | | | |
|---|---|---|---|---|---|---|---|---|---|---|---|---|---|
| A0A7J6K582 | Pyruvate dehydrogenase complex subunit PDH-E2 | 7.73 | 17.33 | 2 | 1.65 | 20.13 | 1 | 0.00 | 13.35 | 1 | Mitochondria | Converts pyruvate into acetyl-CoA | Unknown |
| A0A7J6K9F1 | Ribosomal protein RPL32 | 7.42 | 9.94 | 1 | 2.05 | 18.13 | 1 | 5.14 | 9.94 | 1 | Ribosome | Protein translation | Unknown |
| A0A7J6K6B4 | Glideosome-associated protein with multiple-membrane spans GAPM2B | 7.15 | 6.40 | 2 | 5.69 | 17.60 | 1 | 9.78 | 15.20 | 3 | IMC | Shape and rigidity of apicomplexan zoites, IMC biogenesis | [9,10] |
| A0A7J6K2S0 | Ribosomal protein RPS11 | 5.36 | 26.71 | 2 | 11.35 | 43.48 | 5 | 11.39 | 68.32 | 5 | Ribosome | Protein translation | Unknown |
| Q7Z289 | Gliding-associated protein 45 (GAP45) | 5.26 | 28.16 | 2 | 4.56 | 25.31 | 2 | 7.59 | 12.65 | 1 | IMC | Motility/ assembly of the myosin XIV motor complex | [11,12] |
| A0A7J6JZ28 | Rhoptry neck protein RON3 | 3.99 | 20.28 | 2 | 0.00 | 17.72 | 2 | 0.00 | 22.37 | 1 | Rhoptry | *P. falciparum* RON3 is required for protein export | [13,14] |
| A0A7J6KAB3 | hypothetical protein | 3.59 | 9.17 | 1 | 1.82 | 14.92 | 1 | 1.71 | 12.82 | 1 | Unknown | | |
| Q9U4T9 | Dense granule protein GRA8 | 3.06 | 21.72 | 2 | 1.92 | 16.10 | 1 | 1.99 | 20.22 | 1 | Dense granules, PVM | Modulator of host processes | [3,15,16] |
| A0A7J6K4C9 | Ribosomal protein RPL30 | 2.76 | 11.11 | 1 | 3.81 | 11.11 | 1 | 9.72 | 49.07 | 2 | Ribosome | Protein translation | Based on model systems |
| A0A7J6K6Y0 | Transmembrane protein | 2.66 | 25.61 | 2 | 4.40 | 9.51 | 2 | 2.11 | 31.22 | 1 | Unknown | | |
| A0A7J6KAG7 | Ribosomal protein RPL19 | 2.36 | 12.30 | 1 | 1.89 | 4.81 | 1 | 2.06 | 17.11 | 1 | Ribosome | Protein translation | Unknown |
| A0A7J6K7V6 | Prohibitin, putative | 2.33 | 32.47 | 1 | 2.34 | 28.41 | 1 | 0.00 | 25.09 | 1 | Unknown | | |
| A0A7J6K3G9 | Hypothetical protein | 2.32 | 10.67 | 1 | 2.39 | 10.67 | 1 | 0.00 | 10.67 | 1 | Unknown | | |
| A0A7J6KAQ4 | Ribosomal protein RPL23A | 2.07 | 10.78 | 1 | 2.74 | 24.55 | 1 | 2.13 | 31.14 | 2 | Ribosome | Protein synthesis | Unknown |
| A0A7J6JY44 | rRNA pseudouridine synthase | 1.94 | 6.14 | 1 | 0.00 | 7.72 | 1 | 0.00 | 21.39 | 1 | Unknown | | |
| Q1JSG0 | Formate/nitrite transporter protein | 1.61 | 1.70 | 1 | 4.12 | 8.25 | 1 | 4.59 | 13.59 | 1 | Plasma membrane | Lactate transport | [17] |

| | | | | | | | | | | | | | |
|---|---|---|---|---|---|---|---|---|---|---|---|---|---|
| A0A7J6KDU7 | SAG-related sequence SRS17B | 0.00 | 6.33 | 1 | 2.21 | 16.89 | 2 | 2.12 | 26.39 | 1 | Parasite membrane | Antigen | [7,8] |
| A0A7J6JWW4 | Myosin F | 0.00 | 14.08 | 1 | 1.89 | 13.98 | 2 | 3.91 | 15.31 | 6 | Myosin motor | Actin organization | [18,19] |
| A0A7J6JXG6 | Hypothetical protein | 0.00 | 13.97 | 1 | 2.01 | 11.17 | 1 | 2.06 | 25.70 | 1 | Unknown | | |
| A0A7J6K9Y6 | Myosin-light-chain kinase | 73.46 | 37.70 | 12 | 32.87 | 43.11 | 8 | - | - | - | Plasma membrane | Motility and invasion | [20] |
| A0A7J6K1K2 | Dynein, axonemal, heavy chain 2 family protein | 27.07 | 14.15 | 1 | 24.33 | 21.4 | 1 | - | - | - | Unknown | | |
| B9Q286 | Inner membrane complex protein 18 | 23.69 | 47.76 | 6 | 14.45 | 51.87 | 5 | - | - | - | IMC | Dispensable | [4] |
| Q9U540 | Chaperonin protein BiP | 20.52 | 24.70 | 6 | 12.68 | 13.92 | 3 | - | - | - | ER lumen | Protein folding | Unknown |
| A0A7J6KEI3 | Mitochondrial association factor 1a | 17.61 | 24.60 | 4 | 0 | 12.64 | 1 | - | - | - | Secreted protein | Recruitment of host mitochondria and modulation of the host response | [21] [22,23] |
| A0A7J6K461 | PWI domain-containing protein | 13.67 | 8.95 | 1 | 15.12 | 13.81 | 1 | - | - | - | Unknown | | |
| A0A7J6JYE5 | DEAD (Asp-Glu-Ala-Asp) box polypeptide 17 | 10.97 | 25.09 | 4 | 0 | 18.36 | 2 | - | - | - | Unknown | | |
| A0A7J6KC78 | Pyruvate dehydrogenase complex subunit PDH-E3I | 10.58 | 10.78 | 4 | 3.32 | 16.64 | 2 | - | - | - | Mitochondria | Converts pyruvate into acetyl-CoA | Unknown |
| A0A7J6JZK8 | Hypothetical protein | 10.15 | 9.37 | 1 | 3.63 | 12.31 | 1 | - | - | - | Unknown | | |
| A0A7J6KCY2 | Sortilin-like receptor SORTLR (SORTLR) | 6.24 | 11.42 | 3 | 2.25 | 12.68 | 1 | - | - | - | Golgi, endosomal compartments | Biogenesis of micronemes and rhoptries | [24] |
| A0A7J6JVM1 | Ribosomal protein RPL21 | 4.93 | 36.31 | 1 | 2.28 | 12.74 | 1 | - | - | - | Ribosome | Protein synthesis | Unknown |
| A0A7J6JTW4 | Inner membrane complex protein IMC19 | 4.04 | 17.63 | 3 | 1.88 | 16.84 | 1 | - | - | - | IMC | Unknown | [4] |
| A0A7J6JVI4 | Hypothetical protein | 1.63 | 12.72 | 1 | 3.35 | 9.56 | 1 | - | - | - | Unknown | | |

| | | | | | | | | | | | | | |
|---|---|---|---|---|---|---|---|---|---|---|---|---|---|
| A0A7J6JXD5 | Ribosomal protein RPL11 | 17.90 | 27.43 | 4 | - | - | - | 1.78 | 14.29 | 1 | Ribosome | Protein synthesis | Unknown |
| A0A7J6K9C0 | SAG-related sequence SRS25 | 10.18 | 33.51 | 3 | - | - | - | 15.23 | 54.45 | 3 | Parasite surface | Antigen | [7,8] |
| A0A7J6JYL4 | Ribosomal protein RPL6 | 10.15 | 39.90 | 4 | - | - | - | 0 | 17.1 | 1 | Ribosome | Protein synthesis | Unknown |
| A0A7J6K9M5 | 40S ribosomal protein S26 | 5.80 | 47.32 | 1 | - | - | - | 5.35 | 32.14 | 1 | Ribosome | Protein synthesis | Unknown |
| A0A7J6KEY2 | Carbonic anhydrase-related protein CARP | 5.78 | 13.87 | 3 | - | - | - | 0 | 1.54 | 1 | Rhoptry | Maintenance of rhoptry morphology | [25] |
| A0A7J6KF01 | IMC sutures component ISC3 | 5.73 | 7.86 | 2 | - | - | - | 1.96 | 15.87 | 1 | IMC | Maintenance of the parasite shape | [26] |
| A0A7J6K342 | SPFH domain/Band 7 family protein | 5.49 | 20.00 | 2 | - | - | - | 1.95 | 36.55 | 1 | Unknown | | |
| A0A7J6K3Q7 | Mediator complex subunit MED14 | 5.06 | 12.11 | 1 | - | - | - | 0 | 15.05 | 1 | Unknown | | |
| A0A7J6KAV8 | Microneme protein MIC1 | 4.64 | 4.39 | 2 | - | - | - | 3.59 | 17.98 | 1 | Micronemes | Binds lectin and modulates immune responses | [27-29] |
| A0A7J6K221 | Ribosomal protein RPL27 | 4.26 | 48.63 | 2 | - | - | - | 0 | 21.92 | 1 | Ribosome | Protein synthesis | Unknown |
| A0A7J6JUE4 | Hypothetical protein | 3.62 | 17.90 | 2 | - | - | - | 0 | 14.32 | 1 | Unknown | | |
| A0A7J6KDE7 | Dense granule protein GRA25 | 3.53 | 19.55 | 1 | - | - | - | 0 | 30.45 | 1 | PVM, secreted into host cytoplasm | Immune modulator | [30] |
| A0A7J6KCI3 | Ribosomal protein RPL31 | 1.90 | 29.17 | 2 | - | - | - | 2.45 | 45 | 2 | Ribosome | Protein synthesis | Unknown |
| A0A7J6KC45 | Tubulin-tyrosine ligase family protein | 1.71 | 15.49 | 1 | - | - | - | 0 | 10.48 | 1 | Unknown | | |
| Q86DU2 | Importin alpha, putative | 0.00 | 19.08 | 1 | - | - | - | 0 | 19.08 | 1 | Nucleus | Nucleocytoplasmic transporter | [31] |
| A0A7J6JWC9 | Dense granule protein GRA71 | 0.00 | 13.65 | 1 | - | - | - | 1.72 | 21.98 | 1 | Dense granules | Required for parasite fitness in IFNγ-stimulated cells | [32] |
| A0A7J6K410 | Hypothetical protein | 0.00 | 2.21 | 1 | - | - | - | 0 | 5.71 | 1 | Unknown | | |

| | | | | | | | | | | | | | |
|---|---|---|---|---|---|---|---|---|---|---|---|---|---|
| A0A7J6KC74 | Apicomplexan kinetochore protein AKIT5, putative | 0.00 | 12.92 | 1 | - | - | - | 0 | 12.82 | 1 | Kinetochore | Chromosome segregation | [33] |
| A0A0B5KXC2 | Rhoptry protein ROP8 | - | - | - | 28.8 | 33.91 | 3 | 26.35 | 33.22 | 2 | Rhoptry | Unknown | Unknown |
| Q6J6C2 | Aspartyl protease ASP3 | - | - | - | 27.05 | 25.82 | 7 | 22.58 | 26.59 | 7 | Endosomal-like compartment | Processing of a some microneme and rhoptry proteins | [34] |
| A0A7J6KDJ4 | MSP (Major sperm protein) domain-containing protein | - | - | - | 24.19 | 61.92 | 4 | 15.26 | 53.97 | 2 | Unknown | | |
| A0A7J6K5R5 | Ribosomal protein RPS3 | - | - | - | 15.42 | 22.98 | 4 | 14.04 | 28.51 | 4 | Ribosome | Protein synthesis | Unknown |
| A0A7J6KA11 | Ribosomal protein RPL9 | - | - | - | 11.47 | 14.21 | 2 | 13.79 | 29.47 | 3 | Ribosome | Protein synthesis | Unknown- |
| A0A7J6K279 | Exosome complex exonuclease RRP44, putative | - | - | - | 13.86 | 10.36 | 1 | 12.1 | 20.71 | 1 | Unknown | | |
| A0A7J6K5K9 | ATP synthase subunit alpha | - | - | - | 12.53 | 29.91 | 5 | 11.34 | 23.89 | 3 | Unknown | | |
| A0A7J6K4Z0 | Putative kinesin heavy chain | - | - | - | 12.56 | 17.12 | 5 | 10.98 | 20.9 | 6 | Unknown | | |
| A0A7J6KEZ9 | Protein tyrosine phosphatase, putative | - | - | - | 4.67 | 17.07 | 2 | 10.88 | 15.51 | 5 | Unknown | | |
| A0A7J6K2Z6 | Cyst wall protein CST7 | - | - | - | 9.23 | 16.38 | 4 | 10.35 | 10.7 | 4 | Cyst wall | Unknown | [35] |
| A0A7J6KDU3 | Hypothetical protein | - | - | - | 9.95 | 21.17 | 1 | 8.18 | 30.18 | 1 | Unknown | | |
| A0A7J6K186 | IMC sutures component ISC1 | - | - | - | 4.67 | 22.08 | 2 | 7.81 | 17.08 | 4 | Alveolar sutures of the IMC | Unknown | [4] |
| A0A7J6K3K4 | Ribosomal protein RPS23 | - | - | - | 8.04 | 29.37 | 2 | 7.43 | 29.37 | 2 | Ribosome | Protein synthesis | Unknown |
| A0A7J6KC31 | Hypothetical protein | - | - | - | 13.1 | 35.5 | 5 | 7.36 | 17.48 | 3 | Unknown | | |
| A0A7J6KDR8 | Rhoptry kinase family protein ROP26 (Incomplete catalytic triad) | - | - | - | 4.52 | 18.39 | 3 | 7.27 | 25.98 | 3 | Rhoptry | Unknown | [36] |

| | | | | | | | | | | | | | |
|---|---|---|---|---|---|---|---|---|---|---|---|---|---|
| A0A7J6K2X7 | Ribosomal protein RPS16 | - | - | - | 12.25 | 18.96 | 3 | 7.13 | 36.97 | 4 | Ribosome | Protein synthesis | Unknown |
| A0A7J6JTY2 | GTP-binding nuclear protein ran/tc4 | - | - | - | 11.74 | 14.85 | 3 | 6.7 | 19.65 | 3 | Unknown | | |
| Q8MUM2 | Facilitative glucose transporter GT1 | - | - | - | 4.2 | 9.15 | 1 | 6.51 | 10.04 | 2 | Unknown | | |
| A0A7J6KCN4 | Ribosomal protein RPS2 | - | - | - | 8.78 | 26.02 | 5 | 6.32 | 21.56 | 4 | Ribosome | Protein synthesis | - |
| A0A7J6K3U7 | 40S ribosomal protein S3a | - | - | - | 17.6 | 53.67 | 9 | 6.2 | 33.59 | 4 | Ribosome | Protein synthesis | - |
| B9PK73 | Rhoptry protein 13 | - | - | - | 5.24 | 29.75 | 3 | 5.93 | 18 | 3 | Rhoptry | Not essential | [37] |
| A0A7J6K5C5 | Phosphatidylserine decarboxylase | - | - | - | 7.96 | 16.52 | 3 | 5.8 | 13.11 | 3 | Unknown | | |
| A0A7J6K7Z5 | MIZ/SP-RING zinc finger domain-containing protein | - | - | - | 3.05 | 13.96 | 1 | 5.32 | 11.27 | 1 | Unknown | | |
| A0A7J6JVV5 | Phosphate transporter MPT | - | - | - | 8.45 | 17.95 | 5 | 5.13 | 26.51 | 3 | Mitochondria | Dispensable | [38] |
| A0A140H545 | Hypothetical protein | - | - | - | 2.76 | 29.66 | 1 | 5.06 | 8.74 | 1 | Unknown | | |
| A0A7J6JWZ2 | Ribosomal protein RPS15 | - | - | - | 3.68 | 7.26 | 1 | 4.86 | 6.97 | 1 | Ribosome | Protein synthesis | - |
| A0A7J6K8C3 | Rhoptry protein ROP12 | - | - | - | 0 | 2.97 | 1 | 4.78 | 13.98 | 2 | Rhoptry | Unknown | [13] |
| A0A7J6K3V8 | V-type proton ATPase subunit a1 | - | - | - | 0 | 20.35 | 1 | 4.41 | 21.23 | 2 | Plasma membrane and to acidic vesicles | Maturation of the rhoptry and microneme proteins, and their processing proteases | [39] |
| A0A7J6JYR0 | Ribosomal protein RPL27A | - | - | - | 4.15 | 16.75 | 1 | 4.3 | 35.6 | 2 | Ribosome | Protein synthesis | - |
| A0A7J6K4I8 | Ribosomal protein RPL38 | - | - | - | 2.24 | 42.86 | 1 | 4.28 | 35.71 | 1 | Ribosome | Protein synthesis | - |
| A0A7J6JWV3 | Ras-related protein RAB2 | - | - | - | 0 | 12.15 | 1 | 4.16 | 40.65 | 2 | Early secretory pathway | ER/Golgi transport | [40] |
| A0A7J6JZC9 | SAG-related sequence SRS52A | - | - | - | 7.6 | 34.36 | 1 | 3.84 | 44.13 | 1 | Unknown | | |

| | | | | | | | | | | | | | |
|---|---|---|---|---|---|---|---|---|---|---|---|---|---|
| A0A7J6K147 | Ribosomal protein RPS15A | - | - | - | 3.9 | 14.72 | 2 | 3.81 | 27.71 | 2 | Ribosome | Protein synthesis | - |
| A0A7J6K876 | Glideosome-associated protein with multiple-membrane spans GAPM1A | - | - | - | 3.95 | 22.62 | 1 | 3.52 | 17.05 | 2 | Alveoli | Maintains microtubule stability | [9] |
| A0A7J6JVM3 | Ribosomal protein RPL35 | - | - | - | 3.32 | 40.45 | 1 | 3.5 | 8.99 | 1 | Ribosome | Protein synthesis | - |
| A0A7J6K9H7 | Ribosomal protein RPL24 | - | - | - | 9.06 | 38.06 | 1 | 3.39 | 18.06 | 1 | Ribosome | Protein synthesis | - |
| Q1JTB8 | Rhoptry kinase family protein ROP37 (Incomplete catalytic triad) | - | - | - | 3.54 | 12.5 | 2 | 3.29 | 20.23 | 1 | Rhoptry | Dispensable | [41] |
| A0A7J6KAX9 | Ribosomal protein RPL13A | - | - | - | 1.65 | 23.43 | 1 | 3.12 | 17.83 | 1 | Ribosome | Protein synthesis | Unknown |
| A0A7J6JU81 | 40S ribosomal protein S8 | - | - | - | 5 | 15.61 | 2 | 2.7 | 42.44 | 2 | Ribosome | Protein synthesis | Unknown |
| A0A7J6K232 | Thioredoxin-like protein 2 | - | - | - | 9.33 | 33.33 | 4 | 2.67 | 11.67 | 1 | Cortical microtubules | Unknown | [42] |
| A0A7J6K3X5 | Apical annuli protein AAP4 | - | - | - | 6.98 | 12.12 | 2 | 2.59 | 8.32 | 1 | Apical annuli | Unknown | [43] |
| A0A7J6K482 | Nucleoporin autopeptidase | - | - | - | 1.67 | 8.36 | 1 | 2.57 | 5.7 | 1 | Nuclear pore complex | nuclear transport | [44] |
| A0A7J6KC37 | IMC sutures component ISC6 | - | - | - | 4.97 | 14.96 | 2 | 2.57 | 23.03 | 2 | IMC sutures | Unknown | [26] |
| A0A7J6KDB0 | Microneme protein MIC3 | - | - | - | 1.81 | 8.62 | 1 | 2.56 | 39.16 | 1 | Micronemes | Invasion of the host cell | [45] |
| A0A7J6KEY8 | Helicase associated domain (Ha2) protein | - | - | - | 14.01 | 12.94 | 6 | 2.5 | 10.7 | 2 | Unknown | | |
| A0A7J6K459 | Transverse sutures component TSC5 | - | - | - | 2.83 | 4.25 | 1 | 2.36 | 13.73 | 1 | IMC sutures | Unknown | [26] |
| A0A7J6K0G3 | Hypothetical protein | - | - | - | 1.72 | 14.67 | 1 | 2.15 | 13.16 | 1 | Unknown | | |
| Q6V7J6 | Ribosomal protein RPL37A | - | - | - | 1.63 | 31.25 | 2 | 2.02 | 8.33 | 1 | Ribosome | Protein synthesis | - |

| | | | | | | | | | | | | | |
|---|---|---|---|---|---|---|---|---|---|---|---|---|---|
| A0A7J6K0Z3 | Microneme protein MIC7 | - | - | - | 3.61 | 23.24 | 3 | 1.98 | 24.41 | 1 | Micronemes | Invasion of the host cell | [46] |
| A0A7J6K1R4 | Armadillo repeats only protein ARO | - | - | - | 0 | 39.78 | 1 | 1.95 | 28.1 | 1 | Rhoptry membrane | Apical positioning of rhoptries | [47] |
| A0A7J6JZ56 | Ribosomal protein | - | - | - | 4.79 | 32.72 | 1 | 1.92 | 11.98 | 1 | Ribosome | Protein synthesis | Unknown |
| A0A7J6K3I1 | 60S ribosomal protein L13 | - | - | - | 7.26 | 25.07 | 3 | 1.9 | 15.77 | 1 | Ribosome | Protein synthesis | Unknown |
| A0A7J6KAF2 | Ribosomal protein RPL14 | - | - | - | 6.32 | 26.11 | 4 | 1.88 | 32.48 | 1 | Ribosome | Protein synthesis | Unknown |
| A0A7J6K811 | DEAD/DEAH box helicase domain-containing protein | - | - | - | 1.68 | 10.75 | 1 | 1.86 | 9.7 | 1 | Unknown | | |
| A0A7J6KA90 | Inhibitor of STAT1-dependent transcription TgIST | - | - | - | 5.6 | 13.15 | 3 | 1.84 | 13.82 | 2 | Parasite and host | Inhibits STAT1-dependent proinflammatory gene expression | [48] |
| A0A7J6K9M8 | Rhoptry protein ROP9 | - | - | - | 4.2 | 32.58 | 3 | 1.63 | 21.25 | 2 | Rhoptries | Required for invasion and development | [49] |
| A0A7J6JYL3 | Putative cyclase-associated protein | - | - | - | 4.42 | 11.79 | 1 | 1.6 | 19.49 | 1 | Unknown | | |
| A0A7J6JV63 | Hypothetical protein | - | - | - | 0 | 25.78 | 1 | 0 | 25.78 | 1 | Unknown | | |
| A0A7J6K5L9 | Pyruvate dehydrogenase E1 component subunit beta | - | - | - | 0 | 18.72 | 1 | 0 | 10.85 | 1 | Mitochondria | Converts pyruvate into acetyl-CoA | Unknown |
| A0A7J6JWC0 | Hypothetical protein | - | - | - | 2.73 | 25.98 | 2 | 0 | 17.38 | 2 | Unknown | | |
| A0A7J6K6H7 | E3 ubiquitin ligase, putative | - | - | - | 1.66 | 9.54 | 1 | 0 | 10.97 | 1 | Unknown | | |
| A0A7J6KBM9 | Hypothetical protein | - | - | - | 0 | 6.72 | 1 | 0 | 14.66 | 1 | Unknown | | |
| A0A7J6KD11 | Guanylate-binding protein, N-terminal domain-containing protein | - | - | - | 0 | 20.45 | 1 | 0 | 13.74 | 1 | Unknown | | |

**Table S3. Primers and synthetic DNAs used in this study.** The restriction enzyme sites are underlined.

| **Primer Name** | **Sequence (5'→3')** |
|---|---|
| PvAc-Con-R | TAGGTATGCATACGTGAATGTACTGG |
| Hrp2-SeqF | CTTTTACAATATGAACATAAAGTACAAC |
| GFPSEQR1 | GTGCCCATTAACATCACCA |
| PcDT5U-R1 | GGATGCCATGCACATGCTTAGTACACAT |
| PfVMP1-F1 | TATAGGGGCCCTGTTTAGGAACAAATGATGAAGACAT |
| PfVMP1-R1 | CATTAGGTACCTTTTTTTATTTTTCTTGGTGTCATTTCGTTC |
| PfVMP1-5con | GTGTTATACGATTGTAGAAGTG |
| PfVMP1-3con | AAAACACAGGACATGTTCACA |
| PfVMP1-3con2 | CTCTCAAAATTGTAATCATCCAGGTAC |
| PfA8fl2-F1 | ATGGGTACCCCGGGATCCCCATCGCTTAAAGACGAAGTA |
| PfA8fl2-R1 | CCGCTCGAGGCAGCCTGTTTTATGTATCCAC |
| PfVMP1-fl1F | TATAGGCGGCCGCTGTTTAGGAACAAATGATGAAGACAT |
| PfVMP1-Fl2F | TAAAACCTAGGTGTGAACATGTCCTGTGTTTT |
| PfVMP1-Fl2R | ATCATAGGCGCCGGATCAGTTTTGTATACATTTGTATATGG |
| XTEN-mAID-2×HA | GGTACCAGTGGAAGTGAAACACCTGGAACAAGTGAAAGTGCAACACCAGAAAGTAAAGAAAAAATCAGCATGTCCAAAAGATCCAGCAAAACCACCAGCAAAAGCACAAGTAGTAGGATGGCCACCAGTTAGAAGTTATAGAAAAAATGTAATGGTATCATGTCAAAAATCAAGTGGTGGACCAGAAGCAGCAGCATTTGTAAAAGTATCAATGGATGGTGCTCCATATTTAAGAAAAATTGATTTAAGAATGTATAAATCTGGTGGTGGTGGAAGTTATCCTTATGATGTACCAGATTATGCTGGATCATATCCTTATGATGTTCCAGATTATGCATAACTCGAG |

| | |
|---|---|
| TIR1-HD (It contains sequences of PcDT5U, Tir1, Flag, 2A, hDHFR (HD), PcDT3U, LoxP regions) | ACCGGTATACATATAACATAATAGGAAATCATTAATAATTTAGCGAAATTCTACAAATTTTAAA<br>AAAAATAAAATAATCCTTTTTTGTTATTCCACAATATGTATTGATATATACATGTACAAATATTTT<br>TTTTTTTATTTTGTAAAAAAAATTGTATAGTGTACGAAAATGGGCATGTAGGAAAAATGTAGGG<br>GCATTTCCCATATGGTTGATGGCCATATAAATGTTATATATTAGCTTGTGTAGGGAATAAAAAC<br>GGTTCATAAATAATGGTTGAAAGGAATTTTTATATATTAATATAAACACTAGGTTAGGTAATTC<br>GCTTGTAAGAGGTACTCTCGTTTATGCAAAACTATTTGATATAGCATTTTAACAAGTACACATAT<br>ATATATGTAATATATATACTATATATATCTATTGCATGTGTACTAAACGTGTGCATGGCATCCCC<br>TTTTTCTCGTGTTTAAAACAGTTTGTATGATAAAATATAAAGGATTTGAAAAAGAGAAAAAAAT<br>ATATGATCTCATCCTATATAGCGCCATAATTTTTATTTGGGTTGAATAAAATTTTCTATTAAATTT<br>AGGTGTAAGTAAAATAATGGAATATATATAAGTACAATAAAAAAGTGCATAAATTAAAAAATT<br>TTTATAATAAATATTTTTTTTTAAAAAAGTCAATAATAATATTAAATATATATAACACAGGATTAT<br>ATATGTTCACTACAATTTTTTATATTATAATATAAATTATTTTCAATTTTCATTTTATTTTACATAC<br>ACTTTCCTTTTTTGTCACCATATTTTAATATTCACATATTTAGTTTAAGTACTGGCTATTTCTTTCT<br>ACATTTGCTAGTAACAATTGTGTAGTGCTTATATATATACACACACCTAAAATTTACAAAACTGC<br>AGATGACATATTTTCCAGAAGAAGTAGTAGAACATATATTTTCATTTTTACCAGCACAAAGAGA<br>TAGAAATACTGTTAGTTTAGTATGTAAAGTTTGGTATGAAATAGAAAGATTAAGTAGAAGAGG<br>AGTATTTGTTGGAAATTGTTATGCAGTTAGAGCTGGAAGAGTAGCAGCAAGATTTCCAAATGT<br>TAGAGCATTAACAGTAAAAGGAAAACCACATGGTGCAGATTTTAATTTAGTACCACCAGATTG<br>GGGAGGATATGCAGGACCTTGGATAGAAGCAGCAGCTAGAGGATGTCATGGATTAGAAGAAT<br>TAAGAATGAAAAGAATGGTAGTTAGTGATGAATCATTAGAATTATTAGCAAGAAGTTTTCCTA<br>GATTTAGAGCTTTAGTTTTAATATCATGTGAAGGTTTTAGTACAGATGGATTAGCAGCAGTAGC<br>AAGTCATTGTAAATTATTAAGAGAATTAGATTTACAAGAAAATGAAGTTGAAGATAGAGGACC<br>AAGATGGTTAAGTTGTTTTCCAGATAGTTGTACAAGTTTAGTTAGTTTAAATTTTGCTTGTATAA<br>AAGGAGAAGTAAATGCTGGAAGTTTAGAAAGATTAGTTAGTAGAAGTCCAAATTTAAGAAGTT<br>TAAGATTAAATAGATCAGTTAGTGTAGATACTTTAGCAAAAATATTATTAAGAACACCAAATTT<br>AGAAGATTTAGGAACTGGAAATTTAACAGATGATTTTCAAACAGAATCATATTTTAAATTAACA<br>AGTGCATTAGAAAAATGTAAAATGTTAAGAAGTTTAAGTGGATTTTGGGATGCAAGTCCAGTA<br>TGTTTAAGTTTTATATATCCATTATGTGCACAATTAACAGGTTTAAATTTATCATATGCACCAAC<br>ATTAGATGCAAGTGATTTAACAAAAATGATAAGTAGATGTGTTAAATTACAAAGATTATGGGT<br>ATTAGATTGTATAAGTGATAAAGGATTACAAGTAGTAGCATCAAGTTGTAAAGATTTACAAGA<br>ATTAAGAGTATTTCCAAGTGATTTTTATGTAGCTGGATATAGTGCAGTAACAGAAGAAGGATTA<br>GTAGCTGTATCATTAGGATGTCCAAAATTAAATAGTTTATTATATTTTTGTCATCAAATGACAAA<br>TGCAGCATTAGTAACAGTAGCTAAAAATTGTCCTAATTTTACAAGATTTAGATTATGTATATTAG<br>AACCTGGAAAACCAGATGTAGTAACAAGTCAACCATTAGATGAAGGTTTTGGAGCAATAGTTA |

| | |
|---|---|
| | GAGAATGTAAAGGTTTACAAAGATTAAGTATAAGTGGTTTATTAACAGATAAAGTTTTTATGTA<br>TATTGGAAAATATGCAAAACAATTAGAAATGTTATCAATAGCATTTGCTGGAGATTCAGATAAA<br>GGAATGATGCATGTAATGAATGGATGTAAAAATTTAAGAAAATTAGAAATTAGAGATAGTCCA<br>TTTGGAGATGCAGCTTTATTAGGAAATTTTGCTAGATATGAAACAATGAGAAGTTTATGGATG<br>AGTAGTTGTAATGTAACTTTAAAAGGATGTCAAGTTTTAGCTTCAAAAATGCCAATGTTAAATG<br>TAGAAGTTATAAATGAAAGAGATGGAAGTAATGAAATGGAAGAAAATCATGGAGATTTACCA<br>AAAGTAGAAAAATTATATGTTTATAGAACAACAGCTGGTGCAAGAGATGATGCTCCAAATTTT<br>GTAAAAATATTAGGTGGTGGTGGAAGTGATTATAAAGATGATGATGATAAAGGTTCTGGAGA<br>AGGAAGAGGATCATTATTAACATGTGGAGATGTAGAAGAAAATCCAGGACCAGAATTCATGC<br>ATGGATCTTTAAATTGTATAGTAGCAGTTAGTCAAAATATGGGAATAGGAAAAAATGGAGATT<br>ATCCTTGGCCACCATTAAGAAATGAATTTAGATATTTTCAAAGAATGACAACAACAAGTAGTGT<br>AGAAGGAAAACAAAATTTAGTTATAATGGGAAAAAAAAACATGGTTTAGTATACCAGAAAAAAA<br>TAGACCTTTAAAAGGAAGAATAAATTTAGTATTAAGTAGAGAATTAAAAGAACCACCACAAGG<br>TGCTCATTTTTTAAGTAGATCATTAGATGATGCTTTAAAATTAACAGAACAACCAGAATTAGCA<br>AATAAAGTAGATATGGTTTGGATAGTTGGAGGAAGTTCAGTTTATAAAGAAGCAATGAATCAT<br>CCAGGACATTTAAAATTATTTGTTACTAGAATTATGCAAGATTTTGAAAGTGATACATTTTTTCC<br>AGAAATAGATTTAGAAAAATATAAATTATTACCAGAATATCCTGGTGTATTAAGTGATGTTCAA<br>GAAGAAAAAGGAATTAAATATAAATTTGAAGTATATGAAAAAAATGATTAAGTCGACTATTCC<br>TTTTTCTTATTTATATATTCATACCAATTTATTGTCTTTATAAGCGTAAAATTGGGTATATTGTGT<br>GCATATTTTTTTTTGTACATGTACATGCATGTAAATAGCTAAAATTATGAACATATTTTATTTTTT<br>GTTCAGAAAAAAAAAAACTTTACACACATAAAATGGCTATTCAGACTAGCCATATTTTATATAA<br>ACTAAACCCTATAAATTTATTAACAAATTCAAAATTTAGATTATTGTGCATTTTAATATGTTTGAA<br>AAAGTAAGGCAATATTATTATCATCATCTTTACTGTTGTAATTTCTTTGTCTCTTCAATGATTCAT<br>AAATAGTTAGACTTAATTTTTAAAATGCTTATAATATGATTAGCATAGTTAAATAAAAAAAGTT<br>GAAAAAAAAAAAAAAAAAACATATAAACACAAATGATTTTTTTTCTTCAATTTCTGTATCATATA<br>CCAATAACATTAACAGGTGAATGAGAAGGTTAAGCAATAGAAAGTGATAAAAAAAAATTGTAA<br>AAAAAATGGAGGAAAAAATTAAATTATGGCATAACTTCGTATAGCATACATTATACGAAGTTAT<br>CCTAGG |
| PfUCH-F | GATGCGGCCGCGGTACAATATGGACAAGCACCATTTG |
| PfUCH-R | GTATGGTACCTGCTATTTTCATTCCCATTTGAGC |
| TgVMP1 SgDNA1 | GCCTTCATCTGCAAATGCGTGGTTTTAGAGCTAGAAATAGC |

| | |
|---|---|
| SgDNA-RP | AACTTGACATCCCCATTTAC |
| TgVMP1-Fkd2 | CATGCTCGCCCTTCACTGACAAAAAACGAAACAAAGTTCACAAGTCGTCCGCTAGCA<br>AGGGCTCGGGC |
| TgVMP1-RP | ATGTATGCACGCGTATGTGCCTCTGCGAACAGTTATAGACAGAACACTACCTATAGG<br>GCGAATTGGAGCTCC |
| TgVMP1-5con | GACGCTTCTCCGGTCACCCG |
| TgVMP1-3con | GTTGTACCGGGAAGCAGCAG |
| TGVP1-5con | GATGCCGTACGCGTTTAG |
| TGVP1-3con | GTGACGACTGCATGATGTAC |
| TgTub-F | GCCGCTTAATTAACATGCATGTCCCGCGTTCGTG |
| TgTub-PfVMP1-Rrec | CTTCTTGGCCTCATAAGAATCCATTCGCTAAAAGGGAATTCAAGAAAAAATGCC |
| PfVMP1-Frec | AGCGAATGGATTCTTATGAGGCCAAGAAG |
| PfVMP1-R | TACGATCCTAGGGAATTCCCGCTTCTTGATCTTCCGAGGGGTC |
| BipSig-F | TCGGATATCAGCGAATGACGGCTGCCAAAAAACTCAG |
| BipSig-GFP-Rrec | GTGCCGCAGCCGCTGCGGCGCCATCACTGGCAGCGACAGGTCGG |
| GFP-Bip-Frec | GGCGCCGCAGCGGCTGCGGCACATAAAGGAGAAGAACTTTTCACTG |
| GFP-HDEL-R | ATCCCCTAGGCTACAACTCGTCGTGTTTGTATAGTTCATCCATGCCAT |
| PfVMP1syn (codon optimized for *Homo sapiens*, coding region is highlighted) | ATGGATTCTTATGAGGCCAAGAAGAAAGAGCTGTACCTGCGGCGGAAGGACATCAA<br>CCTGTACTATCACCCCATCAAGACCATCAAGCTGTTCTGCCTGCAACTGCGGAACAT<br>CATCGTGCAGACCTACCAGAAGAACAAGAAGTACAACAAGATCCTGATTCTGGCCCT<br>GCTGATCATCCTGATCCTGTTCAAGATCCGGTATAAGTACGAGCACCTGAACAACTT<br>CATCATCTACATCGAAGTGACCGTGTGGTGGCTGAGCCTGGGCATCCTGTCTAGCAT<br>CGGACTCGGCTGTGGAATGCACAGCGGCGTGCTGTTTCTGTTCCCTCACATCTACAGC<br>ATCTGCAGCACCAGCGAGTACTGCAACAGCCTGAACTTCGACAGCCGGATCAACATG<br>TGGTCCAGCGTGCTGAGCAGCGGCAACTACTTTGAGTGCCTGGGCACCAACGACGAG<br>GACATCACCTTCAGCAGACTGTTCTTCAAGATCTACCCCTACTGCCTGATCTGGGGCA |

| | |
|---|---|
| | TCGGAACAGCTCTGGGAGAGCTGCCTCCTTACCTGACCAGCTACTACGCCGCCAAGA<br>GCAAGCTGAGCGACGAGGAATTCGAAGAGTTCCAGAAGGATATCAAAGAGGGGAAG<br>AAGAATATCATCACCGCCATGAAGATCTGGATGCTGGACTTCATCAAGAAGTATGGC<br>AGCATCTCCGTGTTTCTGCTGAGCTGCTGGCCCAACGTGATGTTCGATCTGTGCGGAA<br>TCTGCTGCGGCCACTTCCTGATGCCTTTCCAGAACTTCTTCATCCCTCTGGTGCTGGG<br>CAAAGCCATCGTCAAGACAATTATCCAGTCCTTCTTCCTGATCTTCGTGTTCAGCAAC<br>AACTACAAAGAACTGCAGCTGAAGATCATCCAGAAGTGCCTGTCCATCCTGCCTCTG<br>CACCTGTTCAGCGAGAACCTGTCCGACGAGAGCTTCCTGGAAAGCTACATCGACGAG<br>AAGATCAGCATCCTGAAGCACGGCAAGAAGTCCTCCAGCAAACTGAACTTCATGTTC<br>CTCTTCAATATCTTCTTCTGCGTGCTGCTGATCGTGTTCTTCATCAGCTGCATCAACCA<br>GATCGCCCAGAAATACCAGAAAAACGTGGACGAGGAAGAACTGGAAGGCCTGCTGA<br>AGTCCAAGAACGAGATGACCCCTCGGAAGATCAAGAAG |
| HsVMP1syn (codon optimized for *Toxoplasma gondii*, coding region is highlighted) | GATATCATGGCAGAGAATGGCAAAAATTGCGACCAAAGGCGTGTGGCCATGAATAA<br>AGAACATCACAATGGAAACTTCACAGATCCATCCTCGGTCAACGAAAAAAAACGTC<br>GGGAGAGGGAAGAACGACAGAACATCGTTTTGTGGAGGCAACCACTTATCACTCTG<br>CAATATTTCAGTCTCGAAATTCTGGTGATCCTTAAGGAGTGGACAAGTAAGTTGTGG<br>CATCGCCAATCTATCGTCGTCAGTTTTCTTCTGCTCCTTGCTGTCCTGATTGCCACCTA<br>TTACGTCGAAGGTGTTCACCAACAATACGTCCAAAGAATCGAGAAACAGTTTCTTCT<br>TTATGCTTATTGGATTGGCCTTGGCATTCTTTCTAGTGTTGGCCTTGGGACTGGCCTCC<br>ACACGTTTTTGTTGTATCTTGGCCCGCACATTGCTAGCGTTACCTTGGCTGCATACGA<br>GTGCAATTCGGTCAATTTCCCTGAGCCTCCTTACCCAGACCAAATTATTTGCCCTGAT<br>GAGGAAGGCACTGAAGGTACTATTTCTCTGTGGTCTATCATTAGTAAAGTCAGGATT<br>GAAGCATGCATGTGGGGGATCGGGACTGCAATCGGAGAATTGCCCCCGTATTTCATG<br>GCAAGAGCTGCACGACTTAGTGGAGCGGAGCCTGACGATGAGGAATACCAGGAATT<br>TGAGGAGATGCTTGAACACGCGGAAAGTGCCCAAGATTTCGCCTCTCGAGCGAAACT<br>CGCGGTTCAAAAGCTTGTGCAAAAGGTGGGATTTTTCGGCATCCTCGCATGTGCGTC<br>CATCCCTAACCCACTCTTTGATTTGGCCGGGATTACTTGTGGACACTTTCTTGTTCCCT<br>TTTGGACCTTTTTTGGTGCTACCTTGATCGGGAAAGCGATCATCAAAATGCATATCCA<br>AAAAATCTTTGTCATCATTACATTCTCCAAACATATCGTTGAGCAGATGGTGGCCTTC<br>ATCGGAGCCGTTCCTGGAATCGGTCCATCTCTGCAAAAACCCTTCCAGGAGTACCTC<br>GAGGCTCAAAGACAAAAATTGCATCATAAAAGCGAAATGGGCACCCCACAGGGCGA<br>GAATTGGCTGTCGTGGATGTTCGAGAAGTTGGTGGTCGTCATGGTGTGCTATTTCATT<br>CTTTCTATTATTAACTCCATGGCGCAGAGCTACGCTAAACGTATCCAACAGCGCTTGA<br>ACTCTGAGGAAAAAACTAAGCGGGAATTCCCTAGG |
