## Supplementary Figures for "An ER-resident lipid scramblase is crucial for biogenesis and function of apicomplexan parasite secretory organelles"

### Supplementary figures and legends

#### List of figures

Figure S1. PfVMP1 and TgVMP1 share the domain architecture of DedA superfamily.

Figure S2. Generation of PfVMP1GFP$_{KI}$ and RHTIR1-VMP1AID parasites.

Figure S3. Dual-knock-down strategy for PfVMP1.

Figure S4. Auxin-inducible knock-down strategy for PfVMP1.

Figure S5. TgVMP1 depletion did not affect SAG1 and IMC1 transport.

Figure S6. Conoid extrusion is not affected upon TgVMP1 depletion.

Figure S7. TgVMP1 depletion did not affect microneme organization.

Figure S8. TgVMP1 is crucial for IVN formation and dense granule biogenesis.

Figure S9. TgVMP1 is crucial for ER organization.

Figure S10. TgVMP1 immunoprecipitation.

Figure S11. TgVMP1 and PfVMP1 are lipid scramblases, and restoration of the ER-localized scramblase activity rescued TgVMP1-depleted parasites.

Figure S12. TgVMP1 depletion elevated calcium level.

Figure S13. A model for the functions of TgVMP1.

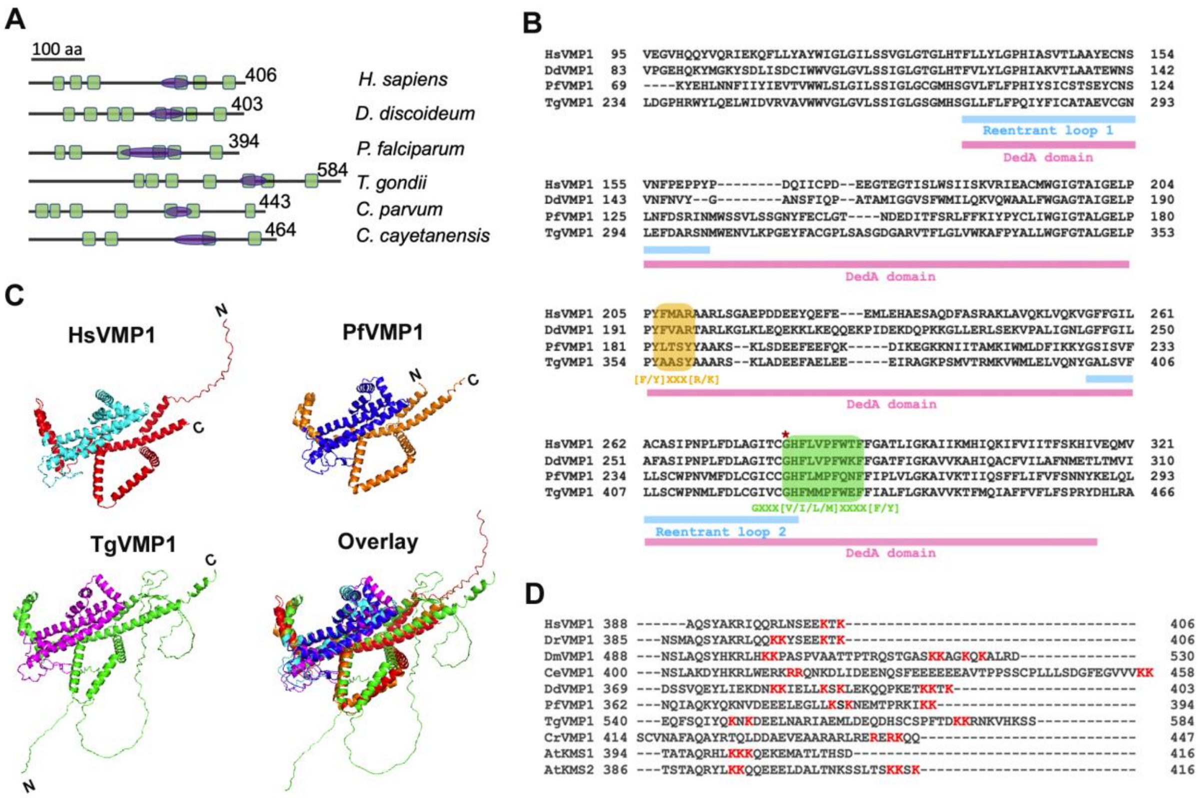


**Figure S1. PfVMP1 and TgVMP1 share the domain architecture of DedA superfamily. A.** The schematic shows domain architectures of the characterized VMP1 proteins (*H. sapiens and D. discoideum*) and their putative apicomplexan homologs (*P. falciparum*, *T. gondii*, *C. parvum*, and *C. cayetanensis*). Transmembrane region (green), DedA domain (purple), and the number of amino acid residues in each protein are indicated. **B.** The amino acid sequence alignment of VMP1 proteins from *H. sapiens* (HsVMP1), *D. discoideum* (DdVMP1), *P. falciparum* (PfVMP1), and *T. gondii* (TgVMP1) shows DedA domain (pink), re-entrant loops (blue), conserved motifs (highlighted in yellow and green), and the conserved "Gly" residue (*). The numbers indicate the position of amino acid residues in the respective full-length proteins. **C.** Shown are the AlphaFold structures along with the DedA domain (HsVMP1: cyan; PfVMP1: blue; TgVMP1: magenta) and overlay of the structures of HsVMP1, PfVMP1 and TgVMP1. **D.** The amino acid sequences of the predicted cytoplasmic tails of VMP1 homologs from *H. sapiens* (HsVMP1), *D. rerio* (DrVMP1), *D. melanogaster* (DmVMP1), *C. elegans* (CeVMP1), *D. discoideum* (DdVMP1), *P. falciparum* (PfVMP1), *T. gondii* (TgVMP1), *C. reinhardtii* (CrVMP1), and *A. thaliana* (AtKMS1 and AtKMS2) were aligned. Shown are the "di-lysine" motifs or their variants (red font).

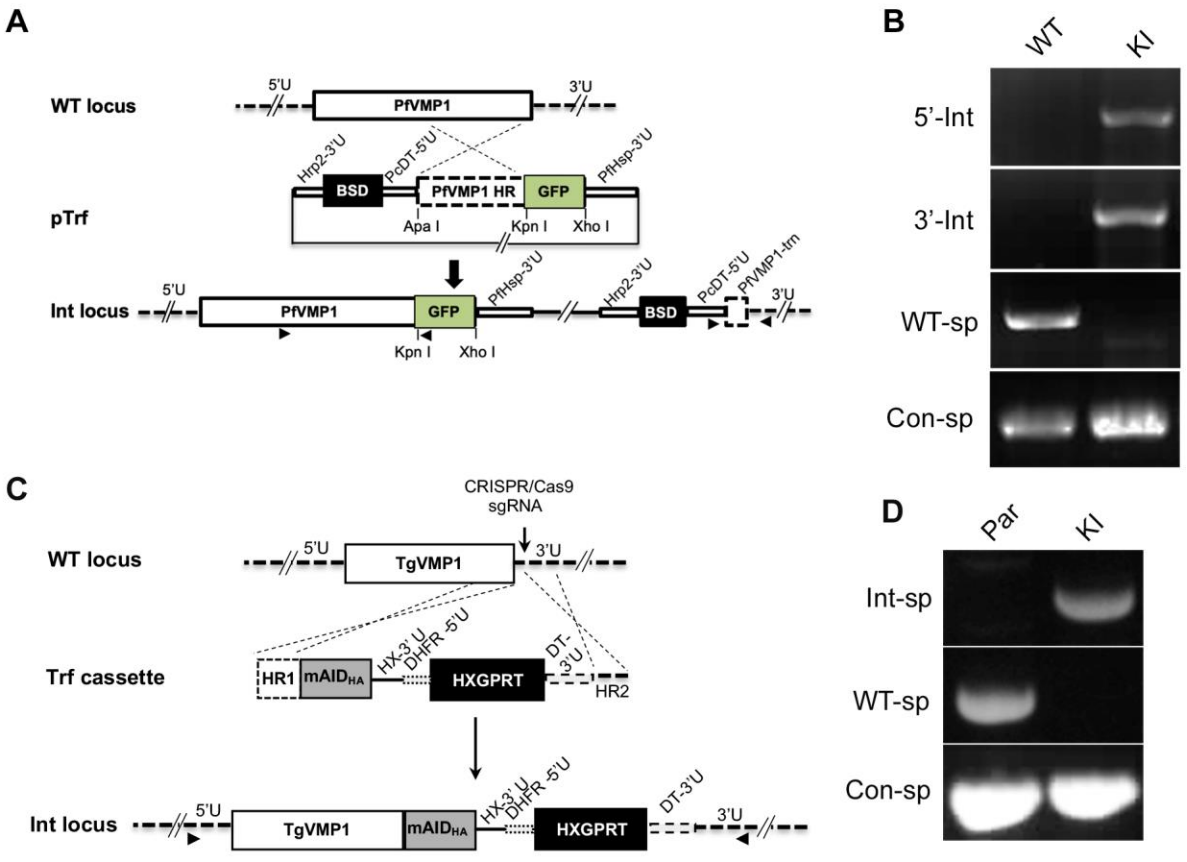


**Figure S2. Generation of PfVMP1GFP$_{KI}$ and RHTIR1-VMP1AID parasites. A.** The schematic shows insertion of the transfection plasmid (pTrf) into the wild-type PfVMP1 locus (WT locus), generating the integration locus (Int locus). The coding regions (PfVMP1, PfVMP1 homology region (PfVMP1 HR), GFP, BSD), untranslated regions (5'U and 3'U of PfVMP1, PcDT-5'U, Hrp2-3'U, PfHsp-3'U), restriction endonuclease sites (vertical lines), and primer location (arrowheads) are indicated. The truncated PfVMP1 coding region (PfVMP1-trn) generated upon single homologous recombination is shown by dotted rectangular box. **B.** The genomic DNA of wild-type (WT) and PfVMP1GFP$_{KI}$ (KI) parasites were used in PCR using locus-specific primers. The ethidium bromide-stained agarose gel image shows PCR products amplified using primers specific for 5'-integration locus (5'-Int), 3'-integration locus (3'-Int), wild-type specific (WT-sp), and positive control locus (Con-sp). The absence of wild-type PfVMP1 and the presence of integration loci confirmed the desired genetic modification. **C.** The coding sequence for mAID-3×HA (mAID$_{HA}$) was inserted in-frame into the 3'-coding sequence of TgVMP1 using CRISPR/Cas9 strategy. The schematic shows wild-type TgVMP1 locus (WT locus), transfection cassette (Trf cassette), and integration locus (Int locus). Different regions of the WT locus (5'U and 3'U), Trf cassette (homology region 1 (HR1), mAID$_{HA}$, HXGPRT-3'UTR (HX-3'U), selection cassette (DHFR-5'U-HXGPRT-DT-3'U), homology region 2 (HR2), integration locus (Int locus), and primer location (arrowhead) are indicated. **D.** The genomic DNA of parental RHTIR1 (Par) and RHTIR1-VMP1AID (KI) parasites were assessed for modification of the target locus by PCR using locus-specific primers. The ethidium bromide-stained agarose gel image shows PCR products amplified using primers specific for integration locus (Int-sp), wild-type locus (WT-sp), and positive control locus (Con-sp). The presence of integration locus and the absence of wild-type locus in RHTIR1-VMP1AID parasites confirmed the desired genetic modification.

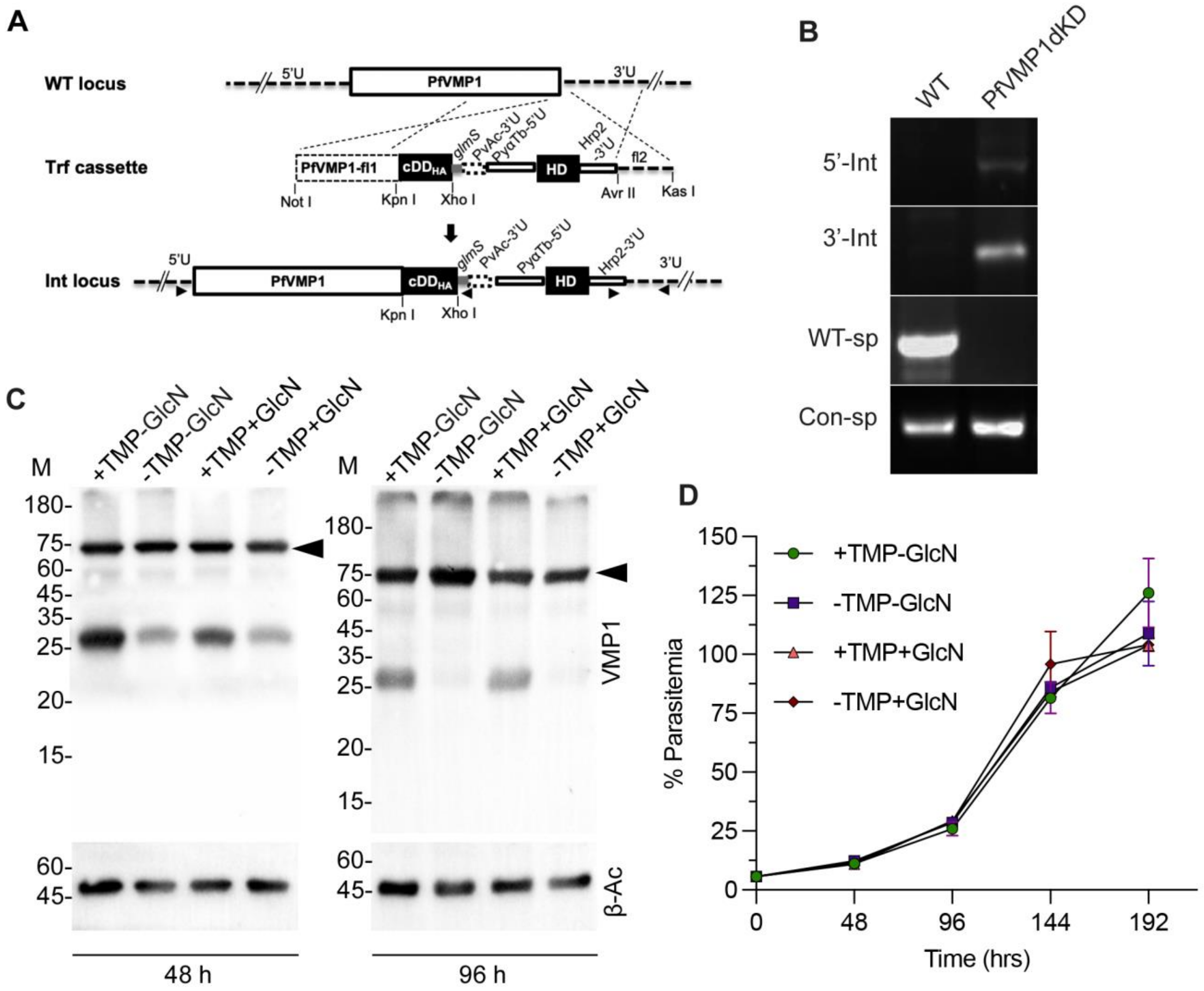


**Figure S3. Dual-knock-down strategy for PfVMP1. A.** The schematic shows replacement of wild-type PfVMP1 coding region (WT locus) by the transfection cassette (Trf cassette) via double-crossover, generating the integration locus (Int locus). The untranslated regions (5'U, 3'U), coding regions (PfVMP1, PfVMP1-flank1 (PfVMP1-fl1), cDD$_{HA}$, human DHFR (HD)), flank-2 (fl2), regulatory regions (*glmS*, PvAc-3'U, PyαTb-5'U, Hrp2-3'U), restriction enzyme sites (vertical lines) and primer position (arrowheads) are labeled. **B.** Wild-type (WT) and PfVMP1dKD genomic DNAs were used as templates in PCR using locus specific primers (5' integration (5'-Int), 3' integration (3'-Int), wild-type (WT-sp), and control (Con-sp)). The ethidium bromide-stained agarose gel shows the indicated PCR products. **C.** PfVMP1dKD parasites were grown under normal (+TMP-GlcN), single knock-down (-TMP-GlcN, +TMP+GlcN) or double knock-down (-TMP+GlcN) conditions for 48 or 96 hrs. Lysates of these parasites were assessed for PfVMP1-cDD$_{HA}$ levels (VMP1) and β-actin (β-Ac) as a loading control by western blotting. The blots show no change in PfVMP1-cDD$_{HA}$ levels (arrowhead) irrespective of the culture condition. The protein size markers (M) are in kDa. **D.** Equal number of PfVMP1dKD parasites were grown under normal (+TMP-GlcN), single knock-down (-TMP-GcN, +TMP+GlcN) or double knock-down (-TMP+GlcN) conditions for 4 consecutive cycles (192 hrs), and the number of parasite-infected RBCs was determined at the end of each cycle. The plot shows % parasitemia over the indicated growth period (Time (hrs)). The data is mean with SD error bar from two independent experiments.

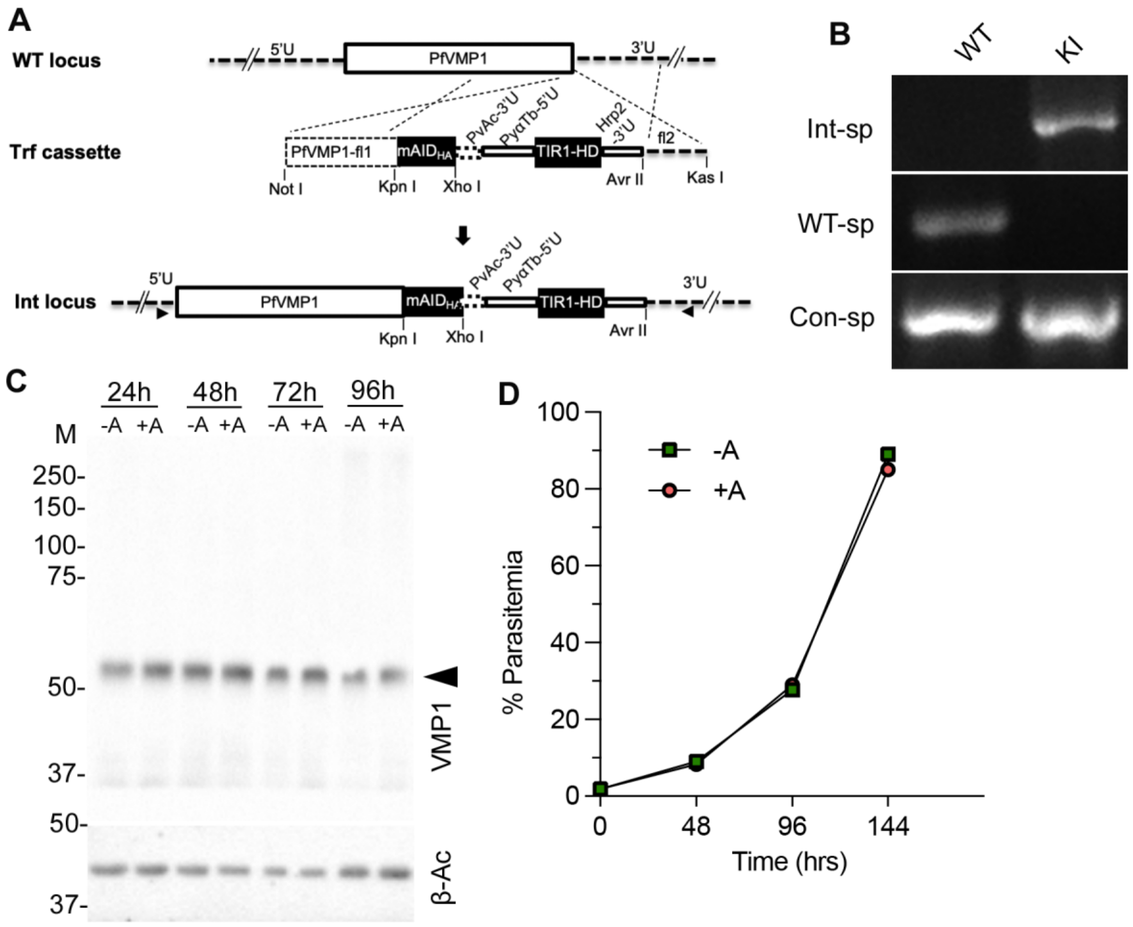


**Figure S4. Auxin-inducible knock-down strategy for PfVMP1. A.** The schematic shows replacement of wild-type PfVMP1 coding sequence (WT locus) by the linear transfection cassette (Trf cassette) via double homologous crossover, generating the integration locus (Int locus). The untranslated regions (5'U, 3'U), coding regions (PfVMP1, PfVMP1-fl1, mAID$_{HA}$, human DHFR (HD)), flank-2 (fl2), regulatory regions (PvAc-3'U, PcDT-5'U, Hrp2-3'U), restriction enzyme sites (vertical lines) and primer positions (horizontal arrowheads) are labeled. **B.** The genomic DNAs of wild-type (WT) and PfVMP1AID (KI) parasites were used in PCR using primers specific for integration (Int-sp), wild-type (WT-sp), and control (Con-sp) loci. The ethidium bromide-stained agarose gel shows the indicated PCR products in WT and KI lanes. **C.** PfVMP1AID parasites were cultured with (+A) or without (-A) 5Ph-IAA for two consecutive cycles, harvested at the indicated time points, and the lysates were evaluated for the levels of PfVMP1AID$_{HA}$ and β-actin as a loading control by western blotting. The western blot shows the levels of PfVMP1AID$_{HA}$ (VMP1, arrowhead) and β-actin (β-Ac) at the indicated time points. **D.** Three PfVMP1AID clonal lines were grown with (+A) or without (-A) 5Ph-IAA for three consecutive cycles (144 hrs), the percentage of parasite-infected RBCs was determined at the end of each cycle, and plotted as % parasitemia over 144 hrs. The data is a representative for the three clones.

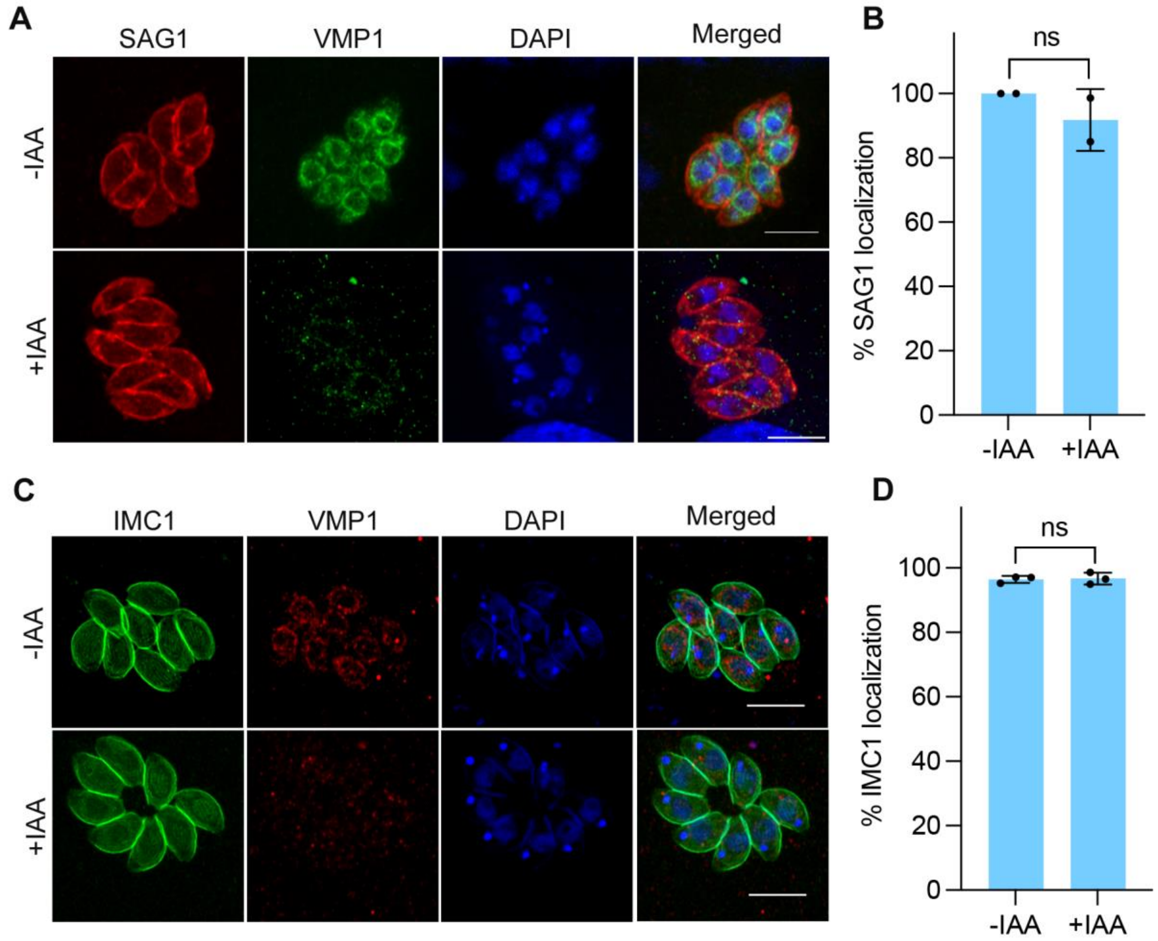


**Figure S5. TgVMP1 depletion did not affect SAG1 and IMC1 transport.** -IAA and +IAA intracellular RHTIR1-VMP1AID parasites were assessed for integrity of the plasma membrane and IMC by IFA using anti-SAG1 and anti-IMC1 antibodies, respectively. The representative Airyscan micrographs show localization of SAG1 (**A**) and IMC1 (**C**). The panels are for SAG1 or IMC1, TgVMP1AID$_{HA}$ (VMP1), nucleus (DAPI), and overlap of the three images (Merged). Scale bar = 5 µm. **B.** The number of parasites showing uniform peripheral localization of SAG1 in "A" was determined and plotted as a percentage of the total parasites observed (% SAG1 localization) on y-axis for the indicated parasites on x-axis. Each data is from 2 independent experiments (-IAA, n=104 cells; +IAA, 112 cells). **D.** The number of parasites showing uniform peripheral IMC1 localization in "C" was determined and plotted as a percentage of the total parasites observed (% IMC1 localization) on y-axis for the indicated parasites on x-axis. Each data is based on at least 100 parasites from each of the 3 independent experiments. The significance of difference between two data sets is indicated (ns: not significant).

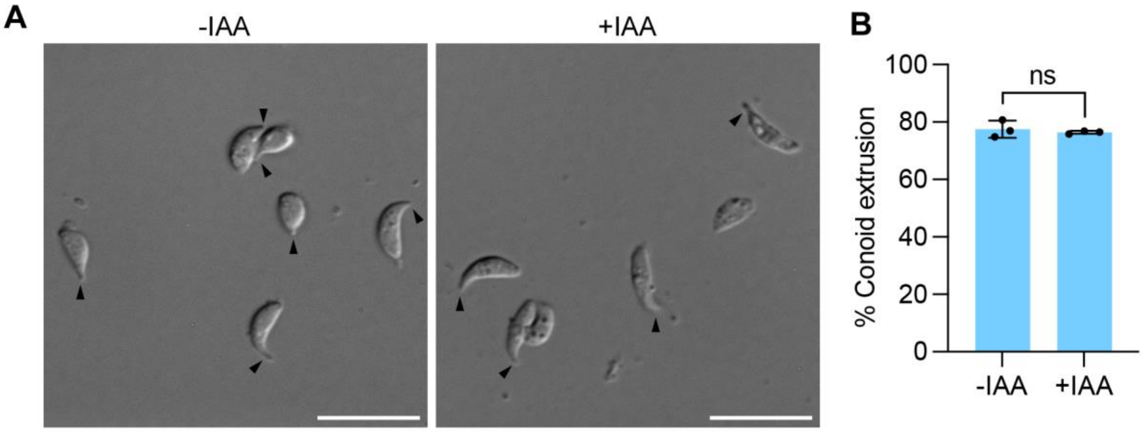


**Figure S6. Conoid extrusion is not affected upon TgVMP1 depletion.** Purified +IAA and –IAA RHTIR1-VMP1AID tachyzoites were treated with the calcium-ionophore A23187, and assessed for conoid extrusion. **A.** Shown are the representative DIC micrographs of the parasites with extruded conoids (arrowheads). Scale bar = 10 µm. **B.** The number of parasites exhibiting conoid extrusion in "A" was counted, and plotted as a percentage of the total number parasites observed (% Conoid extrusion) on y-axis against the indicated parasites on x-axis. Each data is mean with SD error bar based on at least 100 observations in each of the 3 independent experiments. The significance of difference between two data sets is indicated (ns: not significant).

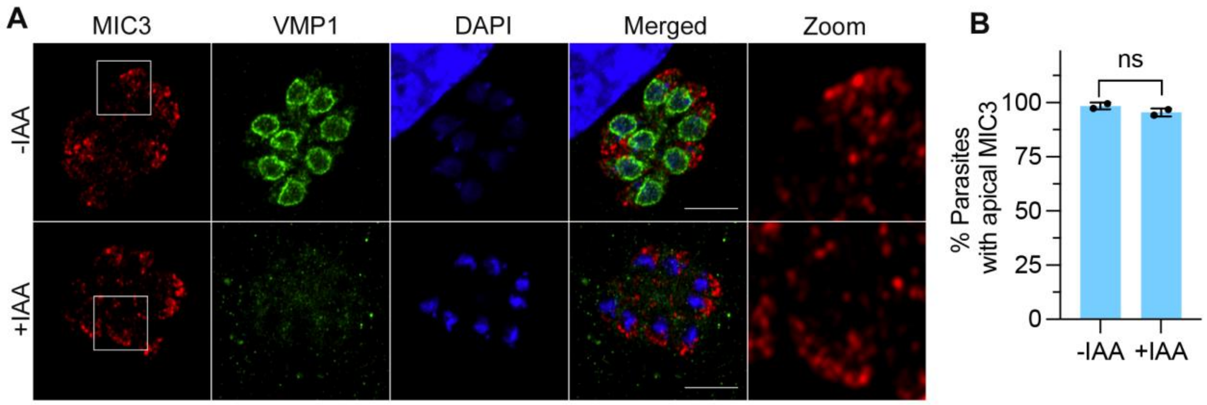


**Figure S7. TgVMP1 depletion did not affect microneme organization.** Intracellular -IAA and +IAA RHTIR1-VMP1AID parasites were evaluated for microneme organization by labelling the parasite with antibodies to MIC3. **A.** The representative Airyscan micrographs show localization of MIC3, TgVMP1AID$_{HA}$ (VMP1), nucleus (DAPI), and overlap of the three images (Merged). Scale bar = 10 µm, and the boxed area in MIC3 panel is zoomed-in (Zoom). Note the similar apical MIC3 localization in –IAA and +IAA parasites. **B.** The number of parasites with apical MIC3 localization was determined, and shown as a percentage of the total number of parasites observed (% Parasites with apical MIC3) on y-axis against the indicated parasites on x-axis. Each data is mean with SD error bar based on at least 100 observations from each of the 2 independent experiments. The significance of difference between two groups is shown (ns: not significant).

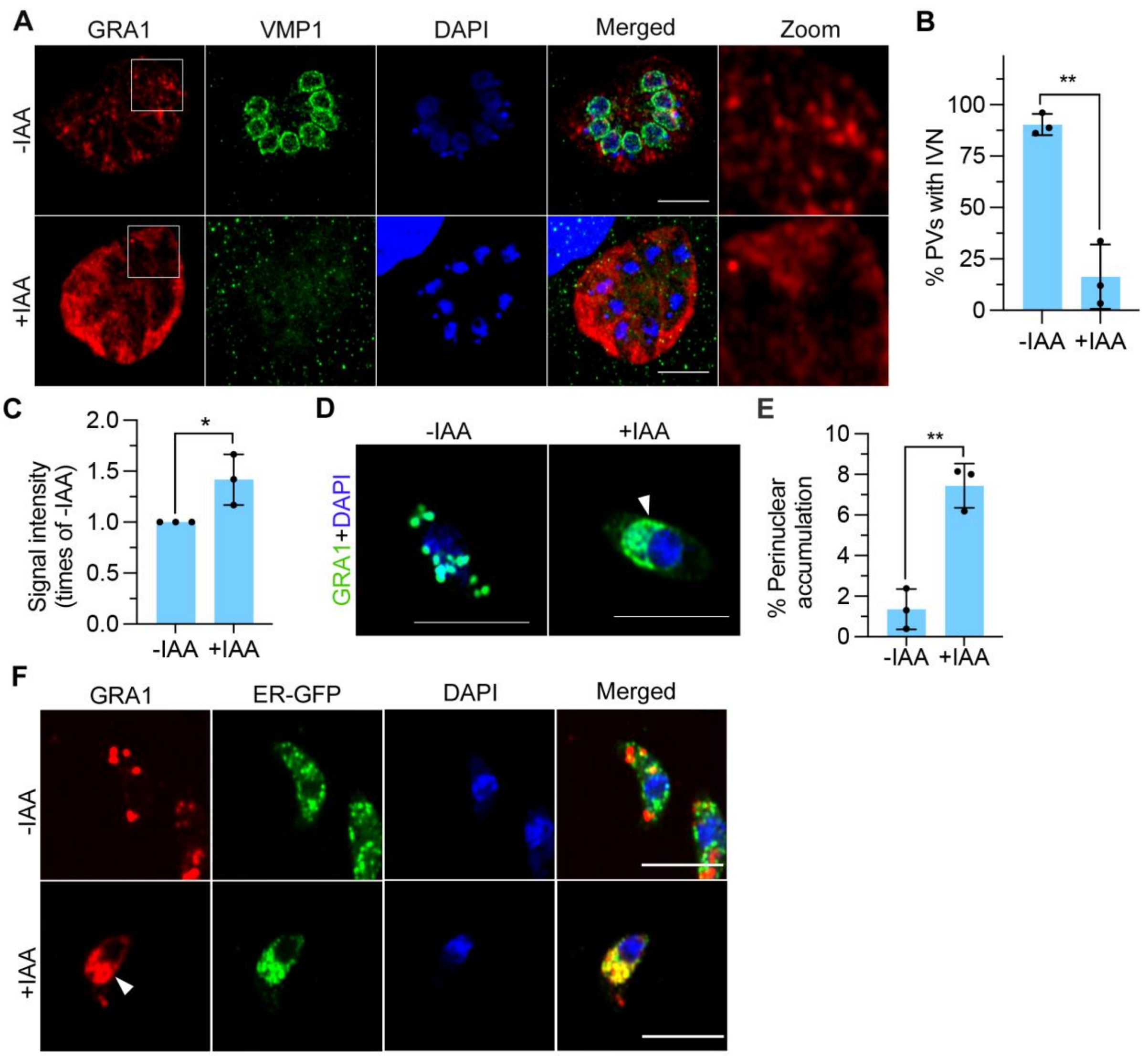


**Figure S8. TgVMP1 is crucial for IVN formation and dense granule biogenesis.** -IAA and +IAA RHTIR1-VMP1AID parasites were evaluated for IVN and dense granule biogenesis using the dense granule protein GRA1 as a marker. **A.** Intracellular –IAA and +IAA parasites were processed for IFA using antibodies to GRA1, and observed for IVN. The representative Airyscan micrographs show localization of GRA1, TgVMP1AID$_{HA}$ (VMP1), nucleus (DAPI), and overlap of all the three images (Merged). Scale bar is 10 µm, and the boxed area in GRA1 panel is zoomed-in (Zoom). Note the GRA1-labeled distinct tubulo-vesicular IVN in the PV of –IAA parasites, whereas GRA1 signal is diffuse or associated with fragmented structures in the PV of +IAA parasites. **B.** The number of PVs showing IVN in "A" was determined, and plotted as a percentage of the total PVs observed (% PVs with IVN) on y-axis against the indicated parasites on x-axis. Each data is mean with SD error bar based on ≥50 PVs from each of the 3 independent experiments. **C.** Purified -IAA and +IAA RHTIR1-VMP1AID parasites were assessed for GRA1 localization by IFA, and assessed for the signal intensity of GRA1. The signal intensity of GRA1 in each of the -IAA and +IAA parasites was measured, and the mean signal intensity of GRA1 in +IAA parasites was plotted as times of the signal intensity of -IAA parasites (Signal intensity (times of –IAA)) on y-axis for the indicated parasites on x-axis. Each data represents mean signal intensity with SD error bar based on multiple parasites (-IAA, n=416; +IAA, n=651) from three independent experiments. **D.** The representative Airyscan micrographs show merged image of GRA1 and DAPI panels for –IAA and +IAA parasites. Scale bar is 5 µm. The arrow head indicates perinuclear accumulation of GRA1 in +IAA parasites, which is absent in -IAA parasites. **E.** The number of parasites showing perinuclear accumulation of GRA1 in "D" was plotted as a percentage of the total number of parasites observed (% Perinuclear accumulation) on y-axis for the indicated parasites on x-axis. Each data is based on multiple parasites (-IAA, n=416; +IAA, n=651) from 3 independent experiments. **F.** The representative Airyscan micrographs show localization of GRA1, ER-GFP, DAPI, and overlap of the three images (Merged) in the indicated parasites. Scale bar is 5 µm. Note the prominent co-localization of GRA1 and ER-GFP in the perinuclear region in +IAA parasites, which is nearly absent in -IAA parasites. The significance of difference between the data sets is indicated with p-value (*= $P<0.05$; **= $P<0.01$).

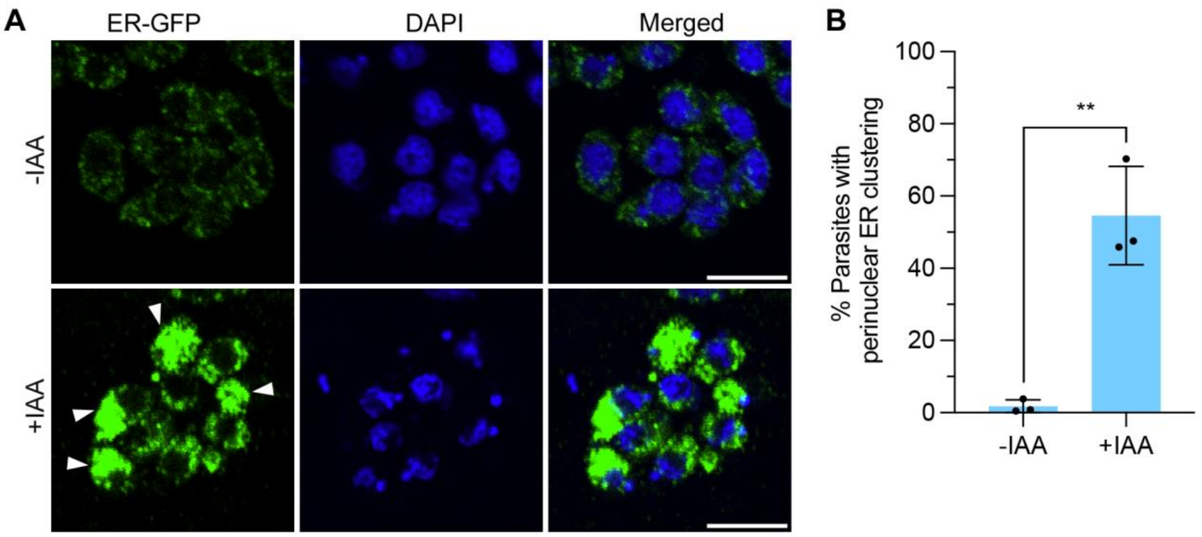


**Figure S9. TgVMP1 is crucial for ER organization.** Intracellular -IAA and +IAA RHTIR-VMP1AID::BiP-GFP-HDEL parasites were evaluated for localization of ER-GFP. **A.** The representative confocal images are for ER-GFP, nucleus (DAPI), and overlap of the two panels (Merged). Note the prominent perinuclear clustering of ER-GFP (arrowheads) in +IAA parasites as opposed to uniform ER-GFP signal in -IAA parasites. Scale bar = 5 µm. **B**. The number of parasites with perinuclear clustering of ER-GFP in "A" was determined, and plotted as a percentage of the total number of parasites observed (% Parasites with perinuclear ER clustering) on y-axis for the indicated parasites on x-axis. Each data is mean with SD error bar based on multiple parasites (-IAA, n = 2058; +IAA, n = 1136) from 3 independent experiments. The significance of difference between the data sets is indicated with p-value (**= P<0.01)

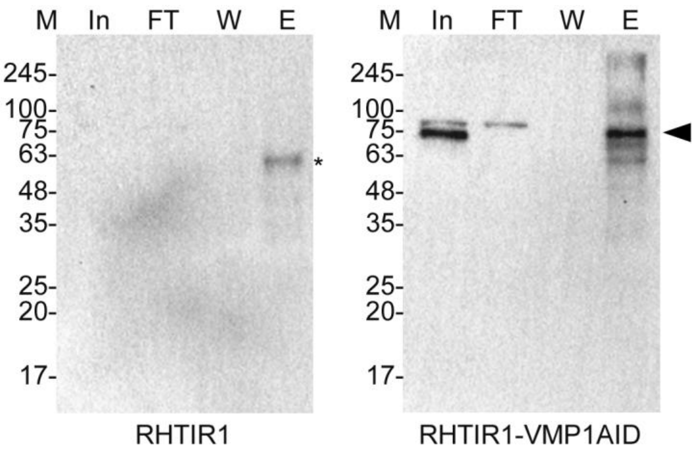


**Figure S10. TgVMP1 immunoprecipitation.** Purified –IAA RHTIR1 and -IAA RHTIR1-VMP1AID tachyzoites were processed for immunoprecipitation using rabbit anti-HA antibodies, and aliquots of the input (In), flow through (FT), wash (W), and eluate (E) samples were processed for western blotting using mouse anti-HA antibodies. The blots show successful immunoprecipitation of $TgVMP1AID_{HA}$ (arrowhead) from RHTIR1-VMP1AID parasites. The eluate sample in RHTIR1 showed a band of about 63 kDa (*), which may be a cross-reactive protein. The protein size markers (M) are in kDa.

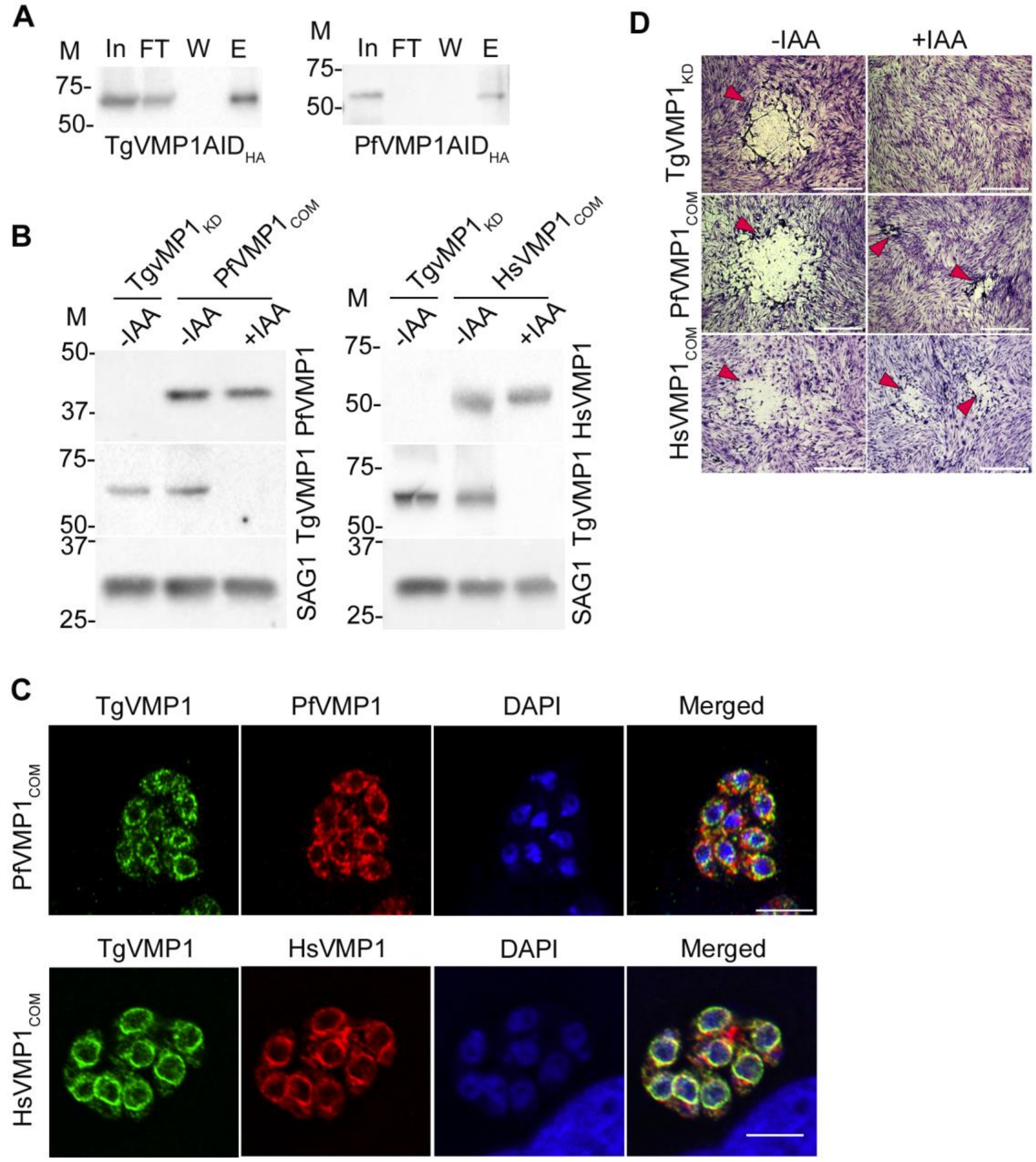


**Figure S11. TgVMP1 and PfVMP1 are phospholipid scramblases, and restoration of the ER-localized scramblase activity rescued TgVMP1-depleted parasites. A.** RHTIR1-VMP1AID and PfVMP1AID parasites were cultured without IAA and 5Ph-IAA, respectively. The parasites were purified and processed for immunoprecipitation using HA-Trap antibodies. Aliquots of the input (In), flow through (FT), wash (W), and eluate (E) samples were processed for western blotting using rabbit anti-HA antibodies. The blots show immunoprecipitation of TgVMP1AID$_{HA}$ and PfVMP1AID$_{HA}$. The protein size markers (M) are in kDa. **B.** RHTIR1-VMP1AID (TgVMP1$_{KD}$), RHTIR1-VMP1AID/PfVMP1 (PfVMP1$_{COM}$), and RHTIR1-VMP1AID/HsVMP1 (HsVMP1$_{COM}$) tachyzoites were cultured with (+IAA) or without (-IAA) IAA. The parasite lysates were evaluated for the presence of TgVMP1AID$_{HA}$ (TgVMP1), PfVMP1$_{Myc}$ (PfVMP1), HsVMP1$_{Myc}$ (HsVMP1), and SAG1 as a loading control. The arrowhead indicates the target protein band and the protein size markers (M) are in kDa. **C.** Intracellular –IAA PfVMP1$_{COM}$ and HsVMP1$_{COM}$ tachyzoites were checked for co-localization of PfVMP1 or HsVMP1 with TgVMP1 by IFA. The Airyscan micrographs show signal for TgVMP1, PfVMP1 or HsVMP1, nuclear stain (DAPI), and overlap of the three images (Merged) for the respective parasites. The scale bar = 5 µm. Note the co-localization of PfVMP1 or HsVMP1 with TgVMP1 (Pearson's correlation coefficient: HsVMP1 = 0.78±0.09, n = 24 images; PfVMP1 = 0.54±0.12, n = 33). **D**. $TgVMP1_{KD}$, $PfVMP1_{COM}$, and $HsVMP1_{COM}$ tachyzoites were grown without (-IAA) or with (+IAA) IAA for 8 days, stained with Giemsa, and observed for plaques. Note the absence of plaques in +IAA $TgVMP1_{KD}$, as compared with clear plaques (arrowhead) in +IAA $PfVMP1_{COM}$ and +IAA $HsVMP1_{COM}$ parasites. Scale bar = 1 mm.

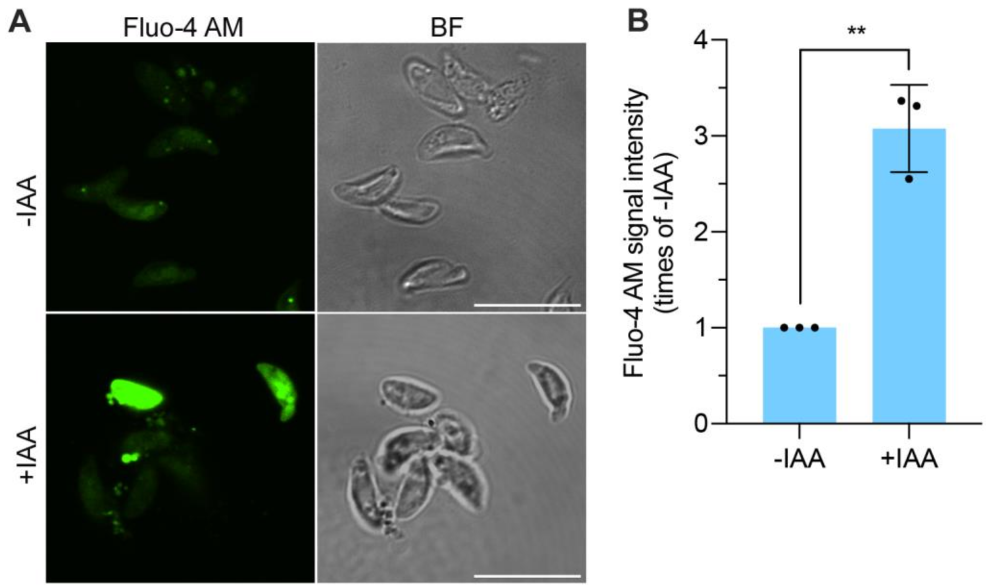


**Figure S12. TgVMP1 depletion elevated calcium level.** Purified –IAA and +IAA RHTIR1-VMP1AID tachyzoites were examined for calcium levels by staining with the calcium sensor Fluo4-AM. **A.** The confocal micrographs show calcium signal (Fluo-4 AM) and bright field (BF) images of the indicated parasites. Scale bar = 10 µm. Note the intense calcium signal in some of the +IAA parasites as compared with –IAA parasites. **B.** The calcium signal intensity was measured for each of the parasites, and the mean signal intensity for +IAA parasites was plotted as times of the signal intensity of -IAA parasites (Fluo4-AM signal intensity (times of –IAA)) on y-axis for the indicated parasites on x-axis. Each data represents mean signal intensity with SD error bar based on multiple parasites (-IAA, n = 415; +IAA, n = 356) from 3 independent experiments. The significance of difference between the data sets is indicated with p-value (**= P<0.01).

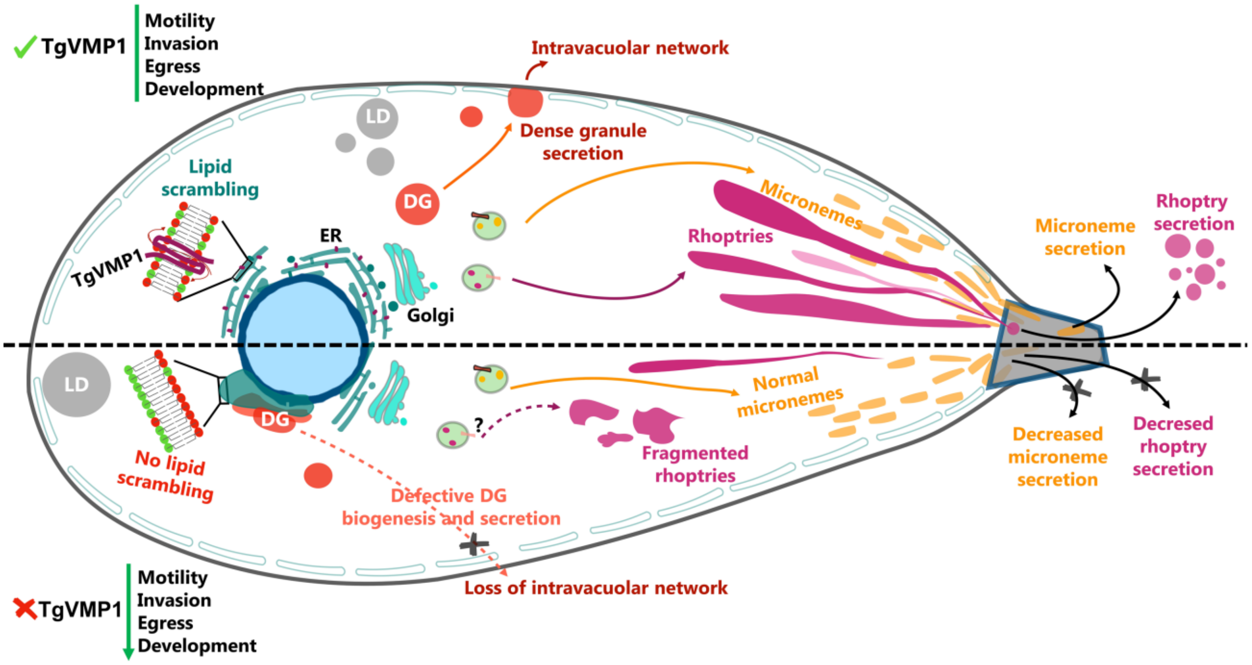


**Figure S13. A model for the functions of TgVMP1**. The schematic shows indicated cellular structures and processes in wild type (upper half) and TgVMP1-depleted tachyzoite (lower half). TgVMP1 is an ER-localized lipid scramblase, and is most likely required for translocation of lipids in ER and/or the precursor vesicles destined to the secretory organelles (micronemes, rhoptries, and dense granules). The loss of TgVMP1 impaired biogenesis of rhoptries and dense granules (DG), function of secretory organelles, lipid droplet (LD) homeostasis, and ER organization, resulting in the decreased parasite motility, host cell invasion, egress, and development.
